# A cell-and-plasma numerical model reveals hemodynamic stress and flow adaptation in zebrafish microvessels after morphological alteration

**DOI:** 10.1101/2022.12.07.519059

**Authors:** Swe Soe Maung Ye, Li-Kun Phng

## Abstract

The development of a functional cardiovascular system ensures successful embryonic development and a sustainable oxygen, nutrient and hormones delivery system for homeostasis in adulthood. While early vessels are formed by biochemical signaling and genetic programming, the onset of blood flow provides mechanical cues that participate in the vascular remodeling of the embryonic cardiovascular system. The zebrafish is a prolific animal model for studying the quantitative relationship between blood flow and vascular morphogenesis due to a combination of favorable factors including blood flow visualization in optically transparent larvae. While imaging techniques can be employed for measuring flow velocity and blood perfusion levels, hemodynamic forces such as the lumen wall shear stress (WSS) and lumen blood pressure cannot be measured directly. In this study, we have developed a cell and plasma blood transport model using CFD to understand how red blood cell (RBC) partitioning asymmetries affect WSS in a network. Furthermore, we employed the CFD model on rheological and morphological alterations of a wildtype network. We discuss how careful consideration of boundary conditions and acquisition of systemic parameters of blood flow are required to understand the intimate relationship between flow physiology and compensatory responses to hemorheological or morphological alterations.

## 1. Introduction

The cardiovascular system is essential for embryonic development, tissue growth and homeostasis in adults by providing sustainable oxygen, nutrients and hormones to tissue. It also aids in the removal of metabolic waste. Angiogenesis, which describes the process of blood vessel network expansion from preexisting vessels is a topic of active research due to the physiological importance of early stage cardiovascular system development in the survival of a growing organism. While many animal models have been employed for the purposes of such studies (Cesarovic et al., 2020; Jia et al., 2020), the zebrafish embryo has been one of the most prolific models for angiogenesis research. This can be attributed to the zebrafish’s rapid maturation time, low cost and relative ease of animal husbandry, extensive taxonomy of transgenic variants for a multitude of vascular disease representation (Hogan and Schulte-Merker, 2017) and most importantly, the relative ease of blood flow and vessel morphology quantification through non-invasive microscopic imaging.

While a functional cardiovascular system for directional convection of blood is necessary in an adult zebrafish, dysfunction of this system is not fatal for embryos in their first week of existence (Ransom et al., 1996). Indeed, in the early stages of organ and tissue development, cutaneous delivery of oxygen and nutrients from environmental sources puts less significance on the oxygen delivery function of a vascular network (Hughes and Perry, 2021). Yet, for a healthy zebrafish population, the cardiovascular system is always established with stereotyped anatomical features that is reproducible across zebrafish larvae in their first week of development. Consequently, researchers believe that the patterning cues for angiogenesis at embryonic stage are not purely dominated by genetic programming and biochemical signaling, and that mechanical forces associated with blood flow play direct roles in vascular development and remodeling. For example, Notch activation by differing blood flow levels amongst inter-connected vessels regulates endothelial cell (EC) migration involved in determining a vessel’s arterial or venous fate in a network (Geudens et al., 2019; Weijts et al., 2018). In vascular remodeling of plexus connections, vessel pruning occurs primarily in vessels that have lower blood perfusion in comparison to adjacent vessel connections (Chen et al., 2012; Kochhan et al., 2013). In contrast to pruning, vessels with low perfusion and wall shear stress (WSS) are targeted by intussusception, which expands network connections in the caudal vein plexus by splitting these vessels via trans-capillary tissue pillar formation (Karthik et al., 2018). During transcellular lumen formation of angiogenic vessels, apical membranes of ECs are exposed to the expansionary forces of blood pressure which drive the lumen invagination process via a tightly regulated process of tension-induced meta-instability in the apical membrane (Gebala et al., 2016). Blood flow has also been found to contribute to dorsal longitudinal anastomotic vessel (DLAV) morphogenesis, where blood flow abrogation has been demonstrated to inhibit anastomosis and formation of contralateral plexus connections (Zygmunt et al., 2012).

These recent studies contribute to the growing narrative that blood flow directs a multitude of cellular events that contribute to angiogenesis and vascular remodeling through nuanced and separate mechanisms. As such, the measurement of hemodynamics in zebrafish is a subdiscipline of growing interest and application within blood vascular development. With confocal fluorescence microangiography and ultramicroscopy, high resolution 3-D geometries of the blood lumen network can be obtained (An et al., 2020; Jährling et al., 2009). Likewise, cardiovascular performance parameters such as blood flow velocity and flow pulsatility can be measured by imaging techniques such as particle tracking velocimetry (Tinevez et al., 2017; Ye et al., 2022), laser-scanning velocimetry (Malone et al., 2007) and optical tweezers (Harlepp et al., 2017). However, high precision maps of the lumen WSS and distribution of the blood pressure within the network are parameters that cannot be directly measured. These hemodynamic forces that are essential to understanding EC mechanotransduction and vascular morphogenesis require mathematical models for their calculation from the experimentally obtained flow and geometry data. While there are coarse-grained techniques to assess hemodynamic forces (Ye et al., 2022), computational fluid dynamics (CFD) models subscribing to Navier Stokes prescription of flow physics has been regarded as the gold standard for mechanistic accuracy and precision. Outside of zebrafish hemodynamics, CFD models employing red blood cell (RBC) and plasma representation of blood transport dynamics has become the expected standard for model prescription in microhemodynamics (Alizadehrad et al., 2012; Doddi and Bagchi, 2009; Dupin et al., 2007; Fedosov et al., 2011; Vahidkhah et al., 2014). Conversely, many CFD studies on zebrafish hemodynamics have typically avoided multiphase representation of blood microrheology due to their focus on larger vessels and fluid chambers such as the heart (Lee et al., 2018; Salman and Yalcin, 2021; Vedula et al., 2017). Others have applied a mixed multi-scale scheme of continuum blood prescription at network level to feed boundary flow conditions into the single vessel scale model that prescribe a two-phase RBC and plasma representation of blood (Zhou et al., 2021). Some CFD studies have omitted RBCs from their blood flow model entirely (Roustaei et al., 2022) or limited the number of flowing RBCs to represent only extremely low hematocrit conditions (Djukic et al., 2016). These coarse-grained modeling approaches are undertaken possibly due to the complexities of the highly frequent inter-RBC collisions and high computational costs associated with deformable RBC simulations. However, the lack of an explicit consideration of RBCs poorly characterizes blood flow in microvascular networks since RBC dynamics directly modulate blood viscosity at micron length scales (Lanotte et al., 2016). The influence of RBCs on blood rheology is particularly important at flow-partitioning bifurcations where plasma and RBCs do not follow symmetric behavior in their flow division (Enjalbert et al., 2021; Pries et al., 1990). Furthermore, the positional asymmetry of the core RBC flow trajectory after bifurcations can heighten the lumen WSS downstream (Balogh and Bagchi, 2019; Ye et al., 2016).

Given the importance of RBC phase representation in blood microrheology, we developed an RBC and plasma CFD model for the study of hemodynamic stress distribution in developing zebrafish. In this study, we aim to demonstrate the usefulness of the CFD approach by analyzing flow and stress alteration patterns in the zebrafish trunk vascular network in response to alterations in hemorheology and network morphology. Additionally, we discuss the process of integrating flow data from experiments into the boundary conditions employed in the CFD model. Essentially, we demonstrate how careful examination and validation of CFD boundary conditions are necessary to properly understand the intimate relationship between flow physiology and compensatory responses to hemorheological or morphological alterations.

## 2. Results

Using microangiography and fluorescence confocal microscopy, we reconstructed a high resolution 3-D *in silico* model of the blood vessel lumen network geometry at the anterior half of the caudal plexus region of the zebrafish trunk at 2 days post fertilization (dpf). The geometry model preserved topological features of the physiological network such as variations in diameters along the vessel axes and irregularity of the local endothelium surface (Fig. 1A). To maintain the hematocrit level, we recycled RBCs within reservoir domains at the anterior of the caudal artery (CA) and posterior ends of the caudal vein (CV) (dashed box domains with cell indices < 344 in Fig. 1B). These 3 reservoir domains feed RBCs into the simulation domain by making copies of the recycled cells whenever they exited the ends of the reservoir. A key inclusion in our CFD model was the dynamics of the RBC phase. By employing a short-range repulsion force scheme, we could perform RBC to RBC contact and collisions with the vessel lumen wall with high regularity and in a robust manner that ensured RBC mesh did not collapse or inter-penetrate due to excessive deformation (Fig. 1C, Movie 1) – for a detailed explanation on this feature, please refer to materials and methods. To represent the pulsatile flow in the lumen network, we employed pulsatile pressure boundaries at the anterior and posterior ends of the CA (Fig. 1Di) that established the oscillating pressure drop across the CA (Fig. 1Dii) that drives the RBC and plasma flow in the CA (Fig. 1Diii). Likewise, pulsatile pressure boundaries at the anterior and posterior ends of the CV (Fig. 1Ei) established the oscillating pressure drop across the CV (Fig. 1Eii) that drives the RBC and plasma flow in the CV (Fig. 1Eiii). Oscillating pressures defined at the anterior and posterior segments of CA and CV setup the oscillating pressure field within the network that determined the dorsal to ventral pressure distributions in the intersegmental vessels (ISVs). In Fig. 1F, we show the resulting oscillatory dorsal and ventral pressures in aISV3 (arterial intersegmental vessel 3, Fig. 1Fi), oscillatory pressure drop across aISV3 (Fig. 1Fii) and the resulting oscillatory RBC and plasma flow rates (Fig. 1Fiii) in response to the pressure gradients.

**Fig. 1.**
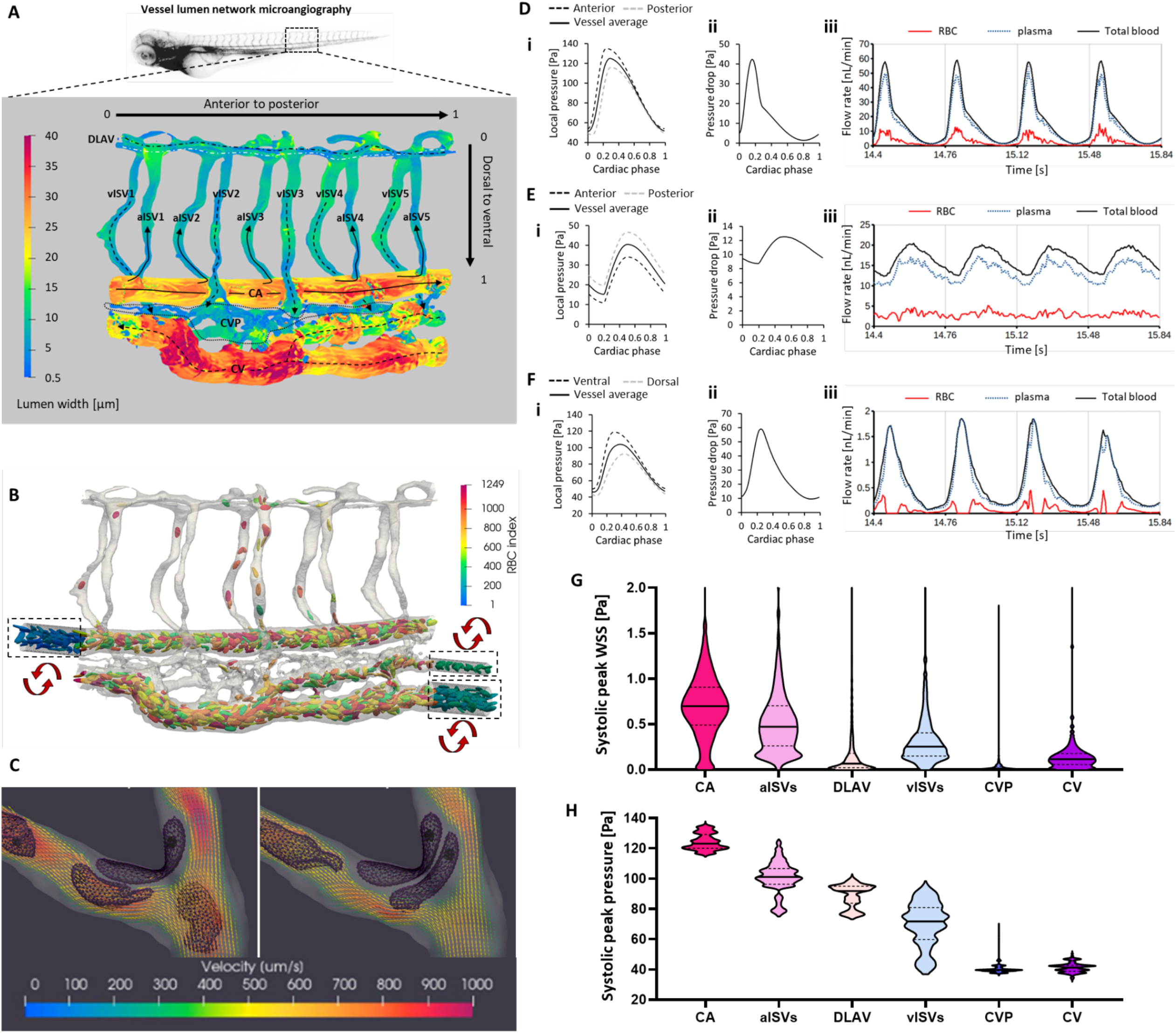
Development of the cell-and-plasma 3-D computational fluid dynamics (CFD) model. **(A)** Morphology of zebrafish trunk network in the caudal vein plexus region obtained using microangiography and confocal microscopy. Shown are the different vessel types of varying diameter (CA: caudal artery, CV: caudal vein, aISV and vISV: arterial and venous intersegmental vessels, DLAV: dorsal longitudinal anastomotic vessel, CVP: caudal vein plexus). **(B)** Red blood cell (RBC) hematocrit in the simulation domain controlled and maintained by recycling cells in the 3 dashed-box domains (RBC indices < 344). **(C)** Collision of RBCs with vessel walls and with one another as part of the explicit consideration of blood dynamics in the CFD model (Movie 1). **(D** and **E)** Pulsatile pressure boundary conditions defined at the anterior and posterior ends of the CA (Di) and CV (Ei) and the resulting pulsatile pressure drop across the CA (Dii) and CV (Eii) that drives the pulsatile flow of RBC and plasma blood phases in the CA (Diii) and CV (Eii). **(F)** Pulsatile pressures arising at the dorsal and ventral ends of aISV3 (i) due to oscillating pressure inputs at anterior and posterior ends of the CA and CV, and the resulting pulsatile pressure drop across aISV3 (ii) that drives the pulsatile flow of RBC and plasma blood phases in aISV3 (iii). **(G)** Hierarchical stratification of WSS in the network predicted by CFD. **(H)** Hierarchical stratification of blood pressure levels in the network predicted by CFD.

We ensured that the CFD flow results were a reasonable match to our experimental reference (Ye et al., 2022) in terms of blood flow peak velocities and time-averaged discharge hematocrit in the various vessel types (Table 1). Summarizing the CFD output, we obtained spatiotemporal maps of wall shear stress (WSS) and lumen blood pressure. From these maps we verified the establishment of WSS hierarchy among vessel types, from high to low (Fig. 1G): CA, aISVs, venous intersegmental vessels (vISVs), CV, dorsal longitudinal anastomotic vessels (DLAV) and the caudal vein plexus capillaries (CVP). As blood flows from high to low pressure regions, the hierarchy of blood pressure levels follows this arterial to venous flow direction. The blood pressure hierarchy given from highest to lowest was CA, aISVs, DLAV, vISVs, CVP and CV (Fig. 1H).

**Table 1.** Comparison of flow velocity and vessel discharge hematocrit in the CA, CV, aISVs and vISVs against experimental reference.

| | | Average peak systolic velocity [$\mu\text{m/s}$] | | | | | Time-averaged discharge hematocrit | | | | |
| --- | --- | --- | --- | --- | --- | --- | --- | --- | --- | --- | --- |
| Data source | No. of embryos at 2dpf | CA | CV | aISVs | vISVs | ISVavg | CA | CV | aISVs | vISVs | ISVavg |
| Ye et al., 2022 | 27 | 2302.5 | 1326.5 | 1002 | 869 | 936 | 16.3 | 16.8 | 4.12 | 4.325 | 4.223 |
| WT model 1 simulation |  | 2307 | 1011 | 1069 | 1062 | 1066 | 17.80 | 18.12 | 4.698 | 3.298 | 3.998 |

### 2.1 RBC hematocrit modulates hemodynamics in a microvascular network

The inclusion of deformable RBCs in the CFD model allowed us to investigate several rheological parameters in the wildtype (WT) geometry. It is a common practice among researchers to employ gata1 knockdown zebrafish models with diminished RBC count (Ransom et al., 1996) to study the effects of blood viscosity reduction on angiogenesis and vessel remodeling (Lee et al., 2016). It is widely assumed that the decrease in blood viscosity will lead to reduced WSS in blood vessels (Vermot et al., 2009). However, when RBCs are removed from the microcirculation two scenarios can occur. First, the adaptive response might be to maintain blood pressure at similar levels as the WT embryo with physiological hematocrit (Fig. 2Ai and Aii). In the second scenario, the adaptive response might be to reduce the blood pressure (Fig. 2Ai versus Aiii) to maintain similar blood flow as the WT embryo with physiological hematocrit (Fig. 2Bi and Biii). Three simulation models were employed to study the implications of the physiological adaptation to RBC removal. Model 1 was the control scenario where RBCs were included in the simulation. Model 2 had no RBCs and maintained the same blood pressure as model 1 at the simulation domain boundary conditions (the nine input boundaries are indicated by black arrows in Fig. 2Ai). Model 3 had no RBCs and maintained the same blood flow rates as model 1 for the CA and CV. Please note that in this article we refer to blood flow as the combined flow of the RBC and plasma mixture while RBC flow specifically refers to the flow of individual phase of discrete RBCs from the mixture. In the case of model 1, blood flow consists of both RBC and plasma flow. However, in the cases of models 2 and 3, the blood flow consists entirely of plasma flow.

**Fig. 2.**
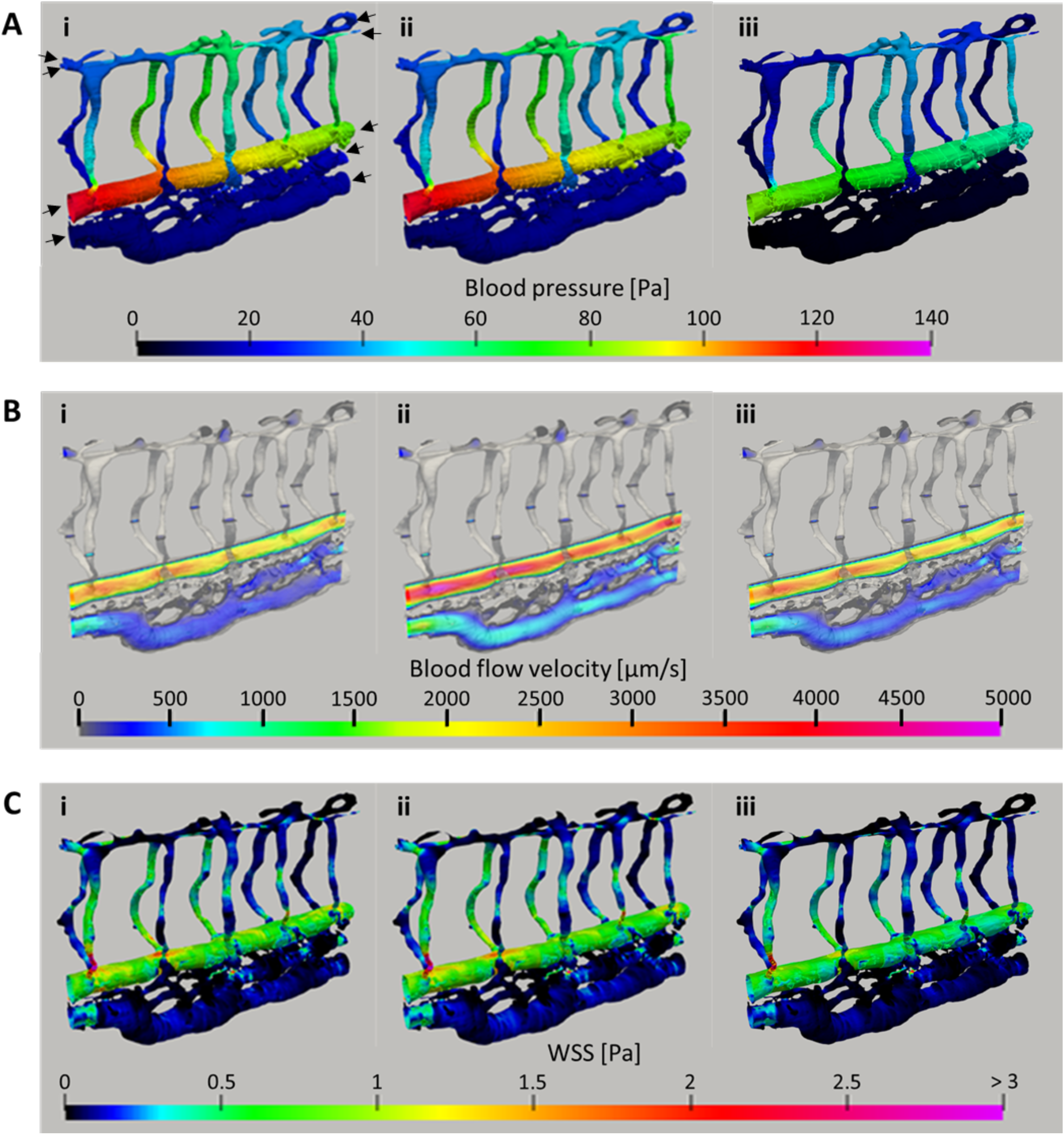
Spatial distribution maps of hemodynamics during the systolic peak of a cardiac cycle for WT flow models with RBCs (control scenario: model 1, i), without RBCs under same blood pressure input as control (model 2, ii), and without RBCs under same blood flow rate input as control (model 3, iii). **(A)** Comparison of the systolic blood pressure amongst the three flow models (Movie 2). **(B)** Comparison of the systolic blood flow velocity amongst the flow models for selected cross-sections of the CA, CV and ISVs (Movie 3). **(C)** Comparison of the systolic WSS amongst the flow models (Movie 4).

The simulations showed a marked increase in the blood flow velocity for model 2 as compared to model 1 and model 3 (Fig. 2B). Notably, because the pressure in the network was similar to model 1, the spatial distribution of WSS in model 2 was also similar (Fig. 2Ci and Cii). Conversely, model 3 showed a general reduction in WSS compared to model 1 (Fig. 2Ci versus Ciii). Movie files of the spatiotemporal oscillations of blood flow velocity, blood pressure and WSS can be viewed in Movie 2, Movie 3 and Movie 4, respectively.

To examine the network perfusion in more detail, we quantified and compared the time-averaged blood flow rate of the pulsatile blood flow in the three models (Fig. 3A). As a result of the reduced blood viscosity under same pressure distribution across the CA and CV, Model 2 displayed 39% and 54% increases in blood flow rate in the CA and CV, respectively, compared to model 1. Conversely, a 27% reduction in the blood pressure in model 3 ensured that the blood flow rates in the CA and CV were similar to model 1 levels. Compared to model 1, blood flow in the ISVs increased for model 2 (+6.93%, +11.2%, +28.0%, +11.4%, +3.57% for aISVs 1 to 5; +6.83%, +2.20%, +27.0%, + 7.49%, +1.49% for vISVs 1 to 5) while they generally decreased for model 3 (−13.0 %, −12.3%, +1.42%, −11.6%, −18.2% for aISVs 1 to 5; −19.6%, −23.3%, −0.498%, −14.4%, −21.4% for vISVs 1 to 5).

**Fig. 3.**
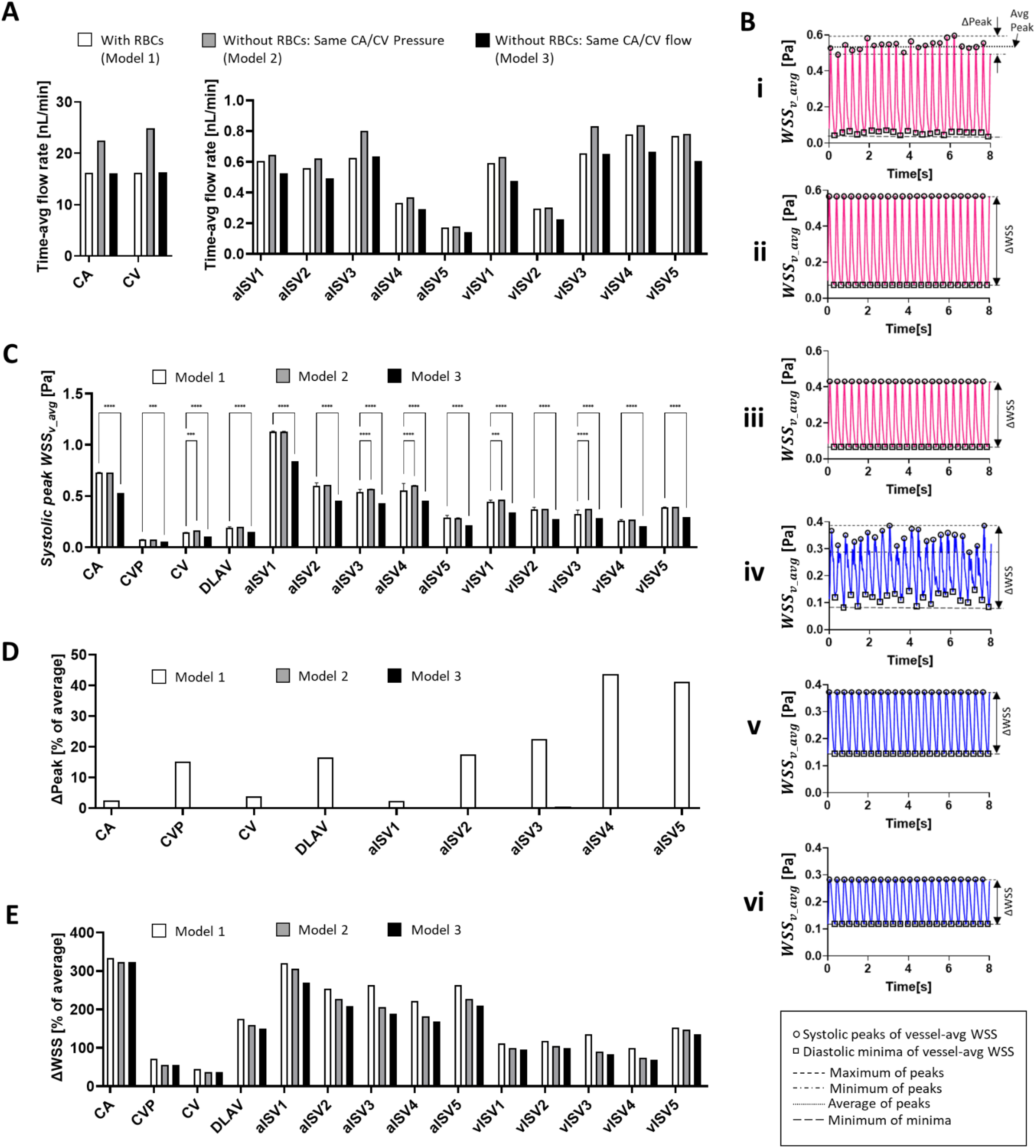
Comparison of hemodynamics for WT flow models with RBCs (control, model 1), without RBCs under same CA/CV blood pressure input as control (model 2), and without RBCs under same CA/CV blood flow rates as with control (model 3). **(A)** Comparison of the time-averaged blood flow rates in the flow models. **(B)** Representative vessel-averaged WSS (*WSS*_*v_avg*_) oscillations against time for aISV3 (i – iii) and vISV3 (iv – vi) under the three flow models. **(C)** Comparison of the *WSS*_*v_avg*_ levels in the flow models. Bar values represent the mean peak WSS and the error bars show the standard deviation of peak WSS. **(D)** Comparison of the *WSS*_*v_avg*_ peak to peak variation width (Δpeak) observed in the three models (refer to Bi for definition). **(E)** Comparison of the *WSS*_*v_avg*_ peak to minima variation width (ΔWSS) observed in the three models (refer to Bi to Bvi for definition).

We averaged the spatial distribution of WSS at each time frame to obtain the oscillation of the vessel-averaged WSS (*WSS*_*v_avg*_) against time in each vessel. Figure 3B shows the temporal profile of *WSS*_*v_avg*_ for models 1 to 3 in aISV3 (i – iii) and vISV3 (iv – vi). From these temporal profiles, we isolated and compared the systolic peak values (circle points in Fig. 3B) in the three models (Fig. 3C – E). The *WSS*_*v_avg*_ levels were similar between the models 1 and 2 except for slight increases in levels for model 2 that were statistically significant in the CV (+13.2%), aISV3 (+5.39%) and aISV4 (+8.88%), and vISV1 (+4.15%) and vISV3 (+15.7%) (Fig. 3C). Model 3 showed lower *WSS*_*v_avg*_ levels than model 1 in all vessel regions (−27.3% in CA; −25.2% in CVP; −25.6% in CV; −21.5% in DLAV; −25.5%, −23.8%, −20.6%, −17.8%, −24.7% for aISVs 1 to 5; −23.4%, −25.1%, −12.2%, −20.5%, −24.6% for vISVs 1 to 5). The temporal fluctuation in peak *WSS*_*v_avg*_ was quantified in terms of the peak-to-peak fluctuation (Δpeak), as shown in Fig. 3D. Compared to models 2 and 3, only model 1 demonstrated fluctuation in *WSS*_*v_avg*_ values (Fig. 3B, error bars in 3C and Fig. 3D). Consequently, the added fluctuation to peak and minima *WSS*_*v_avg*_ from random RBC passages in model 1 resulted in a wider peak-to-minima range (ΔWSS – see Fig. 3B for definition) than models 2 and 3 (Fig. 3E). Regardless of the physiological adaptation applied in models 2 and 3, the consistent phenomena we observed from RBC removal was a lack of irregularity in the temporal oscillation of WSS.

In summary, we found the physiological target of adaptation (adaptive constraint) following a blood viscosity reduction to profoundly affect the resulting hemodynamic outcome. If blood pressure maintenance is the physiological adaptation after the RBC removal, WSS levels generally stay the same but blood perfusion increases. However, if perfusion levels are held constant then both blood pressure and WSS in the vessel network will fall in response to the RBC removal.

Accordingly, we sought to verify which was the likelier scenario for adaptation by imaging the blood flow velocity in Gata1 morpholino-injected zebrafish embryos at 2 dpf. While the treatment does not completely deplete RBCs, it significantly reduces the hematocrit while still allowing RBCs to be present for velocity tracking and blood flow assessment. In the experiment, we examined 7 Gata1 morphants at 2 dpf with varying degrees of hematocrit reduction against 5 wildtype 2 dpf zebrafish (Fig. 4, Table 2 and Section A of supplementary material). We observed that hematocrit reduction did not significantly alter the heart rate, thus suggesting that the blood pressure was likely maintained following a hematocrit reduction (Fig. 4A). Furthermore, we noted that the blood flow velocity (as indicated by RBC velocity) gradually increased in the Gata1 morphants as the RBC flux concentration in the CA and CV was decreased (Fig. 4B). This second point corresponds with the CFD demonstration of model 2 where both the blood flow velocity (Fig. 2B) and blood flow rate in the CA (Fig. 3A) for model 2 were significantly higher than model 1. Hence, we believe that RBC removal from zebrafish embryos is not likely to automatically reduce WSS in the lumen network as assumed in model 3. Instead, our CFD simulations and Gata1 morpholino experiment suggest that hematocrit reduction increases blood flow while maintaining similar levels of blood pressure and WSS.

**Table 2.** Comparison of RBC flux concentration and flow velocity in Gata1 morpholino (Gata1 MO)-injected 2 dpf zebrafish embryos versus wild-type (WT).

| | | Heartrate | peak velocity [$\mu\text{m/s}$] | | |
| --- | --- | --- | --- | --- | --- |
| Group | Fish # | [BPM] | CA | CV | ISV |
| Gata1 MO (0.1uM) | 1 | 174 | 2233.793 | 1765.326 | 1144.232 |
| Gata1 MO (0.1uM) | 2 | 174 | 3921.125 | 2582.458 | 640.4613 |
| Gata1 MO (0.1uM) | 3 | 186 | 2721.673 | 1996.709 | 557.566 |
| Gata1 MO (0.1uM) | 4 | 156 | 2933.562 | 2446.564 | 1025.112 |
| Gata1 MO (0.1uM) | 5 | 180 | 2843.204 | 1921.642 | 1295.455 |
| Gata1 MO (0.1uM) | 6 | 147 | 3110.341 | 2745.665 | 647.62 |
| Gata1 MO (0.1uM) | 7 | 150 | 2082.425 | 1904.402 | 843.8913 |
| WT Control | 1 | 168 | 2267.795 | 1621.395 | 1321.984 |
| WT Control | 2 | 168 | 2371.926 | 2033.35 | 1612.413 |
| WT Control | 3 | 156 | 2239.302 | 1707.77 | 1189.453 |
| WT Control | 4 | 156 | 2033.138 | 1672.274 | 1365.036 |
| WT Control | 5 | 162 | 2691.048 | 1571.064 | 979.5366 |

**Fig. 4.**
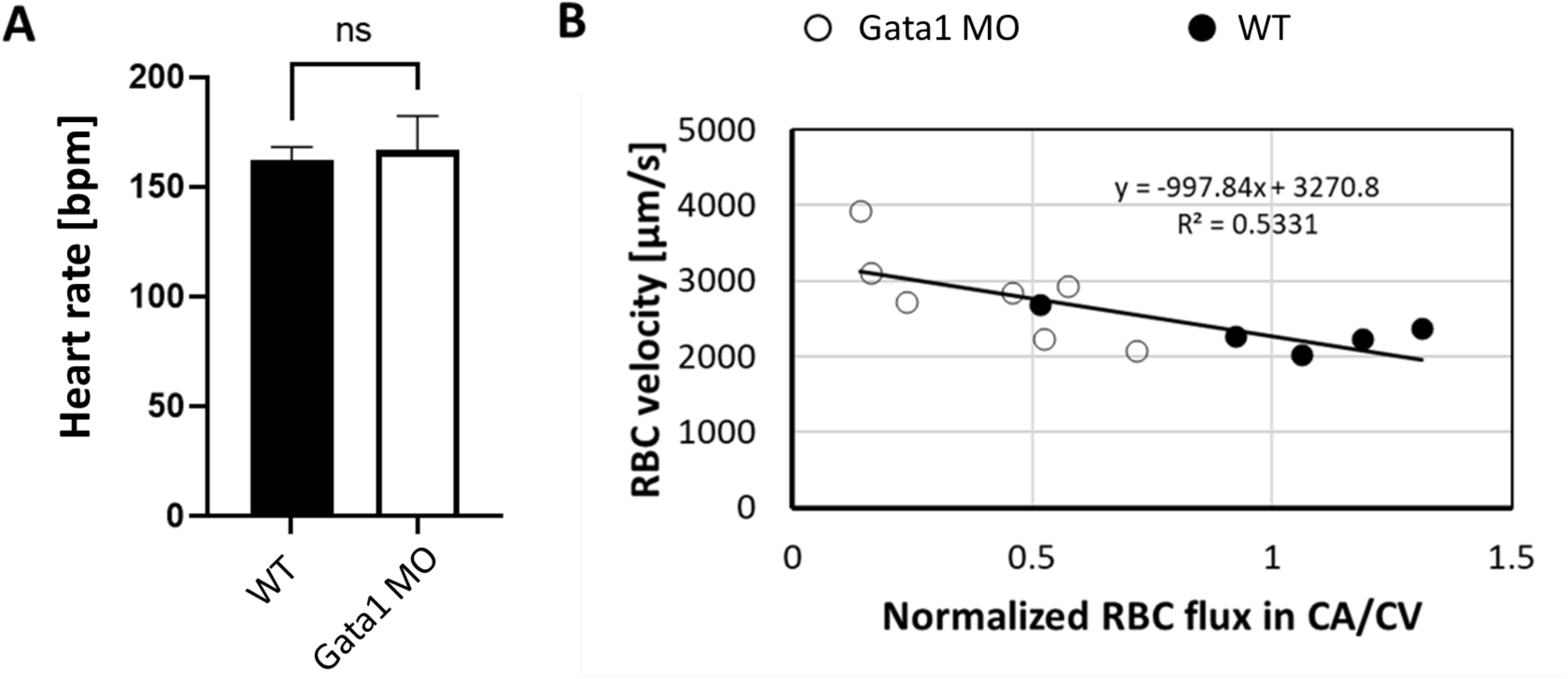
Heart rate in beats per minute [bpm] **(A)** and RBC velocity in the caudal artery **(B)** for zebrafish at 2 dpf after hematocrit reduction in the vascular network in response to Gata1 morpholino treatment. The normalized RBC flux indicates control level hematocrit at a value of 1.

### 2.2 Ventral narrowing of ISV diameter tunes network perfusion and stress gradients

While the application of 3D CFD modeling of hemodynamics in the zebrafish vascular network is not completely novel, a realistic representation of local topologies in the network is a feature often ignored in many CFD models that represent the lumen cross-sectional profile to be smooth and circular (Roustaei et al., 2022; Zhou et al., 2021). Most importantly, the vessel lumen narrows and dilates along the ISV axis at seemingly predestined locations (Fig. 5Ai and iii). Here, we performed parametric modeling of the diameter variation to elucidate on how patterns of diameter variation along vessel segments tune the resulting hemodynamics for specific WSS and pressure distribution patterns. We compared WT lumen diameters obtained from real geometry (WT) against an idealized network where vessel segments had no variation in the lumen diameter along the ISV axis (Fig. 5Aii and iv). We termed this idealized network the smooth-geometry model (SGM). The segment diameters employed in the SGM were obtained from the segment-averaged diameters in the WT network. For this comparison, we maintained the same pressure input at model domain boundaries.

**Fig. 5.**
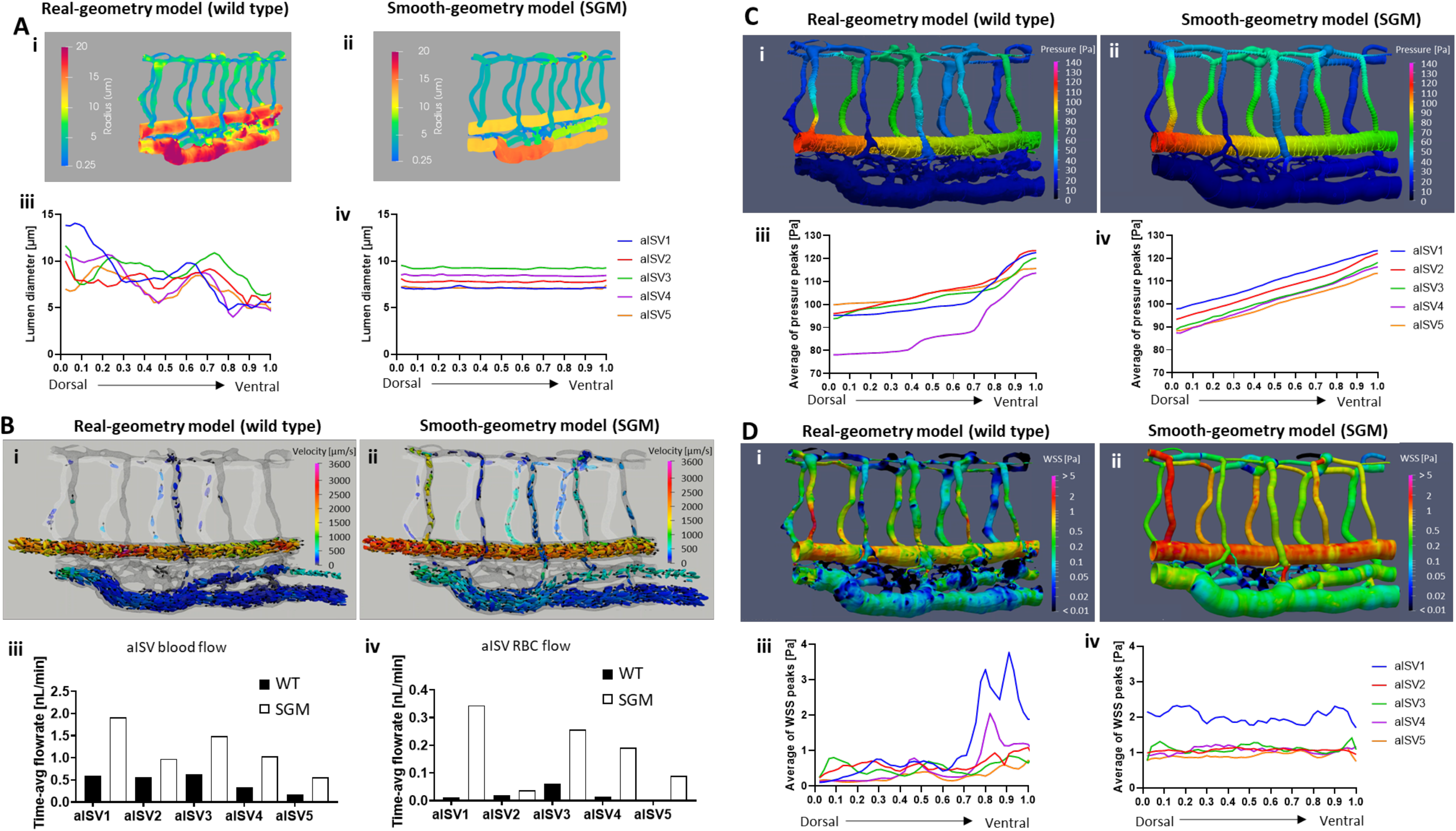
Comparison of hemodynamics between the wild type real-geometry model (WT) and an idealized smooth-geometry model (SGM). **(A)** Spatial distribution maps of lumen diameter in WT (i) versus the SGM (ii) and the diameter gradient in aISVs for WT (iii) versus the SGM (iv) **(B)** Representative spatial distribution maps of RBC flow and velocities in WT (Movie 5) (i) versus the SGM (Movie 6) (ii) at systolic flow peaks; Time-averaged blood flow rates (iii) and RBC flow rates (iv) in the individual aISVs. **(C)** Representative spatial distribution maps of blood systolic pressure in WT (Movie 7) (i) versus SGM (Movie 8) (ii) and the resulting pressure gradients in aISVs for WT (iii) versus SGM (iv). **(D)** Representative spatial distribution maps of systolic WSS in WT (Movie 9) (i) versus SGM (Movie 10) (ii), and the resulting WSS gradients in aISVs for WT (iii) versus SGM (iv).

The SGM showed significantly increased RBC perfusion into the aISVs compared to WT (Fig. 5Bi – Movie 5 and ii – Movie 6). Quantification revealed 218%, 74%,138%, 213% and 228% increase in time-averaged blood flow in aISVs 1 to 5, respectively, for the SGM compared to WT (Fig. 5Biii). The time-averaged RBC flow rate was increased by 3090%, 191%, 413%, 1259% and 18686% in aISVs 1 to 5, respectively, for the SGM compared to WT (Fig. 5Biv). Note that in aISVs 1, 4 and 5 the significant percentage increase in RBC flux were situations where the WT model saw diminishingly low RBC flux into these aISVs. These dramatic increases in RBC perfusion in the SGM highlighted the role of lumen diameter narrowing in aISVs seen in the WT model (Fig. 5Ai and ii) but non-existent in the SGM (Fig. 5Aiii and iv). By narrowing the entrance of aISVs, the amount of RBC entry into aISVs was significantly restricted in the WT model.

Blood pressure distribution between WT and SGM was similar in the CA and CV (Fig. 5Ci – Movie 7 and ii – Movie 8) due to the application of same boundary conditions for the two models. However, there was a marked difference in the systolic pressure gradient across aISVs between the two models. The WT model showed a rising pressure gradient towards the ventral regions due to the lumen narrowing in these regions (Fig. 5Ciii). As the narrowest regions of a vessel segment provide the highest resistance against flow, these regions require proportionately higher pressure gradients to drive blood flow. Conversely, the SGM showed a constant pressure gradient across the aISVs due to the smooth and constant lumen diameter within aISVs (Fig. 5Civ).

Spatial distribution maps of systolic WSS showed spatial irregularity in the WSS level for WT (Fig. 5Di, Movie 9) whereas the SGM displayed smooth distributions of WSS where the levels were stratified by vessel segment (Fig. 5Dii – Movie 10). There was a positive dorsal-to-ventral WSS gradient in the aISVs for WT (Fig. 5Diii) that was absent in the SGM aISVs (Fig. 5Div). Additionally, the vessel-average systolic WSS in the aISVs for the SGM was higher than that of the WT model. This was due to the increased RBC flux and concomitant blood viscosity rise in the SGM aISVs.

In summary, dorsal-to-ventral narrowing of aISVs at 2 dpf appears to confer the network with flow and WSS mitigation features primarily by restricting the entry of RBCs into the aISVs. Furthermore, the lumen narrowing is essential to maintain the dorsal-to-ventral WSS gradient which may serve as an important mechanotaxis cue in the guidance of endothelial cell migration (Tardy et al., 1997) and polarization (Simmers et al., 2007).

### 2.3 Hemodynamic forces in vascular networks of altered morphology or pattern

#### 2.3.1 Lumen diameter alteration

After examining the basic features of the WT network geometry, blood rheology and their relationship with WSS and pressure distributions, we applied the CFD model to alterations in network morphology. We generated in silico vascular networks based on vessel morphologies obtained from experimental manipulations. We took the WT as a baseline geometry and modified it in accordance with reported metrics of the lumen geometry observed from confocal images.

The first group of experiment-based networks were the Marcksl1 phenotypes. In the case where Marcksl1, an actin bundling protein, is overexpressed specifically in endothelial cells (Marcksl1OE), the modified network is a lumen bulbous-expansion phenotype (Kondrychyn et al., 2020) compared to WT (Fig. 6A vs Fig. 6B), where we selected four local regions for lumen volume dilation (dashed yellow circles in Fig. 6B): Region 1: vISV2 (+239%), Region 2: aISV3 (+288%), Region 3: DLAV3 (+300%), Region 4: DLAV4 (+272%). In the case of *marcksl1a* and *marcksl1b* double knockout zebrafish (Marcksl1KO), the modified network is a reduced lumen diameter phenotype (Kondrychyn et al., 2020) (Fig. 6C – E): aISV (−20%), vISV (−23%), DLAV (−21%), CA (−10%), CV (−5%).

**Fig. 6.**
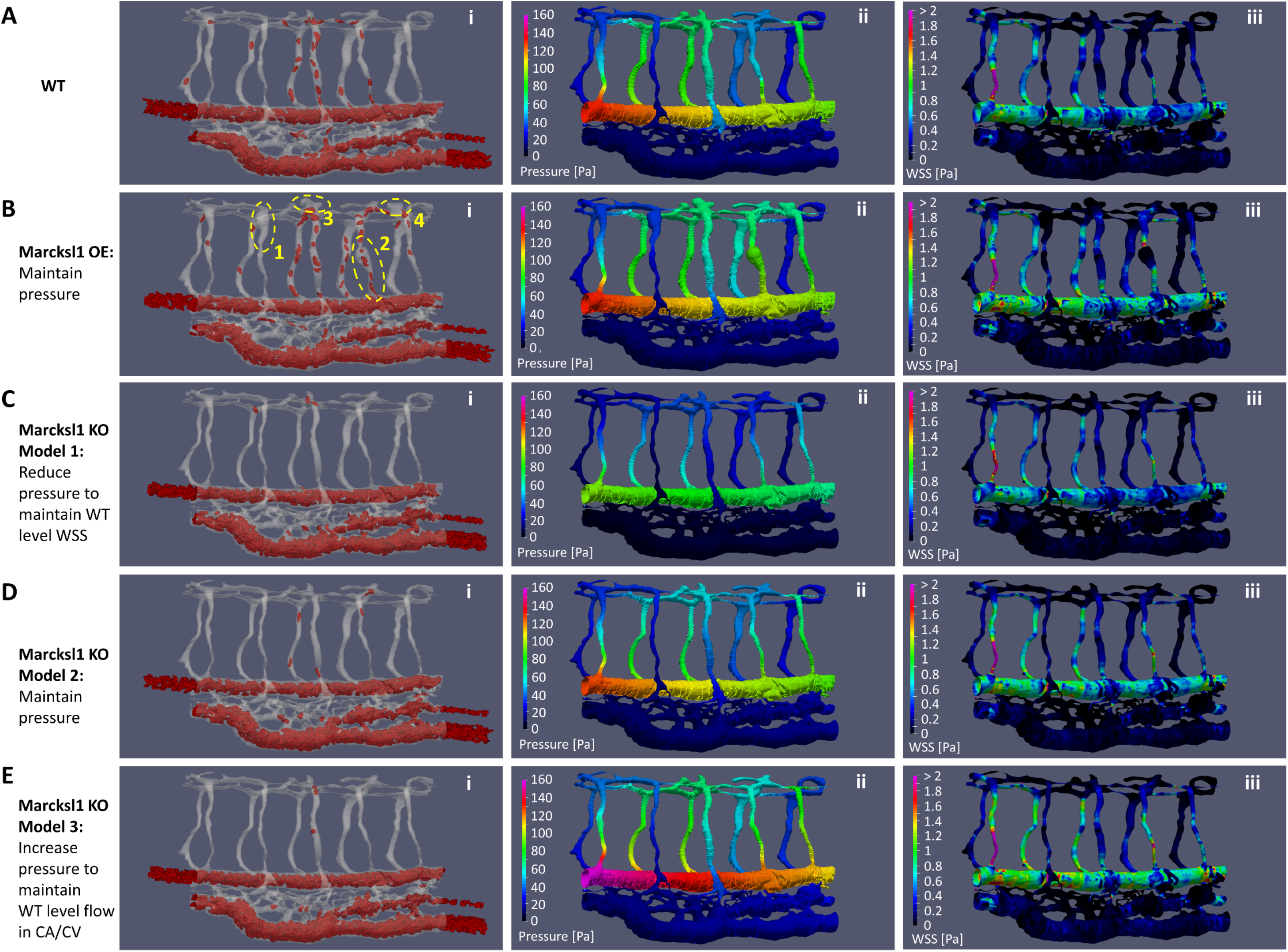
Hemodynamics adaptation scenarios for Marcksl1 OE and Marcksl1 KO. **(A)** WT levels of RBC perfusion (Movie 5) (i), systolic blood pressure distribution (Movie 7) (ii) and systolic WSS distribution (Movie 9) (iii). **(B)** Marcksl1 OE model with similar blood pressure as WT: spatial maps of RBC perfusion (Movie 11) where yellow dashed circles indicate the locally dilated lumen region compared to WT (i), systolic blood pressure distribution (see Movie 12 for pressure oscillations) (ii) and systolic WSS distribution (see Movie 13 for WSS oscillations) (iii). **(C)** Marcksl1 KO Model 1 with reduced blood pressure: spatial maps of RBC perfusion (Movie 14) (i), systolic blood pressure distribution (see Movie 15 for pressure oscillations) (ii) and systolic WSS distribution (see Movie 16 for WSS oscillations) (iii). **(D)** Marcksl1 KO Model 2 with similar blood pressure as WT: spatial maps of RBC perfusion (Movie 17) (i), systolic blood pressure distribution (see Movie 18 for pressure oscillations) (ii) and systolic WSS distribution (see Movie 19 for WSS oscillations) (iii). **(E)** Marcksl1 KO Model 3 with increased blood pressure: spatial maps of RBC perfusion (Movie 20) (i), systolic blood pressure distribution (see Movie 21 for pressure oscillations) (ii) and systolic WSS distribution (see Movie 22 for WSS oscillations) (iii).

We maintained the same pressure in the Marcksl1OE model as the WT model. Since the lumen dilations were localized to a few locations in the ISV network, we did not expect the overall systemic flow to be altered for the Marcksl1OE network. However, we did not expect the same outcome for the Marcksl1KO network. Since the lumen reduction was systemically applied across the entire network, the systemic flow in the Marcksl1KO will likely undergo physiological adaptation. Consequently, we have considered three hemodynamics adaptation scenarios for the Marcksl1KO network. In model 1, the blood pressure was reduced (108 Pa peak systolic blood pressure in model 1 versus 140 Pa in the WT) to maintain a WSS level close to the WT model (Fig. 6Cii and iii). In model 2, the blood pressure was maintained to same levels as WT, thereby causing a proportionate increase in WSS levels due to the lumen reduction (Fig. 6Dii and iii). In model 3, the blood pressure was increased (160 Pa peak systolic blood pressure) to maintain the same blood flow rate (16.0 nL/min) as WT (16.2 nL/min) despite the lumen reduction (Fig. 6Eii and iii). This adaptation scenario for model 3 resulted in a more significant rise in WSS than observed in model 2.

We sought to verify which adaptation scenario was most likely by imaging the blood flow in Marcksl1KO zebrafish (15 embryos) versus WT (5 embryos) at 2 dpf (Table 3, Section B and C of supplementary material). Through fluorescent labelling of RBCs using *Tg(gata1:dsRed)^sd2^* zebrafish, we measured the RBC flux into the ISV networks. RBC-perfused lumen diameters of the trunk vascular network were obtained by performing maximum intensity time-stack projections of the RBC trajectories in accordance with the procedure described in Ye and colleagues (Ye et al., 2022). From these two measurements we could classify the degree of phenotyping for Marcksl1KO zebrafish in terms of RBC perfusion levels 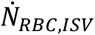 [RBCs/s per ISV] into the ISV network. Representative images of these groups are shown in Fig. 7 Ai – iii. In the high perfusion group were 4 embryos with 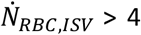. In the moderate (mid) perfusion group were 6 embryos with 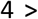 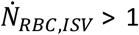. In the low perfusion group were 3 embryos with 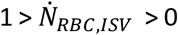. In the zero perfusion group were 2 embryos with no RBC perfusion into the ISV network (Fig. 7B).

**Table 3:** Comparison of flow velocity and RBC perfusion levels in the ISV networks of Marcksl1 KO and WT zebrafish embryos at 2 dpf from experiments and simulations.

| Group | Remarks | Fish # | Heartrate<br>[BPM] | peak velocity [$\mu\text{m/s}$] | | | RBC flux per<br>ISV [rbc/s] |
| --- | --- | --- | --- | --- | --- | --- | --- |
|  |  |  |  | CA | CV | ISV |  |
| WT Control | Good ISV perfusion | 1 | 168 | 2267.795 | 1621.395 | 1321.984 | 4.79 |
| WT Control | Good ISV perfusion | 2 | 168 | 2371.926 | 2033.35 | 1612.413 | 6.8 |
| WT Control | Good ISV perfusion | 3 | 156 | 2239.302 | 1707.77 | 1189.453 | 8.12 |
| WT Control | Good ISV perfusion | 4 | 156 | 2033.138 | 1672.274 | 1365.036 | 5.48 |
| WT Control | Moderate ISV perfusion | 5 | 162 | 2691.048 | 1571.064 | 979.5366 | 0.88 |
| Marcksl1 KO | Good ISV perfusion | 6 | 162 | 2084.462 | 1864.945 | 1297.457 | 5.22 |
| Marcksl1 KO | Good ISV perfusion | 10 | 156 | 2348.667 | 1829.953 | 1967.858 | 6.42 |
| Marcksl1 KO | Good ISV perfusion | 13 | 138 | 2713.477 | 1959.212 | 1642.746 | 5.29 |
| Marcksl1 KO | Good ISV perfusion | 15 | 138 | 2622.458 | 1968.722 | 1659.116 | 6.14 |
| Marcksl1 KO | Moderate ISV perfusion | 1 | 138 | 2341.108 | 1686.766 | 1040.722 | 2.24 |
| Marcksl1 KO | Moderate ISV perfusion | 2 | 144 | 2035.337 | 1933.135 | 1308.026 | 3.33 |
| Marcksl1 KO | Moderate ISV perfusion | 12 | 138 | 2228.737 | 1279.644 | 950.9839 | 1.98 |
| Marcksl1 KO | Moderate ISV perfusion | 3 | 162 | 2443.097 | 2057.51 | 1094.051 | 1.2 |
| Marcksl1 KO | Moderate ISV perfusion | 7 | 140 | 1927.842 | 1743.457 | 1474.2 | 1.11 |
| Marcksl1 KO | Moderate ISV perfusion -abnormal<br>CA morphology | 9 | 140 | 1351.363 | 876.3137 | 1457.34 | 1.05 |
| Marcksl1 KO | Poor ISV perfusion | 5 | 140 | 1073.229 | 910.1901 | 411.1627 | 0.05 |
| Marcksl1 KO | Poor ISV perfusion | 8 | 126 | 1701.285 | 1186.483 | 650.5616 | 0.16 |
| Marcksl1 KO | Poor ISV perfusion | 14 | 120 | 940.721 | 496.5036 | 508.7208 | 0.04 |
| Marcksl1 KO | No ISV perfusion | 4 | 150 | 680.1337 | 1168.777 | 0 | 0 |
| Marcksl1 KO | No ISV perfusion | 11 | 144 | 0 | 0 | 0 | 0 |
| WT simulation | Caudal artery peak pressure 140 Pa | - | 167 | 2306.18 | 1010.87 | 1066 | 3.49198 |
| Marcksl1 KO simul. | Caudal artery peak pressure 108 Pa | Model 1 | 130 | 1445.41 | 580.783 | 477.265 | 0.43382 |
| Marcksl1 KO simul. | Caudal artery peak pressure 140 Pa | Model 2 | 130 | 1922.51 | 769.522 | 804.37 | 0.67287 |
| Marcksl1 KO simul. | Caudal artery peak pressure 160 Pa | Model 3 | 167 | 2388.76 | 944.235 | 929.28 | 0.53711 |

**Fig. 7.**
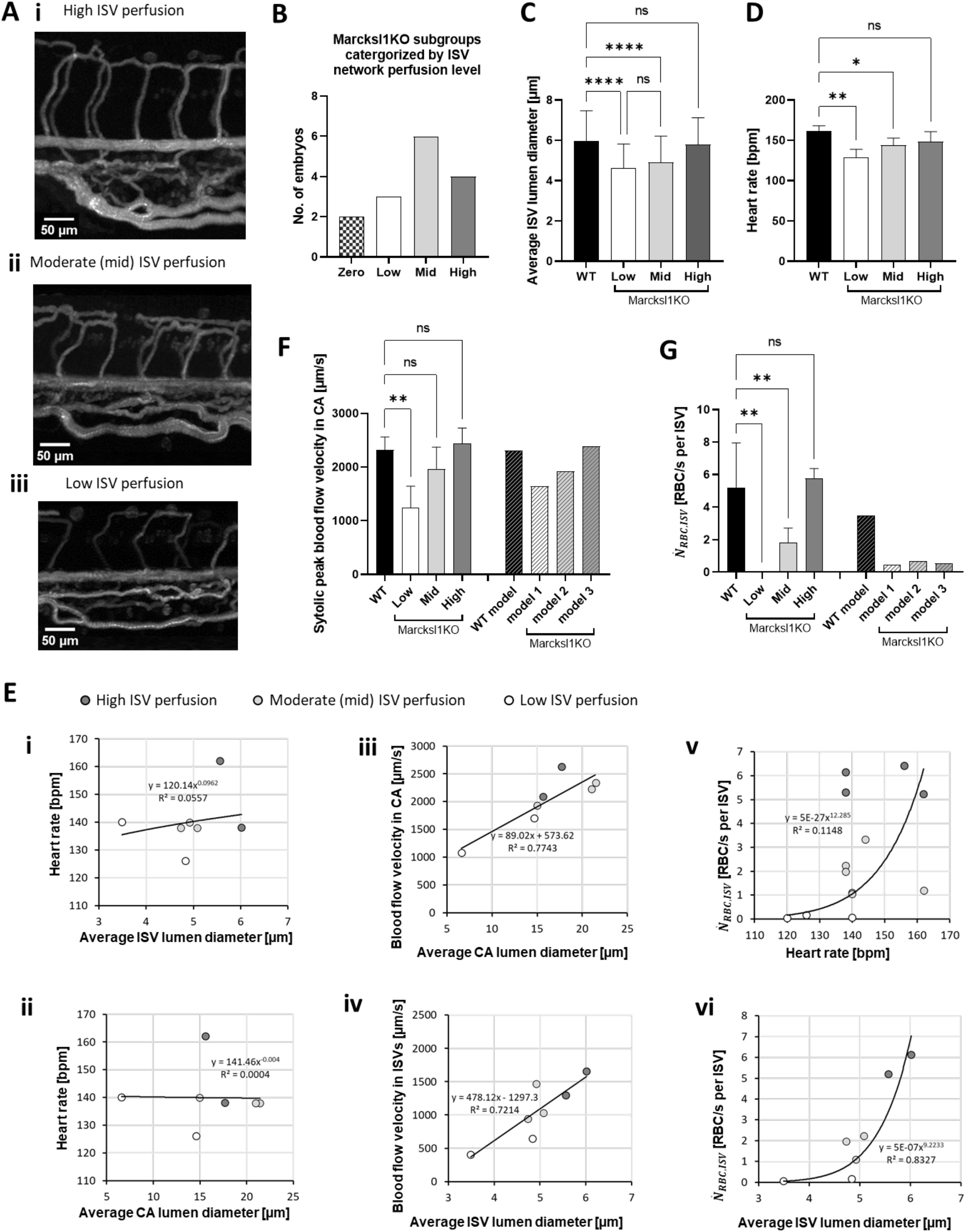
Characterization of RBC flow in the local trunk network for 2 dpf Marcksl1 KO experiment. **(A)** Representative maximum intensity time-stack projection images of lumen perfusion by RBC flow in the ISV networks from high (i), moderate/mid (ii) to low (iii) perfusion. **(B)** Population sample distribution grouped by ISV RBC-perfusion level from high to zero. **(C)** Comparison of average ISV lumen diameters of Marcksl1 KO zebrafish in different perfusion level groups versus WT. **(D)** Comparison of heart rate of Marcksl1 KO zebrafish versus WT. **(E)** Relationships between lumen diameter alteration and cardiovascular adaptation for 2 dpf Marcksl1 KO experiment: (i) heart rate against average ISV lumen diameter; (ii) heart rate against average CA lumen diameter; (iii) CA systolic blood flow velocity against average CA lumen diameter; (iv) ISV peak blood flow velocity against ISV lumen diameter; (v) average RBC flux into ISV network 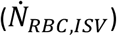 against heartrate; (vi) 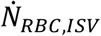 against average ISV lumen diameter. **(F)** Comparison of systolic peak blood flow velocity in the CA for the Marcks1 KO zebrafish versus the WT and simulation models. **(G)** Comparison of systolic peak blood flow velocity in the ISV network for the Marcks1 KO zebrafish versus the WT and simulation models.

There was a statistically significant reduction in average ISV lumen diameter for the low perfusion group (4.61 μm) and moderate perfusion group (4.91 μm) compared to high perfusion group (5.80 μm) (Fig. 7C). In terms of heart rate, there was a statistically significant reduction in the average heart rate for the low (129 bpm) and mid (144 bpm) groups compared to WT (162 bpm), but not for the high group (149 bpm) (Fig. 7D). We interpreted this to suggest a lowered blood pressure adaptation in the experimental Marcksl1KO subset showing flow phenotyping of reduced ISV perfusion by RBCs (9 out of 13 ISV-perfused mutants). Regression analysis however showed that the heart rate in the mutants did not correlate well with neither the ISV nor CA lumen diameter variation (Fig. 7Ei and ii). Instead, in both the CA and ISVs, it was observed that blood flow velocity was highly correlated with the vessel lumen diameter (Fig. 7Eiii and iv). Likewise, the RBC perfusion degree in ISV networks given by 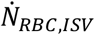 was poorly correlated with the heartrate (Fig. 7Ev) but instead decreased exponentially with a reduction in the average ISV lumen diameter (Fig. 7Evi). To summarize the experiments, lumen reduction in the Marcksl1KO zebrafish appears to elicit a strong trend of blood flow velocity reduction and a consistent level of pressure reduction in mutants (versus WT) that exhibit an ISV perfusion reduction phenotype. Furthermore, ISV perfusion by RBCs appears to drop dramatically to zero when the lumen diameter reduction approaches 3 – 4 μm as demonstrated by results in Fig. 7Evi.

We sought to match these three conditions (velocity trend, pressure trend and ISV perfusion trend) best with the 3 model scenarios we performed for the Marcksl1KO network. Results in Fig. 7F show that models 1 to 3 appear to show a qualitative match with the experimental trend of ascending CA blood velocity. However, the pressure in model 3 was set at 15% higher than WT which we did not observe by proxy of the heart rate trend between WT and Marcksl1KO zebrafish. Furthermore, we did not see an increase in RBC perfusion for the ISV network in model 3 (Fig. 7G). Hence, model 3 which describes the adaptive target of maintaining WT level blood flow rate fails to describe any of the three experimental scenarios, and can be ruled out as a possible adaptation scenario. Likewise, model 2 that posits a maintenance of blood pressure following lumen reduction is not corroborated by the experimental data. Considering the relative level change in CA blood flow velocity between WT and the mutant in the low group (−46.7%) and mid group(−15.2%), model 1 (−28.7% versus WT model) could reasonably match the flow phenotype that is a transition from mid to low RBC perfusion in RBCs. This is further supported by the ISV lumen reduction applied in our three Marcksl1KO models (−21.5%), which falls halfway between the low group (−28.2%) and mid group (−15.3%) from the experiment. Based on the combined findings from CFD and experiment, we posit that model 1, which describes a lowered blood pressure adaptation appears to be the likeliest physiological adaptation for Marcksl1KO networks.

With the likely adaptation scenario for Marcksl1KO identified, we compared the WSS and blood pressure levels in the various vessel types and their respective stress gradients for the Marcks1KO (model 1), Marcksl1OE and WT model networks. Figure 8A shows the time-averaged pressure drop (*P*_*drop_v*_) across ISVs (ventral-to-dorsal in aISVs and dorsal-to-ventral in vISVs) for the various Marcksl1 models. As a result of the reduced arterial pressure adaptation, Marcksl1 KO model 1 ISV networks saw lowered flow-driving pressure gradients than compared to the WT (−21.5%, +3.29%, +4.00%, −24.8% and +6.57% difference in aISVs 1, 2, 3, 4 and 5; −18.5%, −22.4%, −17.33%, −19.9% and −19,9% in vISVs 1, 2, 3, 4 and 5). The exceptions of aISVs 2, 3 and 5 that showed slightly higher *P*_*drop_v*_ than WT was due to the local rerouting of network flow as a result of the network-wide lumen reductions. Naturally, the reduced *P*_*drop_v*_ levels in Marcksl1 KO model 1 coupled with the lumen diameter reduction resulted in substantial decreases in blood flow rate (Fig. 8B: −49.3%, −52.6%, −51.0%, −41.8% and −44.3% in aISVs 1, 2, 3, 4 and 5 compared to WT; −47.6%, −32.8%, −40.5%, −44.1% and −50.5% in vISVs 1, 2, 3, 4 and 5 compared to WT) and RBC flow rate (Fig. 8C: −72.8%, −100%, −73.4%, −100% and −100% in aISVs 1, 2, 3, 4 and 5 compared to WT; −100%, −100%, −90% and −100% in vISVs 1, 2, 3 and 4 compared to WT) for the ISVs. Consequently, despite lumen reductions, we observed either statistically significant decreases in the systolic peak *WSS*_*v_avg*_ (for aISV1, aISV4, vISV1 and vISV5) or similar levels in ISVs for model 1 Marksl1 KO versus WT model (Fig. 8D).

**Fig. 8.**
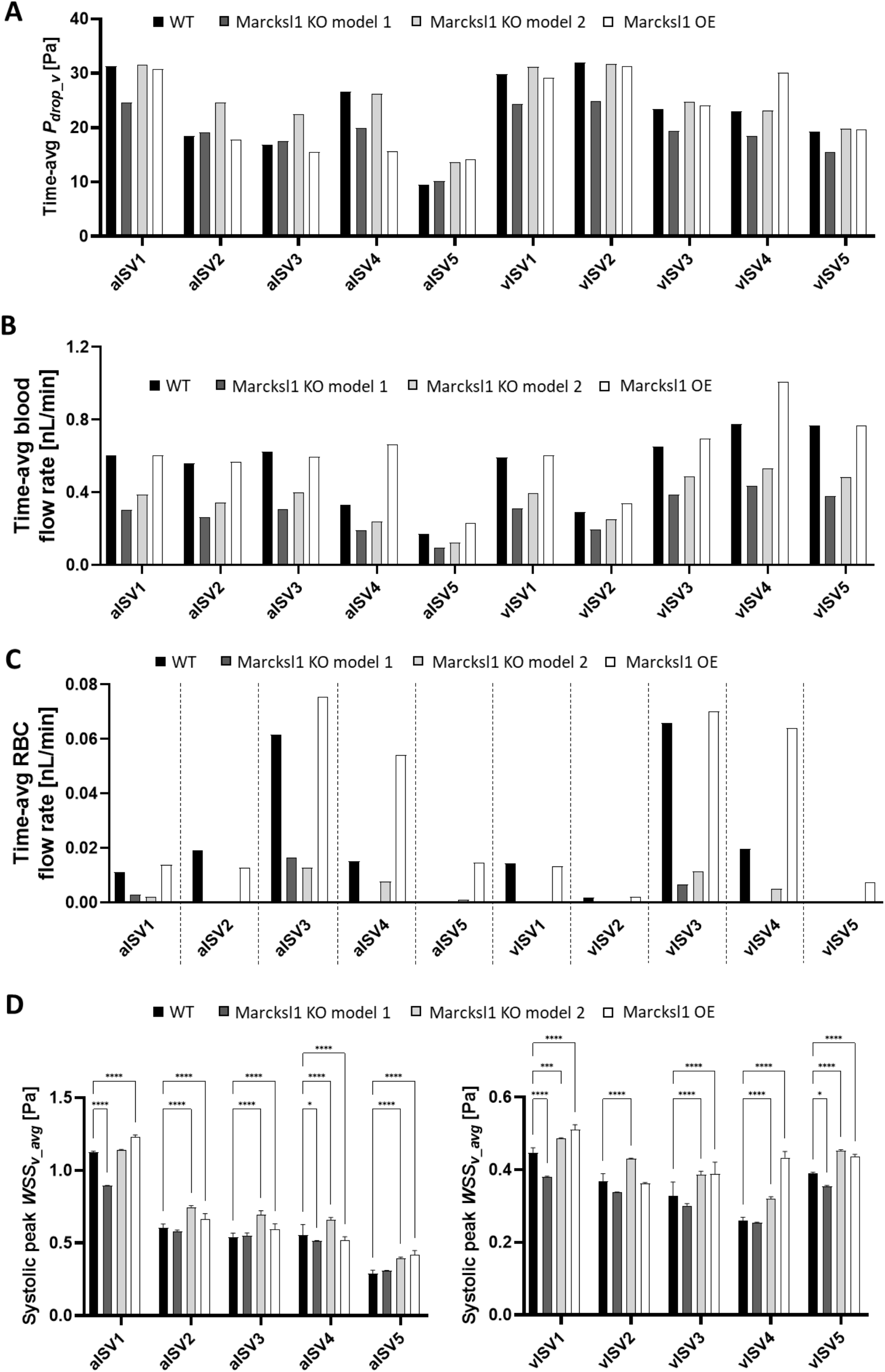
Comparison of hemodynamic parameters amongst WT and Marcksl1 network models. **(A)** Time-averaged pressure drop (*P*_*drop_v*_) along vessel segments amongst models; *P*_*drop_v*_ is taken to be positive in the ventral to dorsal direction for aISVs and dorsal to ventral direction for vISVs. **(B)** Time-averaged blood flow rate into ISVs amongst models. **(C)** Time-averaged RBC flow rate into ISVs amongst models. **(D)** Systolic peak vessel-averaged WSS (*WSS*_*v_avg*_) in ISVs amongst models.

In the Marcksl1 OE model, we saw a general maintenance of *P*_*drop_v*_ levels (Fig. 8A) and blood flow rates (Fig. 8B) for most ISVs except aISV4 (−40.9% change in *P*_*drop_v*_ levels and +101% in blood flow rate from WT), aISV5 (+49.2% change in *P*_*drop_v*_ levels and +35.7% in blood flow rate from WT) and vISV4 (+30.8% change in *P*_*drop_v*_ levels and +29.4% in blood flow rate from WT). These were direct effects from the lumen dilations applied in regions 2, 3 and 4 (Fig. 6Bi) that allowed lower resistance ISV-DLAV flow paths for blood transport. Likewise, the RBC perfusion for aISV3, aISV4, aISV5, vISV4 and vISV5 saw considerable increase in levels compared to WT (Fig. 8C: +22.2%, +256%, +2960%, +226% in aISV3, aISV4, aISV5, vISV4 and +0.008 nL/min versus 0 in vISV5). Consequently, there were statistically significant increase in the systolic peak *WSS*_*v_avg*_ for most ISVs in the Marcksl1 OE model compared to WT except vISV2 showing statistically similar levels as WT and aISV4 showing statistically lower systolic peak *WSS*_*v_avg*_ levels than WT. Generally, the effect of local lumen dilation was to permit higher blood and RBC perfusion, thereby augmenting both flow velocity and viscosity and leading to WSS rises in the ISV network. However, when the dilation was applied over a fairly broad segment of the ISV, as in the case of region 2 in aISV4 then WSS was lowered due to over-dominant WSS mitigation effects of the lumen expansion.

Movies pertaining to Figure 6 can be found in the supplementary information section (Fig. 6Ai – Movie 5; Aii – Movie 7; Aiii – Movie 9; Bi – Movie 11; Bii – Movie 12; Biii – Movie 13; Ci – Movie 14; Cii – Movie 15; Ciii – Movie 16; Di – Movie 17; Dii – Movie 18; Diii – Movie 19; Ei – Movie 20; Eii – Movie 21; Eiii – Movie 22)

#### 2.4.2 ISV-DLAV network mispatterning

The PlexinD1-Semaphorin signaling pathway controls the patterning of ISVs by repelling ECs from migrating into the somites during sprouting angiogenesis (Torres-Vázquez et al., 2004). Here we represent a misregulation of ISV patterning based on the *plexinD1* (PlxD1) mutant phenotype with a modification of our WT geometry with 4 arterial venous shunts (AVS) bypassing the DLAV in the ISV network flow. The AVS are indicated by dashed lines in Fig. 9A. Plotting the RBC trajectories and flow velocity distribution in Fig. 9Ai (Movie 23) indicated the tendency of RBCs to travel along the AVS paths in our PlxnD1 network model. The resulting spatial distribution of WSS from the network mispatterning and blood shunting highlights local regions of dead flow and low WSS (indicated by red bold lines in Fig. 9Aii, Movie 24). As a result of the flow bypass and setting up of dead-flow zones in the ISV-DLAV network, only 16.4% of RBCs flowing into the ISV network enter the DLAV (Fig. 9B), therefore highlighting a considerable loss of RBC perfusion into the DLAV directly due to blood shunting by the AVS.

**Fig. 9.**
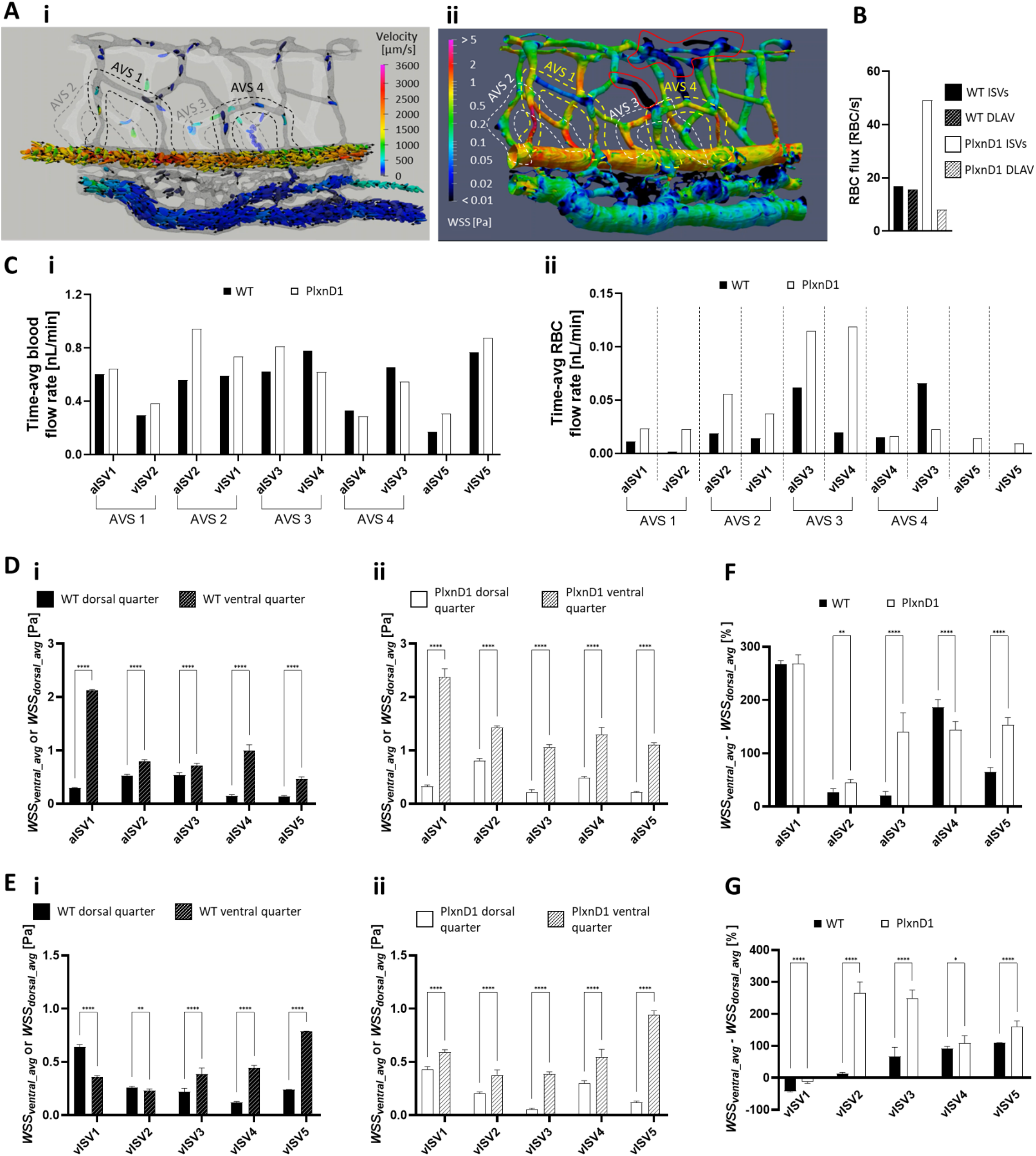
Hemodynamic alterations arising from network mispatterning in the PlxnD1 network model. **(A)** Spatial distribution map of RBC trajectories/velocities at systolic peak (i, see Movie 23 for flow oscillations) and WSS (ii, see Movie 24 for WSS oscillations) in the PlxnD1 network model. Dashed lines indicate the arterial-venous shunts that bypass DLAV connections in the ISV network and red bold lines indicate the low-WSS dead flow regions. **(B)** The flux bypass in ISV-DLAV network in PlxnD1 versus the ISV-DLAV network in WT. **(C)** Time-averaged blood (i) and RBC (ii) flow rate into ISVs amongst models. **(D)** Average systolic peak ventral (*WSS*_*ventral_avg*_) and dorsal WSS (*WSS*_*dorsal_avg*_) in aISVs for WT (i) and PlxnD1 (ii) models. **(E)***WSS*_*ventral_avg*_ and *WSS*_*dorsal_avg*_ in vISVs for WT (i) and PlxnD1 (ii) models. **(F)** Comparison of the ventral to dorsal WSS asymmetry in WT aISVs versus PlxnD1 aISVs. **(G)** Comparison of the ventral to dorsal WSS asymmetry in WT vISVs versus PlxnD1 vISVs.

There was a general rise in the blood and RBC flow rates in the PlxnD1 network due to the increased network connections of the plexus-like mispatterning. AVS 2 and AVS 3 were particularly prolific in increasing the perfusion states (Fig. 9C i and ii) of aISV2 (+68.5% for blood and +193% for RBC), aISV3 (+30.0% for blood +86.6% for RBC), vISV1 (+24.4% for blood and +161% for RBC) and vISV4 (−20.3% for blood and +507% for RBC). Next, we examined the WSS alterations arising from the mispatterning. Generally, the WSS level in the PlxnD1 ISV network was either higher or similar to the WT model in the ventral quarter of the ISVs due to the increased network flow rates. Dorsal quarter WSS in both the aISVs (Fig. 9D i and ii) and vISVs (Fig. 9E i and ii) showed similar levels for both the WT and PlxnD1. More importantly, we observed the consistent trend of higher WSS levels in ventral regions of the ISVs compared to the dorsal regions except for vISV2 in the WT network, where lumen diameter was fairly constant across the vISV2. In addition to the WSS tuning by diameter tuning (Fig. 5), we observed that the WSS gradient was further augmented by the dorsal flow bypassing in the PlxnD1 network. As shown in Fig. 9F for aISVs and Fig. 9G for vISVs, there were statistically significant increases in the WSS differences between ventral and dorsal regions for the PlxnD1 network versus WT in most ISVs except statistically significant decreases in aISV4 and vISV1.

In summary, the mispatterned network phenotype that entails blood shunting via AVS may raise the perfusion and WSS level in the modified ISV network. However, it comes at the expense of compromised perfusion to more dorsal regions of the network such as the DLAV. The resulting flow bypass instigates both perfusion and WSS asymmetry in a dorsal versus ventral stratification of levels beyond that seen in WT.

## 3. Discussion

In this work, we have incrementally improved the CFD modeling technique for examination of microhemodynamics in zebrafish microvascular networks. This was achieved by explicitly modelling the motion and viscoelastic deformation of ellipsoid capsules that represent RBCs in the blood mixture where RBCs and plasma are treated as separate phases. Doing so allowed us to consider the viscosity modulation effects of RBCs and their mechanical deformations on blood flow and lumen forces. We should however point out that inclusion of RBCs as explicit deformable particles is compute-intensive and necessitates simulating the geometrical domain of interest over a long time-scale before the multiphase mixture can equilibrate and reach developed-flow conditions. In our case, we typically ran each simulation case for at least 20 cardiac pulsation cycles which translated to 3 weeks of simulation downtime per case.

Another complexity with applying CFD models to biological data sets is the non-trivial problem of boundary conditions and their salient contribution to the overall CFD simulation result. As we demonstrated with the Marcksl1 KO flow visualization experiments, the systemic reduction of lumen diameters does not elicit simple adaptations whereby we may conveniently assume the lumen WSS to rise. Even within the mutant population, the degree of morphological alteration varies and the responding flow alteration compared to WT is also variable. These biological variations cause a fair amount of doubt about the boundary conditions and suggest that even with accurate *in silico* recreation of the 3-D lumen network geometry, the CFD simulation should consider network-specific flow boundary conditions obtained from experimental imaging. While blood velocity can be measured by particle imaging velocimetry (PIV) or particle tracking velocimetry (PTV), there is still no direct way of measuring the lumen blood pressure from in situ experiments. Our approach has been to match the velocities of various segments of the network by iteratively altering the pressure inputs defined at the anterior and posterior inlets/outlets of our network. In doing so, the pressure field and pressure gradients should naturally match the relative pressures of the physiological flow we seek to recapitulate in our model.

From the comparison of the RBC-flow model versus the no-RBC flow models, we could highlight the possible response of a microvascular network to a systemic reduction in blood viscosity. Contrary to the assumption that network WSS should fall due to the lowered viscosity, we found a counter-intuitive result of WSS maintenance from WT levels in our CFD results after matching the flow levels of our model with what we observed in flow visualization experiments with Gata1 morphants. Once again, this underlies the importance of obtaining correct boundary conditions from experiments to make the most reasonable interpretations of the physiology predicted by CFD modeling. In this particular case of hematocrit reduction, it is not clear if the flow increase after RBC removal is a response to the reduction in oxygen-carrying capacity of RBC-absent blood or if there is a homeostatic control of the set-point WSS (Baeyens et al., 2015) at specific stages of the zebrafish development. At the larval stages of development, the small size of the zebrafish allows (Kimmel et al., 1995) cutaneous delivery of oxygen via passive diffusion from the environment into tissue (Hughes and Perry, 2021). Hence, it appears quite unlikely that the organism is adapting its flow to compensate for losses to oxygen carrying capacity of the RBC-absent blood. We thus lean towards the opinion that the physiological adaptation to hematocrit reduction is possibly a biosystems optimization towards a set-point WSS relative to the organism’s developmental age.

In our examination of the flow partitioning at ISV/CA bifurcations, we observed that the RBC phase does not follow the same partitioning tendency as the overall blood mixture. RBC entrance into ISVs appears to be more sensitive to lumen diameters in the ventral regions of the aISVs rather than the overall pressure gradient, especially when the lumen diameters approach calibers below 5 μm. We observed this determinism in both the flow visualization experiments with Marcksl1 KO embryos and our 3 CFD models of the Marcksl1 KO network. In the experiment we saw that the RBC perfusion level into the ISV network dramatically cuts to zero between the transitional diameters 3 – 4 μm (Fig. 7Ev). Supporting this, our CFD model employed three driving arterial pressure levels from high to low and we saw no significant difference in the ISV RBC perfusion levels between the high to low pressure cases (Fig. 7G).

Finally, as mentioned earlier, the lumen blood pressure in situ is notoriously difficult to obtain from biological samples. Based on careful cross validation of other hemodynamic parameters such as RBC flow velocity, hematocrit, heart rate and even vessel wall motions in a cardiac cycle, CFD techniques could reasonably provide spatiotemporal maps of pressure in addition to WSS. These two forces together may provide a more holistic discussion of mechanotransduction and signaling cues for angiogenesis and vascular remodeling in the microvascular network. Our future plan is to progress this methodology and perform studies linking experimentally-obtained perfusion dynamics and CFD-obtained flow forces to distinct vascular morphogenetic processes in individual developing zebrafish under long-term imaging.

## 4. Models and Methods

### 4.1 Zebrafish handling and experiments

Zebrafish (*Danio rerio*) were raised and staged according to established protocols (Charles B. Kimmel et al., 1995). Zebrafish lines used were *Tg(kdrl:EGFP)^s843^*(Jin et al., 2005), *Tg(gata1:dsRed)^sd2^* (Traver et al., 2003) and *marcksl1a^rk23^;marcksl1b^rk24^* (Kondrychyn et al., 2020). For microangiography experiments, 1 nl dextran tetramethyl rhodamine (MW = 2000 kDa, Invitrogen) at 10mg/ml was injected into the Duct of Cuvier of 2 dpf zebrafish. Depletion of RBC formation was achieved by injecting 1 nl of 0.1mM Gata1 morpholino into 1- to 2-cell stage embryos.

For imaging experiments, embryos were embedded in 0.8% low-melting agarose in E3 medium containing 0.16 mg/ml Tricaine. To obtain 3D images of blood vascular network, confocal z-stacks were acquired using a 40X 1.25NA objective lens on an inverted Olympus IX83/CSU-Wi spinning disc confocal microscope (Yokogawa) with a Zyla 4.2 CMOS camera (Andor). To image RBC flow, 1000 frames at 180 fps were captured using a sCMOS camera (PCO, pco.edge 4.2 CL) mounted on a florescent stereomicroscope (Leica, M205FA). Images were processed using Fiji (NIH).

### 4.2 Zebrafish trunk blood flow analysis

In this project, we have developed a method for analyzing luminal pressures and wall shear stresses in the zebrafish trunk circulation using experimental images of the network morphology and computational representation of the physiological flow patterning that results from this morphology. To do this, we introduce fluorescent heavy molecular weight dextran into the blood vessels through microangiography. The dextran serves as a contrast agent for obtaining lumen morphology in confocal microscopy. Using the three-dimensional reconstruction of the confocal images, we obtain an *in-silico* model of the zebrafish trunk network that is analyzed with computational fluid dynamics (CFD).

### 4.3 Computational fluid dynamics

There are three major solver components to our blood flow model for zebrafish trunk circulation analysis. **1)** The deformation of the red blood cell (RBC) membrane by flow forces and internal forces is calculated via the coarse-grained spectrin model (CGSM). **2)** The movement of plasma and cytoplasmic fluid is calculated by the Lattice Boltzmann method (LBM). **3)** The fluid-structure interaction (FSI) solver that calculates the exchange of momentum between the RBC membrane and the surrounding fluid follows the immersed boundary method (IBM).

#### 4.3.1 RBC deformation model: Coarse-grained spectrin model (CGSM)

The zebrafish RBC is modelled as a flattened ellipsoid capsule with an RBC membrane shell enveloping a cytosolic interior and a nucleus (Fig. 1C). The membrane is represented by a hexagonal mesh where edges represent the cytoskeleton (CSK) of spectrin chains joined to actin filaments (Boal, 1994; Pivkin and Karniadakis, 2008) and the triangle surface elements represent the plasma membrane (PM). Likewise, the nucleus is defined as a three-dimensional meshwork where enjoining edges act as compressible springs for the prescription of nuclear deformation. Tension in the RBC membrane arising from deformation consists of the elastic component and the viscous component:

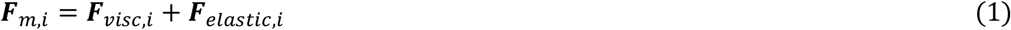

where ***F***_*visc,i*_ is the viscous force and ***F***_*elastic,i*_ is the elastic force at membrane vertex *i*. The elastic force may be obtained by differentiating the local elastic energy (*U*_*elastic,i*_) with respect to the nodal displacement (***s_i_***) produced by the deformation:

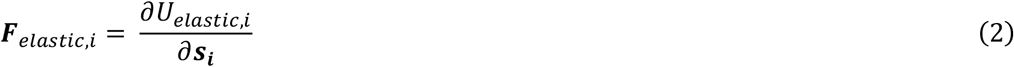

The elastic constitution of the RBC membrane can be decomposed to the individual mechanical aspects of the PM and CSK:

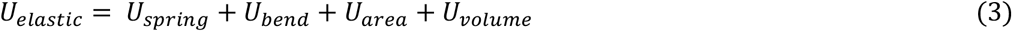

##### Shear elasticity of the CSK and stiffness of the RBC nucleus

Shearing properties of the CSK were modelled using the worm-like chain (WLC) extension model and the power-law (POW) compression model (Fedosov et al., 2010a):

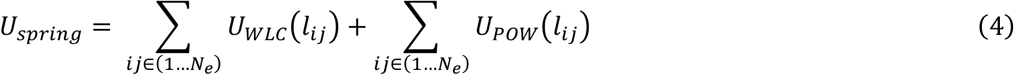

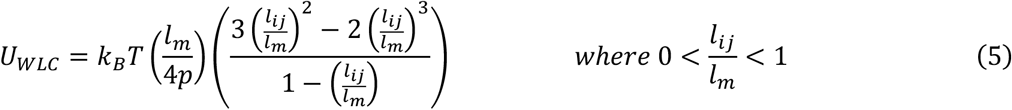

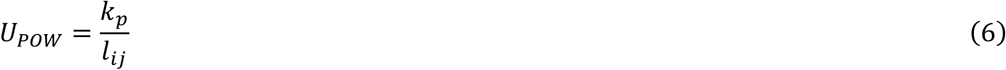

where *l_ij_* is the mesh edge length between vertices *i* and *j*, *l_m_* is the maximum allowable edge length in the WLC model and *p* is the persistence length. *k*_*B*_*T* is the energy per unit mass which may be considered as a membrane constant for an isothermal consideration. *k_p_* is the spring constant for the POW compression spring model. The resting shear modulus (*E*_*s*0_) of the WLC-POW network can be given by the following relation(Fedosov et al., 2010b):

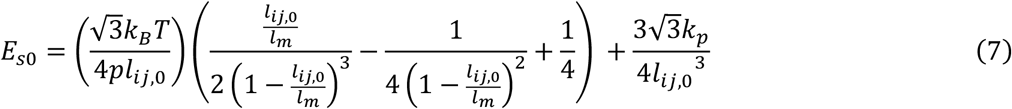

where *l*_*ii*,0_ is the mesh edge length for the RBC at rest.

Using the WLC-POW springs we are able to exhibit the strain hardening behavior of RBC membranes (see Fig. 10A). We employed the same WLC-POW spring elements for the discretized network of springs that model the stiffness of the RBC nucleus.

**Fig. 10.**
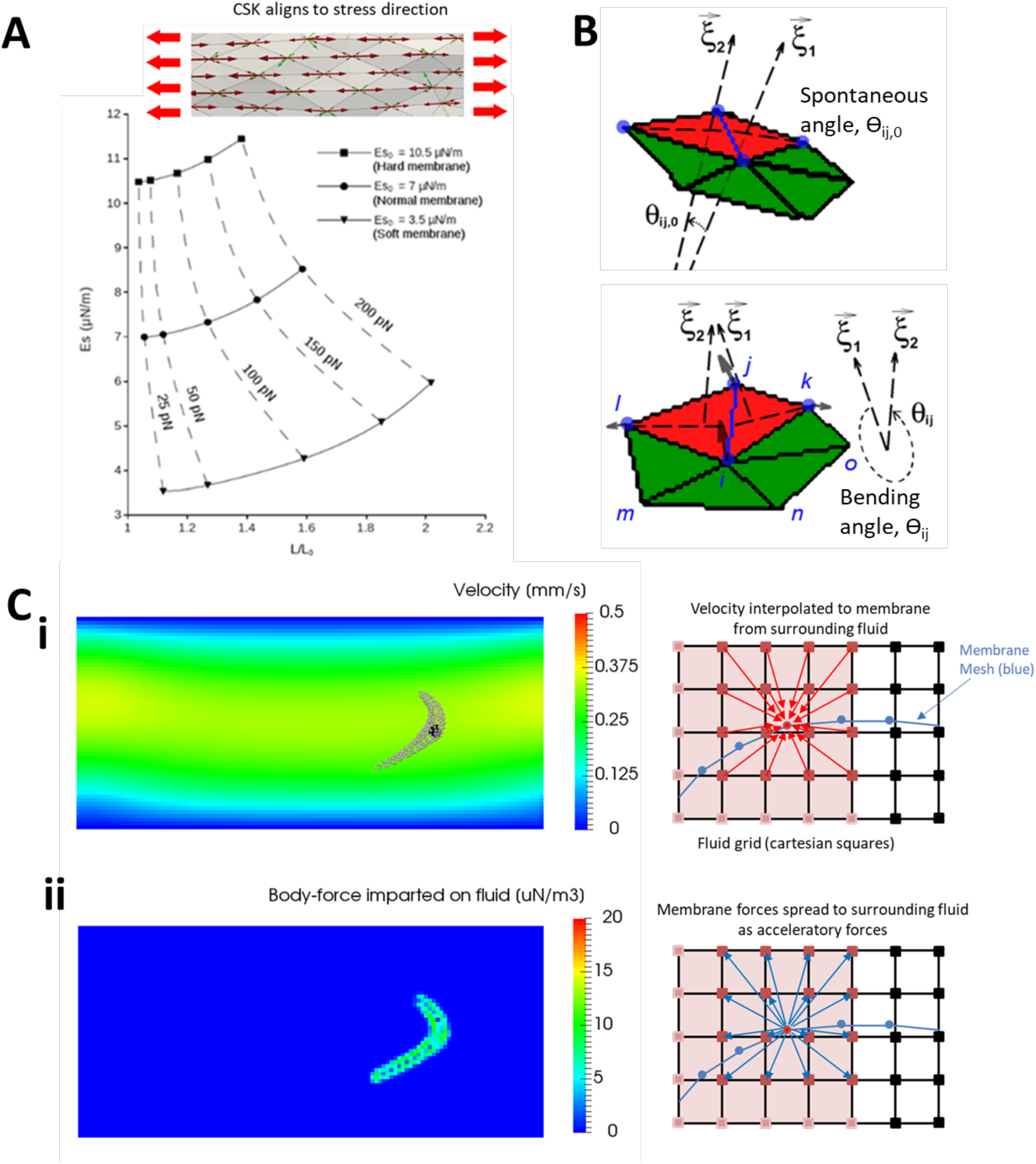
The fundamental components and physics of the cell and plasma phase CFD model. **(A)** Elastic strain-hardening behavior of the WLC-POW network within the physiological range of shear elastic modulus (*E*_*s*0_ = 3.5, 7 and 10.5 μN/m). **(B)** Schematics of the bending angle determination for calculating bending energy in the plasma membrane. **(C)** Schematics of the immersed boundary method where membrane velocities are interpolated from the surrounding fluid velocities (solved by LBM) in accordance with eq. 19 & 20 and forces of deformation in the membrane (solved by CGSM) are spread to the fluid as body-forces (accelerations) to update fluid velocities in the LBM in the next time step (ii).

##### Bending elasticity

The bending energy in the mesh network was calculated using the angle *θ_ij_* subtended by the normals (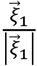 and 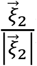) of the two adjoining triangular mesh elements along their shared mesh edge *ij* (see Fig. 10B):

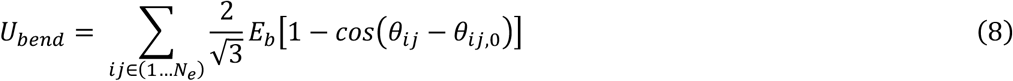

where *E_b_* is the bending modulus, *θ*_*ij*,0_ is the local spontaneous angle that a free sheet of spectrin polymers will adopt in their unstressed state. Since we assume the RBC rest state to be the reference energy state in our model, *θ*_*ij*,0_ is the angle between triangular mesh elements for the RBC at rest.

##### Area elasticity

The PM has been reported to be highly incompressible and area dilations exceeding 2-4 % produce immediate lysis (Evans, 1989; Evans et al., 1976). Incompressibility of the PM was prescribed by the area penalty model:

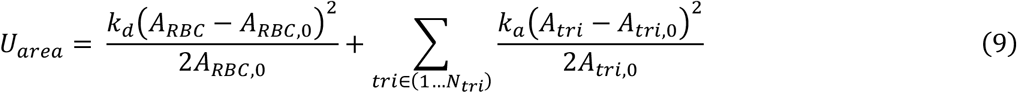

where the area deformation energy is a combination of both global and local surface area changes. *A*_*tri*_ is the current area of the local triangular mesh element and *A*_*tri*,0_ is the area of the element when the RBC is at rest. *A_RBC_* is the current surface area of the RBC membrane and *A*_*RBC*,0_ is the membrane surface area of the RBC at rest. *k_d_* represents the global area compressibility coefficient while *k_a_* is the local compressibility coefficient. The compressibility coefficients may be related to the area compressibility modulus by the following relation (Fedosov et al., 2010b):

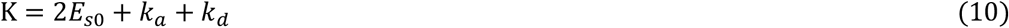

As Κ (0.432 N/m) is much larger than *E*_*s*0_ (7 μN/m), it is practically determined by the *k_a_* + *k_d_* contribution in the model. For simplicity in the model, *k_a_* and *k_d_* were set to equal values.

##### Volume constraint

The volume incompressibility is enforced to ensure the RBC volume is conserved. This is achieved by an isotropic penalty term similar to the area incompressibility energy function:

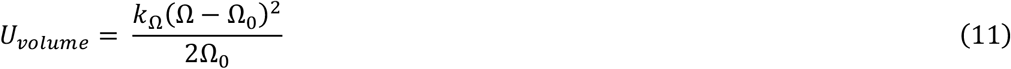

where Ω is the current cytoplasmic volume of the RBC, Ω_0_ is the original cytoplasmic volume based on the mean corpuscular volume (MCV) of 100 fL and *k*_Ω_ is the volume correction penalty coefficient set as 220 N/m^3^.

##### Membrane viscosity

The rate of surface deformation in the membrane is regulated by membrane viscosity. We follow the dissipative spring model proposed by (Español, 1998):

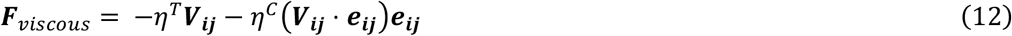

where ***V***_***ij***_ is the relative velocity between mesh vertices *i* and *j*, ***e***_***ij***_ is the unit displacement vector between *i* and *j*, *η^T^* and *η^C^* are dissipative coefficients related to the surface viscosity (*η_m_*) of the membrane by the following relation:

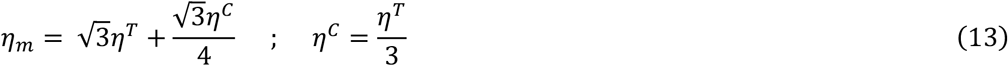

#### 4.3.2 Plasma and cytosol transport model: Lattice Boltzmann Method (LBM)

The LBM solves the Boltzmann equation numerically by discretizing the mass and momentum of fluid particles on a fixed computational lattice. In the LBM, the behavior of the macro-system (defined by pressure, viscosity and density) is given by the kinetics and sum of its microstates. Microstates are defined in the discrete packets by the density distribution function (*f_i_*(***x***)) and the discretized transport equation involves the streaming and collision of *f_i_*(***x***) in the lattice. In the streaming process, 19 discrete streaming directions and lattice velocities are defined as follows (Rosis and Coreixas, 2020):

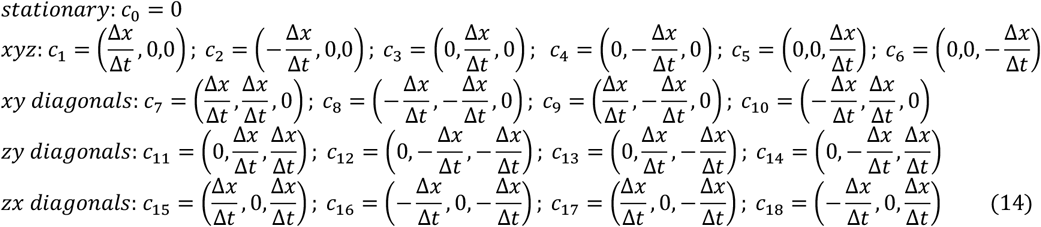

where Δ*x* is the space step size in the lattice, Δ*t* is the time step size, and symbol *i* denotes the flux direction index. The LB equation with a general force term can be expressed as (Guo et al., 2002):

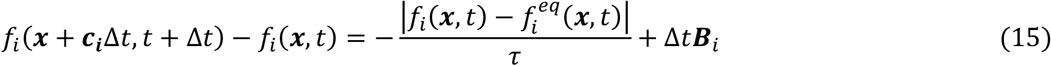

where ***B***_*i*_ is the body force term represented on the microstate and *τ* is a relaxation parameter towards the equilibrium distribution 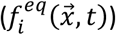 which can be obtained through:

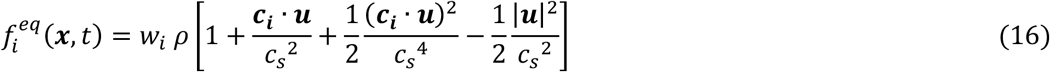

where *ρ* is the fluid density and ***u*** is the velocity; the two are calculated by the equivalency of the macroscopic state to the sum of microstates: *ρ* = ∑_*i*_ *f_i_*, ***u*** = ∑_*i*_ *f_i_* ***c***_***i***_/*ρ*, and 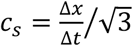 (speed of sound in the lattice). The weights, *w_i_*, used in Eq. (16) are given by 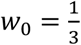, 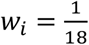 for *i* = 1 − 6 and 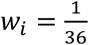 for *i* = 7 − 18.

The microstate body force term in Eq. (16) is given by:

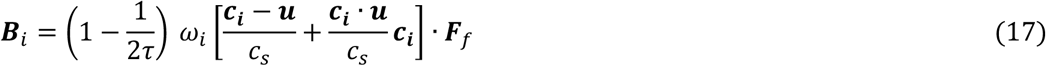

where ***F***_*f*_ is the fluid-structure interaction body force acting on the fluid from the IBM coupling with the RBC membrane solver. Proof of the equivalency between the LBM discretization and the continuum Navier-Stokes equations has been given through the Chapman-Enskog expansion of the LB equation (Li, 2015) The pressure (*P*) and kinematic viscosity (*ν*) of the fluid is represented in the LBM system by:

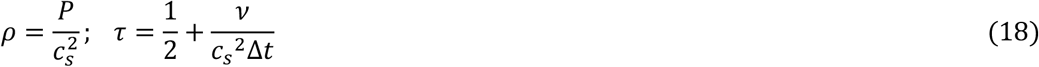

#### 4.3.3 Fluid structure interaction: Immersed boundary method (IBM)

The IBM introduced by (Peskin, 1977) enforces the fluid-structure interaction in the simulation by providing the bidirectional coupling between the fluid motion and the membrane dynamics. Velocity interpolations onto the membrane surface grid from the LBM-derived velocity field (Fig. 10Ci) using the discrete delta function are performed to update the trajectories of the RBC membrane nodes as follows:

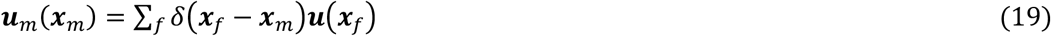

where *δ*(***x***_*f*_ − ***x***_*m*_) is the discrete delta function that is given by:

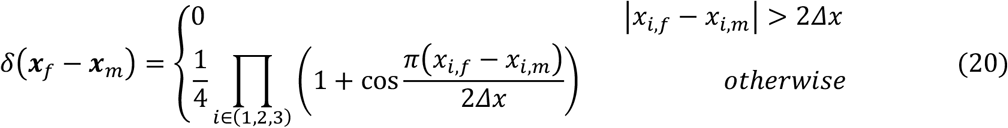

where *x_i_* = {*x*_1_, *x*_2_, *x*_3_} = *x*, *y*, *z*

The viscoelastic resistance of the RBC membrane against hydrodynamic forces and its effect on the surrounding fluid is similarly enforced whereby the membrane force ***F***_*m*_(***x***_*m*_) arising from the membrane deformation at membrane coordinates ***x***_*m*_ is distributed to the surrounding fluid nodes (coordinates ***x***_*f*_) (Fig. 10Cii) as a body force ***F***_*f*_(***x***_*f*_) using the discrete delta function:

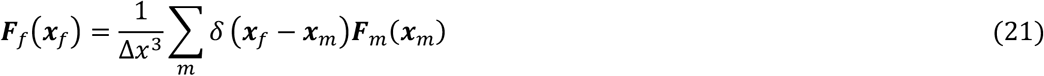

The force density spreading and membrane node velocity interpolation were performed on a 4*Δx* × 4*Δx* × 4*Δx* region.

##### Inter-RBC and RBC to lumen wall collisions

In order to preserve the robustness and numerical integrity of RBC motions in the simulation while maintaining a larger time step for efficient computational downtime, we had to enforce explicit control of inter-surface contact. We prevented the interpenetration of RBC and lumen wall mesh by enforcing a reflection velocity correction (***u***_*m_correc*_) for a membrane node when the separation distance (*r_sep_*) between the membrane node and a neighboring membrane or lumen wall surface was < 0.2 μm:

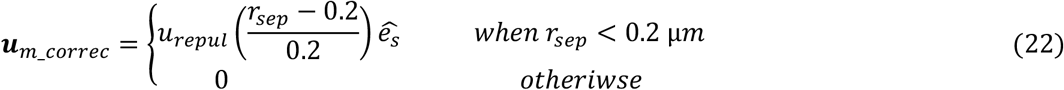

Where *u_repul_* is the coefficient for velocity reflection where we found best results using 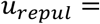 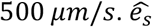 gives the direction of the velocity reflection, which is taken in the direction of the surface normal vector of the neighboring surface. Note that in Eqn. 22 all distances are to be taken in μm. The velocity of the local membrane node ***u***_*m*_(***x***_*m*_) evaluated in eqn 19 is modified by addition of ***u***_*m_correc*_ for the final velocity to update the trajectory of the RBC mesh node in the next time step:

Because we applied explicit correction of the RBC mesh node velocity, we have to impart this momentum to the surrounding fluid and this is done by modifying ***F***_*m*_(***x***_*m*_) in eqn. 21 with the addition of the force correction term ***F***_*m_correc*_:

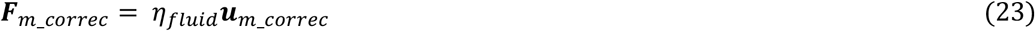

Where *η_fluid_* represents a fluid dissipative friction to velocity corrections, for best results we used the value *η_fluid_* = 1 × 10^−8^ N·s/m.

## Acknowledgements

We thank members of the Phng Lab and Satoru Okuda for discussions and RIKEN BDR Aquatic Facility for zebrafish care. LKP is supported by RIKEN BDR core funding and JSPS KAKENHI (22H02624 and 22H05168). SSMY is supported by RIKEN Special Postdoctoral Researcher Program and JSPS KAKENHI (JP20K20190).

## Movie Legends

**Movie 1**: Inter-cellular collisions between deformable red blood cells (RBCs) and RBC-vessel-wall collisions can produce extensive deformation of impinged RBCs and cause flow passage reductions. Utilization of short-range repulsion forces mediated contact mechanics and ensured a robust numerical solver.

**Movie 2**: Comparison of blood flow velocity in a sagittal cross section along the caudal artery axis. From left to right: model 1 – the wild-type (WT) geometry with RBCs included in the flow model, model 2 – the RBC-absent flow model that maintains same pressure as model 1, model 3 – the RBC-absent flow model that maintains same blood flow rate as model 1.

**Movie 3**: Comparison of the spatial distribution of lumen blood pressure. From left to right: model 1 – the WT geometry with RBCs included in the flow model, model 2 – the RBC-absent flow model that maintains same pressure as model 1, model 3 – the RBC-absent flow model that maintains same blood flow rate as model 1.

**Movie 4**: Comparison of the spatial distribution of lumen wall shear stress (WSS). From left to right: model 1 – the WT geometry with RBCs included in the flow model, model 2 – the RBC-absent flow model that maintains same pressure as model 1, model 3 – the RBC-absent flow model that maintains same blood flow rate as model 1.

**Movie 5**: Oscillating RBC flow velocities in the WT network.

**Movie 6**: Oscillating RBC flow velocities in the smooth geometry model (SGM) network.

**Movie 7**: Oscillating spatial distribution of blood pressure in the WT network.

**Movie 8**: Oscillating spatial distribution of blood pressure in the SGM network.

**Movie 9**: Oscillating spatial distribution of WSS in the WT network.

**Movie 10**: Oscillating spatial distribution of WSS in the SGM network.

**Movie 11**: Oscillating RBC flow velocities in the Marckl1 over-expression phenotype (Marcksl1 OE) network.

**Movie 12**: Oscillating spatial distribution of blood pressure in Marcksl1 OE network.

**Movie 13**: Oscillating spatial distribution of WSS in Marcksl1 OE network.

**Movie 14**: Oscillating RBC flow velocities in the *marckl1a;marckl1b* double knockout phenotype (Marcksl1 KO) network with 22% reduced arterial pressure compared to WT (model 1).

**Movie 15**: Oscillating spatial distribution of blood pressure in the Marcksl1 KO network with 22% reduced arterial pressure compared to WT (model 1).

**Movie 16**: Oscillating spatial distribution of WSS in the Marcksl1 KO network with 22% reduced arterial pressure compared to WT (model 1).

**Movie 17**: Oscillating RBC flow velocities in the Marcksl1 KO network with same arterial pressure compared to WT (model 2).

**Movie 18**: Oscillating spatial distribution of blood pressure in the Marcksl1 KO network with same arterial pressure compared to WT (model 2).

**Movie 19**: Oscillating spatial distribution of WSS in the Marcksl1 KO network with same arterial pressure compared to WT (model 2).

**Movie 20**: Oscillating RBC flow velocities in the Marcksl1 KO network with 17% increased arterial pressure compared to WT (model 3).

**Movie 21**: Oscillating spatial distribution of blood pressure in the Marcksl1 KO network with 17% increased arterial pressure compared to WT (model 3).

**Movie 22**: Oscillating spatial distribution of WSS in the Marcksl1 KO network with 17% increased arterial pressure compared to WT (model 3).

**Movie 23**: Oscillating RBC flow velocities in the PlexinD1 knockout (PlxnD1) network.

**Movie 24**: Oscillating spatial distribution of WSS in the PlxnD1 knockout network.

## References

Alizadehrad D, Imai Y, Nakaaki K, Ishikawa T, Yamaguchi T. 2012. Quantification of red blood cell deformation at high-hematocrit blood flow in microvessels. J Biomech 45:2684–2689. doi:10.1016/j.jbiomech.2012.08.026

An D, Yu P, Freund KB, Yu D-Y, Balaratnasingam C. 2020. Three-Dimensional Characterization of the Normal Human Parafoveal Microvasculature Using Structural Criteria and High-Resolution Confocal Microscopy. Invest Ophth Vis Sci 61:3. doi:10.1167/iovs.61.10.3

Baeyens N, Nicoli S, Coon BG, Ross TD, Dries KV den, Han J, Lauridsen HM, Mejean CO, Eichmann A, Thomas J-L, Humphrey JD, Schwartz MA. 2015. Vascular remodeling is governed by a VEGFR3-dependent fluid shear stress set point. Elife 4:e04645. doi:10.7554/elife.04645

Balogh P, Bagchi P. 2019. Three-dimensional distribution of wall shear stress and its gradient in red cell-resolved computational modeling of blood flow in in vivo-like microvascular networks. Physiological Reports 7:e14067. doi:10.14814/phy2.14067

Boal DH. 1994. Computer simulation of a model network for the erythrocyte cytoskeleton. Biophys J 67:521–529. doi:10.1016/s0006-3495(94)80511-9

Cesarovic N, Lipiski M, Falk V, Emmert MY. 2020. Animals in cardiovascular research. Eur Heart J 41:200–203. doi:10.1093/eurheartj/ehz933

Chen Q, Jiang L, Li C, Hu D, Bu J, Cai D, Du J. 2012. Haemodynamics-Driven Developmental Pruning of Brain Vasculature in Zebrafish. Plos Biol 10:e1001374. doi:10.1371/journal.pbio.1001374

Djukic TR, Karthik S, Saveljic I, Djonov V, Filipovic N. 2016. Modeling the Behavior of Red Blood Cells within the Caudal Vein Plexus of Zebrafish. Front Physiol 7:455. doi:10.3389/fphys.2016.00455

Doddi SK, Bagchi P. 2009. Three-dimensional computational modeling of multiple deformable cells flowing in microvessels. Phys Rev E 79:046318. doi:10.1103/physreve.79.046318

Dupin MM, Halliday I, Care CM, Alboul L, Munn LL. 2007. Modeling the flow of dense suspensions of deformable particles in three dimensions. Phys Rev E 75:066707. doi:10.1103/physreve.75.066707

Enjalbert R, Hardman D, Krüger T, Bernabeu MO. 2021. Compressed vessels bias red blood cell partitioning at bifurcations in a hematocrit-dependent manner: Implications in tumor blood flow. Proc National Acad Sci 118:e2025236118. doi:10.1073/pnas.2025236118

Español P. 1998. Fluid particle model. Phys Rev E 57:2930–2948. doi:10.1103/physreve.57.2930

Evans EA. 1989. [1] Structure and deformation properties of red blood cells: Concepts and quantitative methods. Methods Enzymol 173:3–35. doi:10.1016/s0076-6879(89)73003-2

Evans EA, Waugh R, Melnik L. 1976. Elastic area compressibility modulus of red cell membrane. Biophys J 16:585–595. doi:10.1016/s0006-3495(76)85713-x

Fedosov DA, Caswell B, Karniadakis GE. 2010a. A Multiscale Red Blood Cell Model with Accurate Mechanics, Rheology, and Dynamics. Biophys J 98:2215–2225. doi:10.1016/j.bpj.2010.02.002

Fedosov DA, Caswell B, Popel AS, Karniadakis GE. 2010b. Blood Flow and Cell-Free Layer in Microvessels. Microcirculation 17:615–628. doi:10.1111/j.1549-8719.2010.00056.x

Fedosov DA, Pan W, Caswell B, Gompper G, Karniadakis GE. 2011. Predicting human blood viscosity in silico. Proc National Acad Sci 108:11772–11777. doi:10.1073/pnas.1101210108

Gebala V, Collins R, Geudens I, Phng L-K, Gerhardt H. 2016. Blood flow drives lumen formation by inverse membrane blebbing during angiogenesis in vivo. Nat Cell Biol 18:443–450. doi:10.1038/ncb3320

Geudens I, Coxam B, Alt S, Gebala V, Vion A-C, Meier K, Rosa A, Gerhardt H. 2019. Artery-vein specification in the zebrafish trunk is pre-patterned by heterogeneous Notch activity and balanced by flow-mediated fine-tuning. Development 146:dev181024. doi:10.1242/dev.181024

Guo Z, Zheng C, Shi B. 2002. Discrete lattice effects on the forcing term in the lattice Boltzmann method. Phys Rev E 65:046308. doi:10.1103/physreve.65.046308

Harlepp S, Thalmann F, Follain G, Goetz JG. 2017. Hemodynamic forces can be accurately measured in vivo with optical tweezers. Mol Biol Cell 28:3252–3260. doi:10.1091/mbc.e17-06-0382

Hogan BM, Schulte-Merker S. 2017. How to Plumb a Pisces: Understanding Vascular Development and Disease Using Zebrafish Embryos. Dev Cell 42:567–583. doi:10.1016/j.devcel.2017.08.015

Hughes MC, Perry SF. 2021. Does blood flow limit acute hypoxia performance in larval zebrafish (Danio rerio)? J Comp Physiol B 191:469–478. doi:10.1007/s00360-020-01331-z

Jährling N, Becker K, Dodt H-U. 2009. 3D-reconstruction of blood vessels by ultramicroscopy. Organogenesis 5:227–230. doi:10.4161/org.5.4.10403

Jia T, Wang C, Han Z, Wang X, Ding M, Wang Q. 2020. Experimental Rodent Models of Cardiovascular Diseases. Frontiers Cardiovasc Medicine 7:588075. doi:10.3389/fcvm.2020.588075

Jin S-W, Beis D, Mitchell T, Chen J-N, Stainier DYR. 2005. Cellular and molecular analyses of vascular tube and lumen formation in zebrafish. Development 132:5199–5209. doi:10.1242/dev.02087

Karthik S, Djukic T, Kim J-D, Zuber B, Makanya A, Odriozola A, Hlushchuk R, Filipovic N, Jin SW, Djonov V. 2018. Synergistic interaction of sprouting and intussusceptive angiogenesis during zebrafish caudal vein plexus development. Sci Rep-uk 8:9840. doi:10.1038/s41598-018-27791-6

Kimmel C B, Ballard WW, Kimmel SR, Ullmann B, Schilling TF. 1995. Stages of embryonic development of the zebrafish. Developmental dynamics : an official publication of the American Association of Anatomists 203:253--310.

Kimmel Charles B., Ballard WW, Kimmel SR, Ullmann B, Schilling TF. 1995. Stages of embryonic development of the zebrafish. Dev Dyn 203:253–310. doi:10.1002/aja.1002030302

Kochhan E, Lenard A, Ellertsdottir E, Herwig L, Affolter M, Belting H-G, Siekmann AF. 2013. Blood Flow Changes Coincide with Cellular Rearrangements during Blood Vessel Pruning in Zebrafish Embryos. Plos One 8:e75060. doi:10.1371/journal.pone.0075060

Kondrychyn I, Kelly DJ, Carretero NT, Nomori A, Kato K, Chong J, Nakajima H, Okuda S, Mochizuki N, Phng L-K. 2020. Marcksl1 modulates endothelial cell mechanoresponse to haemodynamic forces to control blood vessel shape and size. Nat Commun 11:5476. doi:10.1038/s41467-020-19308-5

Lanotte L, Mauer J, Mendez S, Fedosov DA, Fromental J-M, Claveria V, Nicoud F, Gompper G, Abkarian M. 2016. Red cells’ dynamic morphologies govern blood shear thinning under microcirculatory flow conditions. Proc National Acad Sci 113:13289–13294. doi:10.1073/pnas.1608074113

Lee J, Fei P, Packard RRS, Kang H, Xu H, Baek KI, Jen N, Chen J, Yen H, Kuo C-CJ, Chi NC, Ho C-M, Li R, Hsiai TK. 2016. 4-Dimensional light-sheet microscopy to elucidate shear stress modulation of cardiac trabeculation. J Clin Invest 126:3158–3158. doi:10.1172/jci89549

Lee J, Vedula V, Baek KI, Chen J, Hsu JJ, Ding Y, Chang C-C, Kang H, Small A, Fei P, Chuong C, Li R, Demer L, Packard RRS, Marsden AL, Hsiai TK. 2018. Spatial and temporal variations in hemodynamic forces initiate cardiac trabeculation. Jci Insight 3:e96672. doi:10.1172/jci.insight.96672

Li J. 2015. Appendix: Chapman-Enskog Expansion in the Lattice Boltzmann Method. Arxiv. doi:10.48550/arxiv.1512.02599

Malone MH, Sciaky N, Stalheim L, Hahn KM, Linney E, Johnson GL. 2007. Laser-scanning velocimetry: A confocal microscopy method for quantitative measurement of cardiovascular performance in zebrafish embryos and larvae. Bmc Biotechnol 7:40–40. doi:10.1186/1472-6750-7-40

Peskin CS. 1977. Numerical analysis of blood flow in the heart. J Comput Phys 25:220–252. doi:10.1016/0021-9991(77)90100-0

Pivkin IV, Karniadakis GE. 2008. Accurate Coarse-Grained Modeling of Red Blood Cells. Phys Rev Lett 101:118105. doi:10.1103/physrevlett.101.118105

Pries AR, Secomb TW, Gaehtgens P, Gross JF. 1990. Blood flow in microvascular networks. Experiments and simulation. Circ Res 67:826–834. doi:10.1161/01.res.67.4.826

Ransom DG, Haffter P, Odenthal J, Brownlie A, Vogelsang E, Kelsh RN, Brand M, Eeden FJ van, Furutani-Seiki M, Granato M, Hammerschmidt M, Heisenberg CP, Jiang YJ, Kane DA, Mullins MC, Nüsslein-Volhard C. 1996. Characterization of zebrafish mutants with defects in embryonic hematopoiesis. Dev Camb Engl 123:311–9. doi:10.1242/dev.123.1.311

Rosis AD, Coreixas C. 2020. Multiphysics flow simulations using D3Q19 lattice Boltzmann methods based on central moments. Phys Fluids 32:117101. doi:10.1063/5.0026316

Roustaei M, Baek KI, Wang Z, Cavallero S, Satta S, Lai A, O’Donnell R, Vedula V, Ding Y, Marsden AL, Hsiai TK. 2022. Computational simulations of the 4D micro-circulatory network in zebrafish tail amputation and regeneration. J Roy Soc Interface 19:20210898. doi:10.1098/rsif.2021.0898

Salman HE, Yalcin HC. 2021. Computational Modeling of Blood Flow Hemodynamics for Biomechanical Investigation of Cardiac Development and Disease. J Cardiovasc Dev Dis 8:14. doi:10.3390/jcdd8020014

Simmers MB, Pryor AW, Blackman BR. 2007. Arterial shear stress regulates endothelial cell-directed migration, polarity, and morphology in confluent monolayers. Am J Physiol-heart C 293:H1937–H1946. doi:10.1152/ajpheart.00534.2007

Tardy Y, Resnick N, Nagel T, Jr MAG, Jr CFD. 1997. Shear Stress Gradients Remodel Endothelial Monolayers in Vitro via a Cell Proliferation-Migration-Loss Cycle. Arteriosclerosis Thrombosis Vasc Biology 17:3102–3106. doi:10.1161/01.atv.17.11.3102

Tinevez J-Y, Perry N, Schindelin J, Hoopes GM, Reynolds GD, Laplantine E, Bednarek SY, Shorte SL, Eliceiri KW. 2017. TrackMate: An open and extensible platform for single-particle tracking. Methods 115:80–90. doi:10.1016/j.ymeth.2016.09.016

Torres-Vázquez J, Gitler AD, Fraser SD, Berk JD, Pham VN, Fishman MC, Childs S, Epstein JA, Weinstein BM. 2004. Semaphorin-Plexin Signaling Guides Patterning of the Developing Vasculature. Dev Cell 7:117–123. doi:10.1016/j.devcel.2004.06.008

Traver D, Paw BH, Poss KD, Penberthy WT, Lin S, Zon LI. 2003. Transplantation and in vivo imaging of multilineage engraftment in zebrafish bloodless mutants. Nat Immunol 4:1238–1246. doi:10.1038/ni1007

Vahidkhah K, Diamond SL, Bagchi P. 2014. Platelet Dynamics in Three-Dimensional Simulation of Whole Blood. Biophys J 106:2529–2540. doi:10.1016/j.bpj.2014.04.028

Vedula V, Lee J, Xu H, Kuo C-CJ, Hsiai TK, Marsden AL. 2017. A method to quantify mechanobiologic forces during zebrafish cardiac development using 4-D light sheet imaging and computational modeling. Plos Comput Biol 13:e1005828. doi:10.1371/journal.pcbi.1005828

Vermot J, Forouhar AS, Liebling M, Wu D, Plummer D, Gharib M, Fraser SE. 2009. Reversing Blood Flows Act through klf2a to Ensure Normal Valvulogenesis in the Developing Heart. Plos Biol 7:e1000246. doi:10.1371/journal.pbio.1000246

Weijts B, Gutierrez E, Saikin SK, Ablooglu AJ, Traver D, Groisman A, Tkachenko E. 2018. Blood flow-induced Notch activation and endothelial migration enable vascular remodeling in zebrafish embryos. Nat Commun 9:5314. doi:10.1038/s41467-018-07732-7

Ye SS, Ju M, Kim S. 2016. Recovery of cell-free layer and wall shear stress profile symmetry downstream of an arteriolar bifurcation. Microvasc Res 106:14–23. doi:10.1016/j.mvr.2016.03.003

Ye SSM, Kim JK, Carretero NT, Phng L-K. 2022. High-Throughput Imaging of Blood Flow Reveals Developmental Changes in Distribution Patterns of Hemodynamic Quantities in Developing Zebrafish. Front Physiol 13:881929. doi:10.3389/fphys.2022.881929

Zhou Q, Perovic T, Fechner I, Edgar LT, Hoskins PR, Gerhardt H, Krüger T, Bernabeu MO. 2021. Association between erythrocyte dynamics and vessel remodelling in developmental vascular networks. J Roy Soc Interface 18:20210113. doi:10.1098/rsif.2021.0113

Zygmunt T, Trzaska S, Edelstein L, Walls J, Rajamani S, Gale N, Daroles L, Ramírez C, Ulrich F, Torres-Vázquez J. 2012. ‘In parallel’ interconnectivity of the dorsal longitudinal anastomotic vessels requires both VEGF signaling and circulatory flow. J Cell Sci 125:5159–5167. doi:10.1242/jcs.108555

