## Supplementary material for "A cell-and-plasma numerical model reveals hemodynamic stress and flow adaptation in zebrafish microvessels after morphological alteration"

### Flow visualization experiments for CFD validation and CFD boundary conditions

We performed particle velocimetry tracking of dsred-gata1 RBCs to:

1) Obtain blood flow velocities in the caudal artery (CA), caudal vein (CV) and intersegmental vessels (ISVs) pertaining to the domain simulated in our CFD model.
2) Obtain RBC perfusion levels in the ISV network by counting total number of RBCs flowing in the ISVs and then divided by the number of ISVs in the imaged domain (usually 10 ISVs based on the magnification settings of the region of interest).

### Section A:

#### Gata1 Morpholino injected Zebrafish, a hematocrit reduction experiment and perfusion quantification experiment

### MO Gata1 #1: HR = 174 bpm; Moderate hematocrit

CA 29 beats in 10 seconds

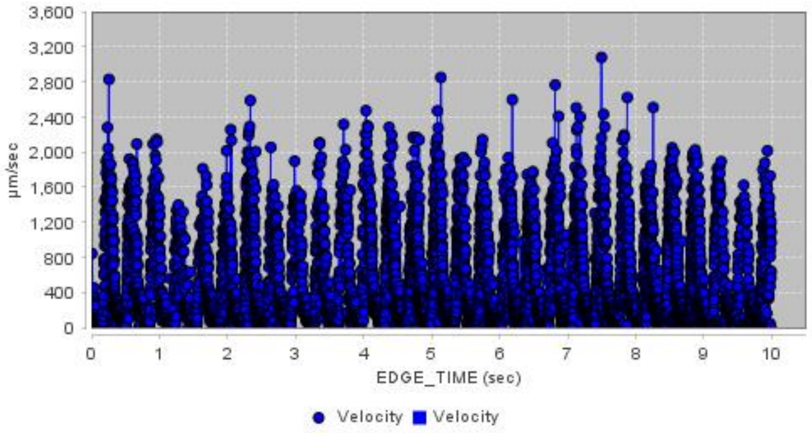


CV

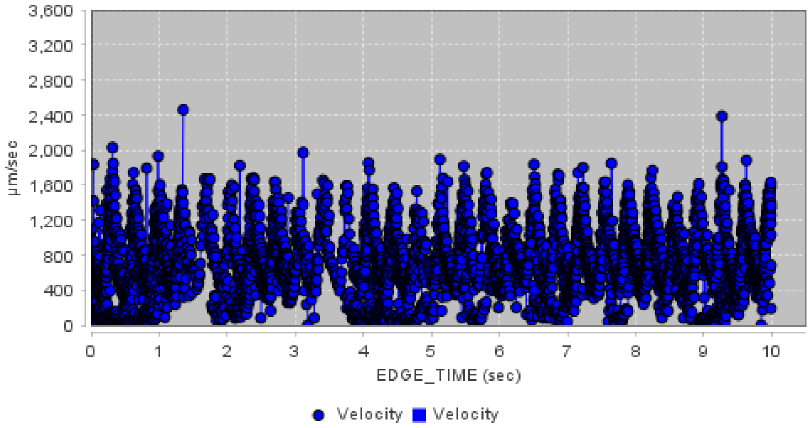


ISVs

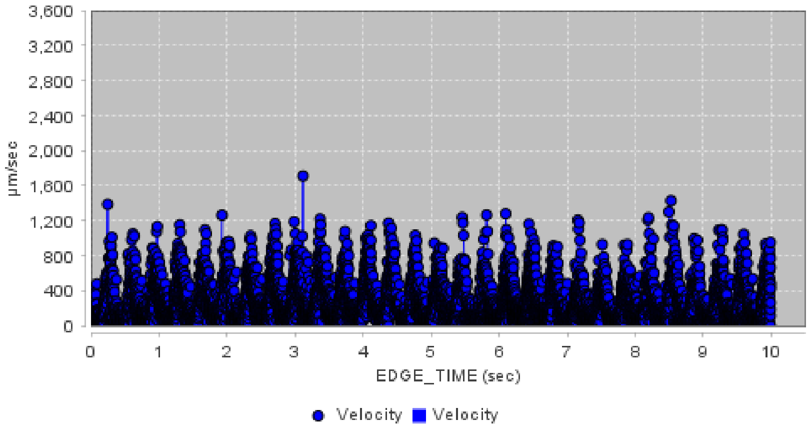


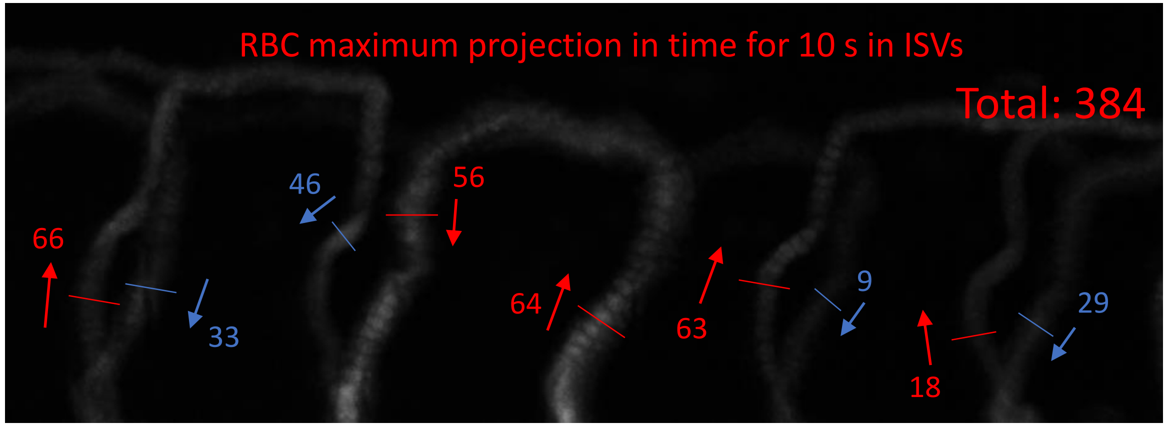


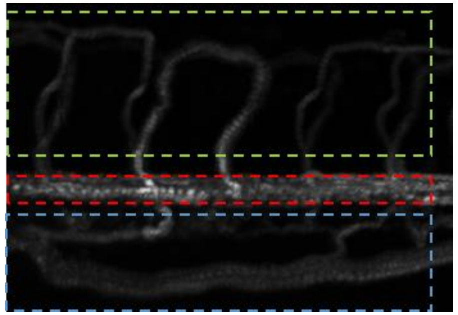

ISVs

CA

CV

### MO Gata1 #2: HR = 174 bpm; Low hematocrit

CA 29 beats in 10 seconds

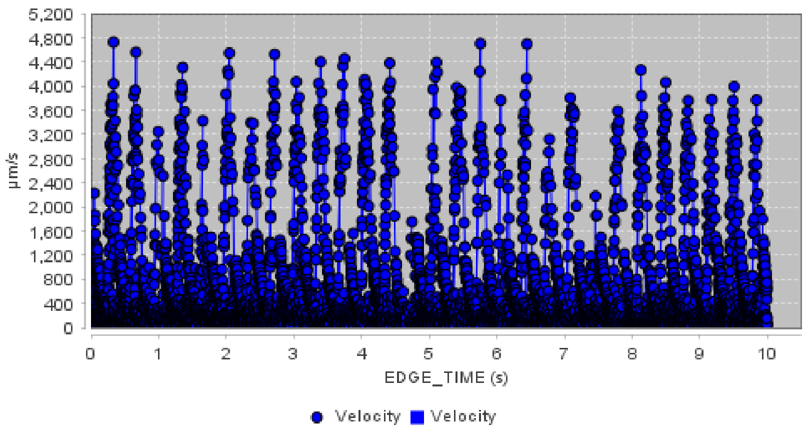


CV

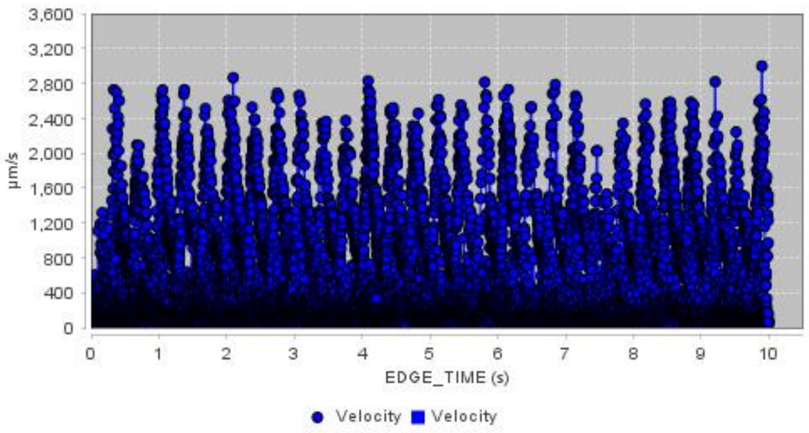


ISVs

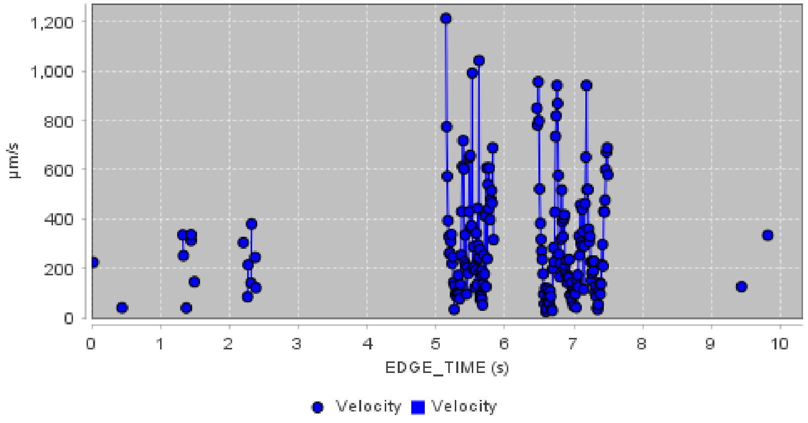


RBC maximum projection in time for 10 s in ISVs

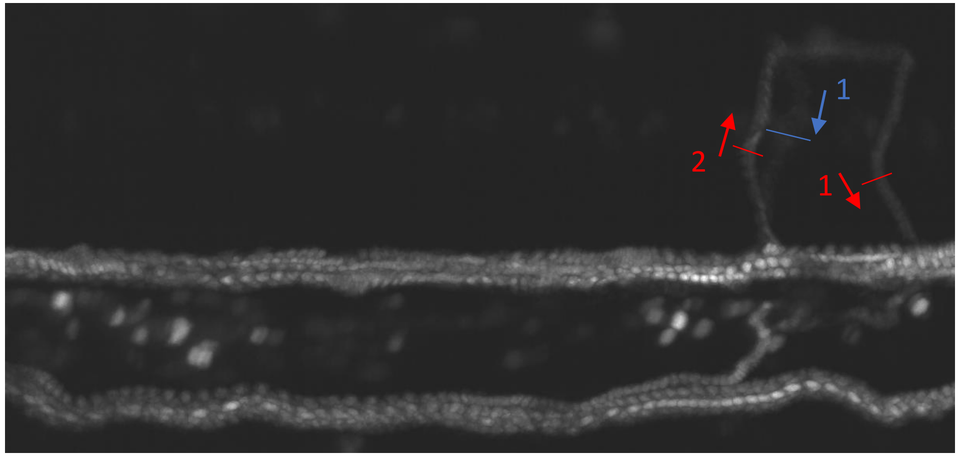


RBC count in ISVs for 10 s

Total: 4

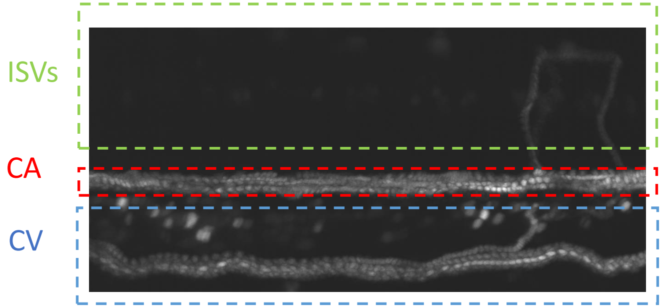

### MO Gata1 #3: HR = 186 bpm; Low hematocrit

CA
31 beats in 10 seconds

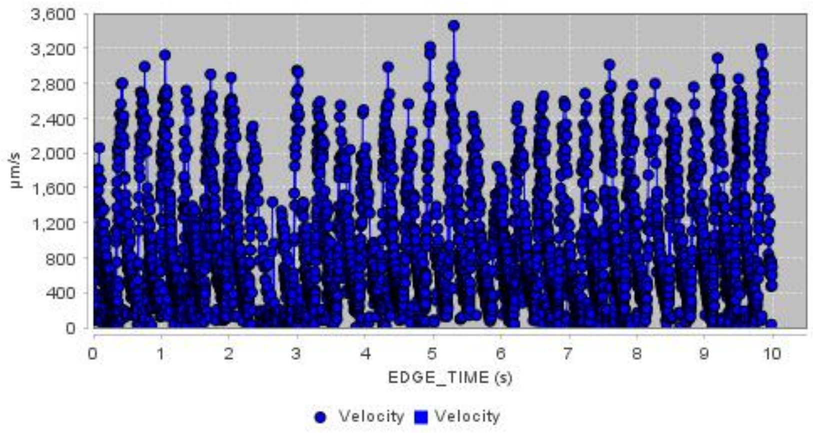


CV

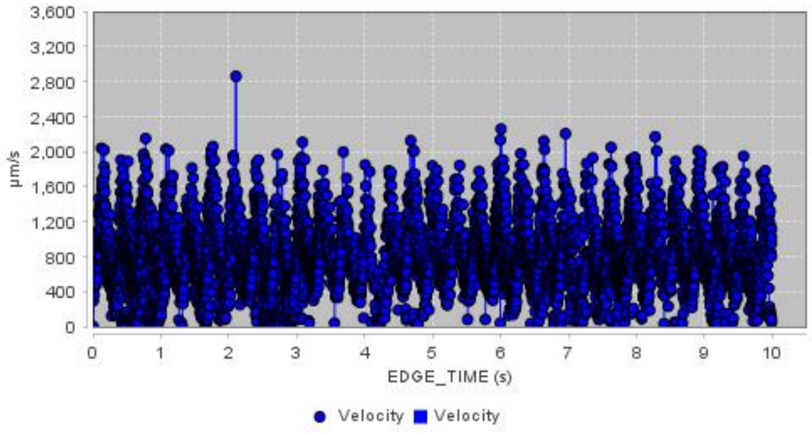


ISVs

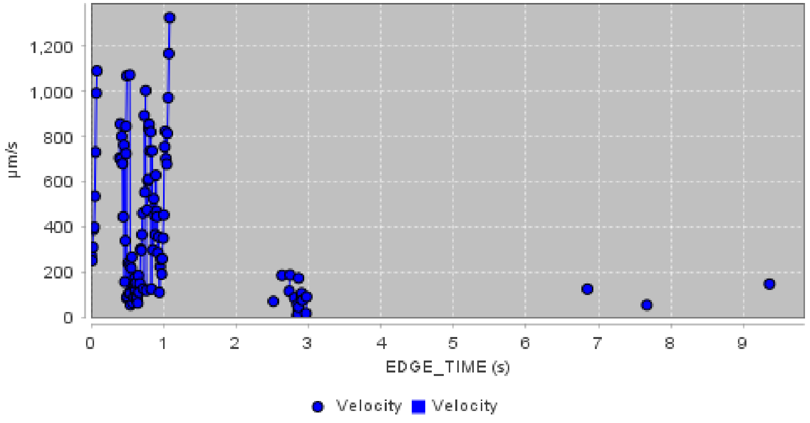


RBC maximum projection in time for 10 s in ISVs

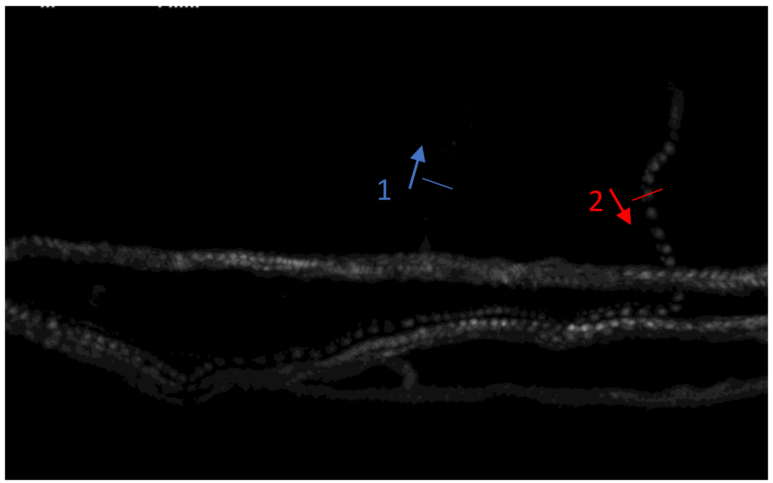


RBC count in ISVs for 10 s

Total: 3

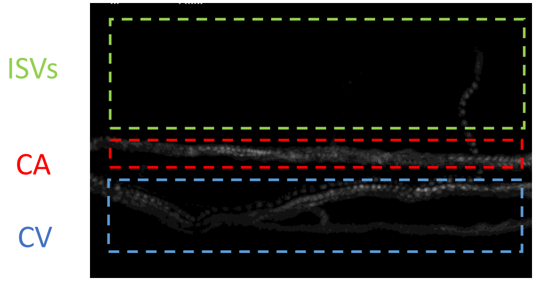

### MO Gata1 #4: HR = 156 bpm; Moderate hematocrit

CA

26 beats in 10 seconds

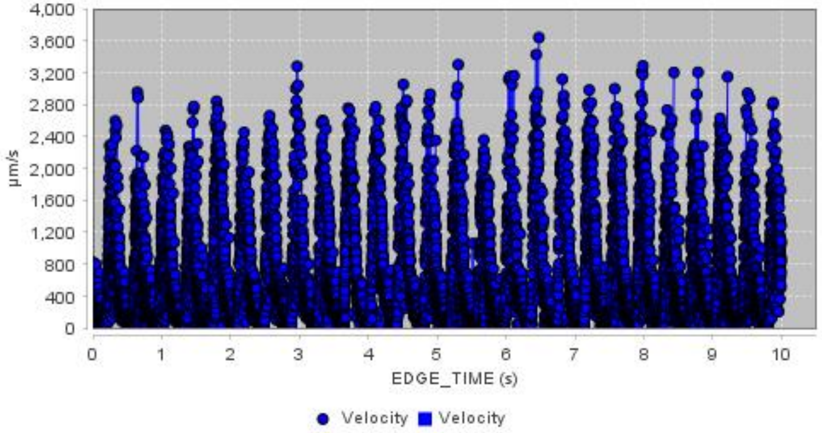


CV

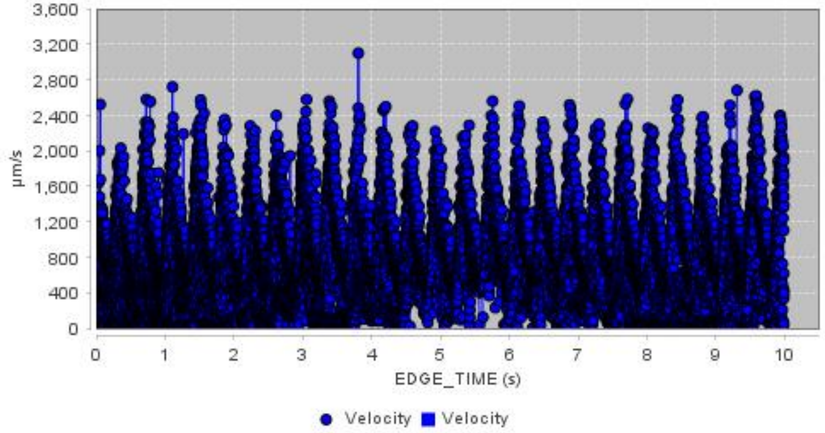


ISVs

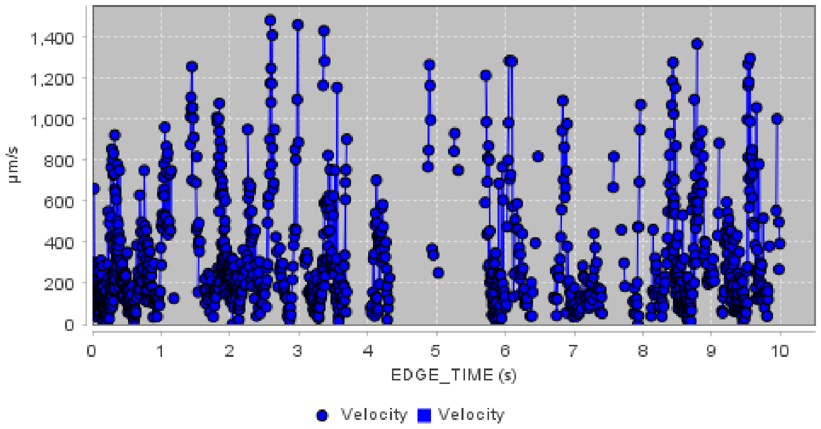


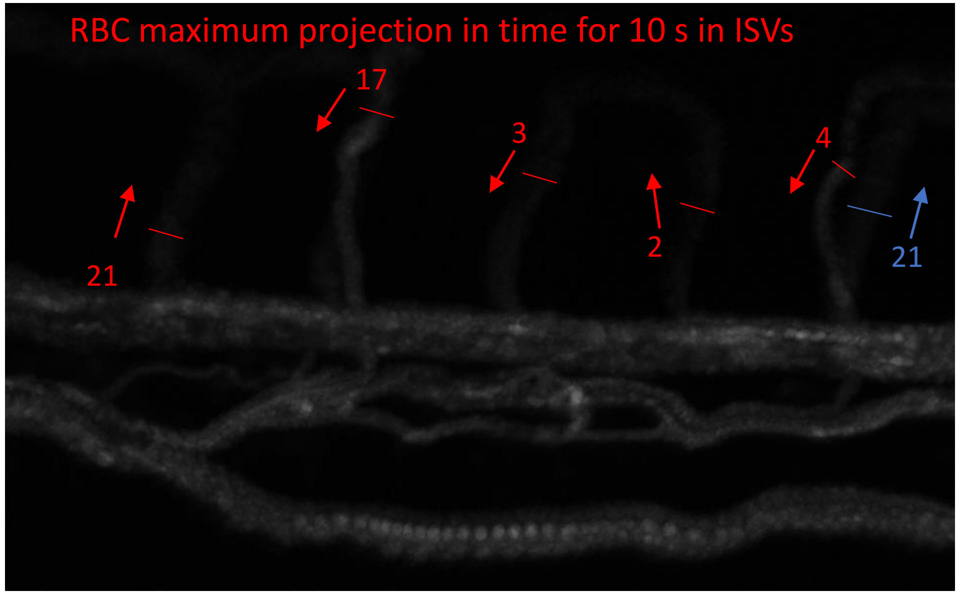


RBC count in ISVs for 10 s

Total: 68

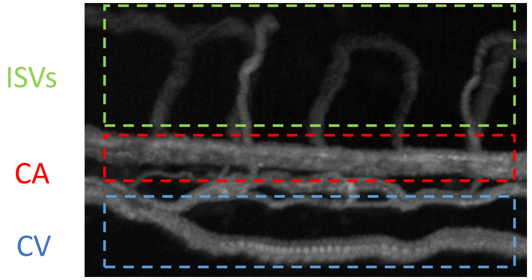

### MO Gata1 #5: HR = 180 bpm; Moderate hematocrit

CA

30 beats in 10 seconds

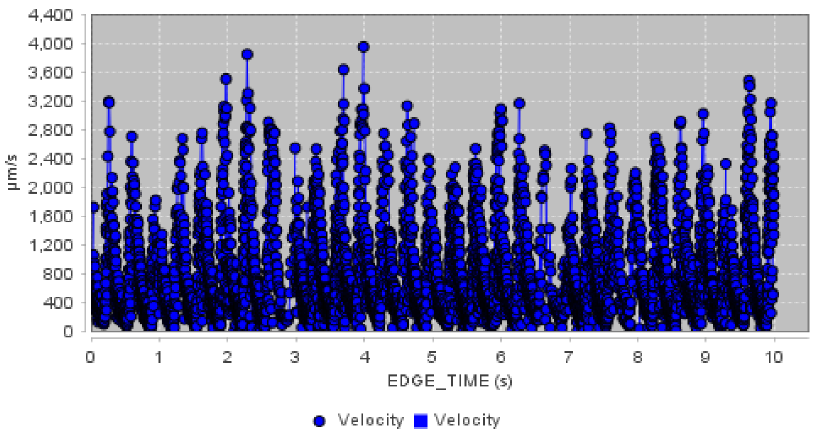


CV

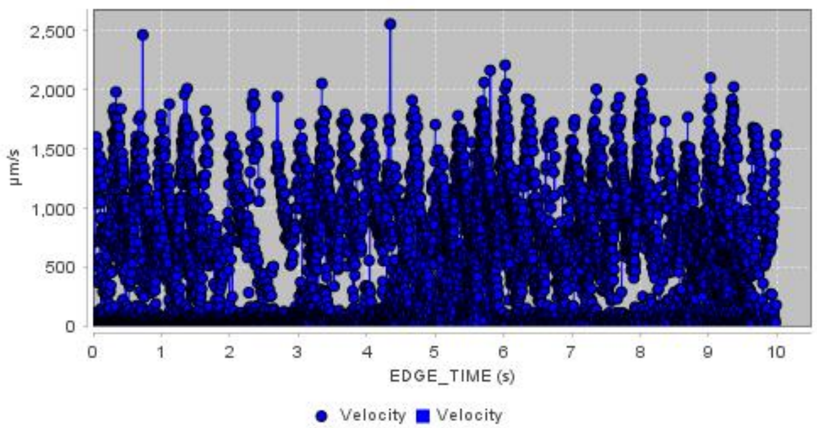


ISVs

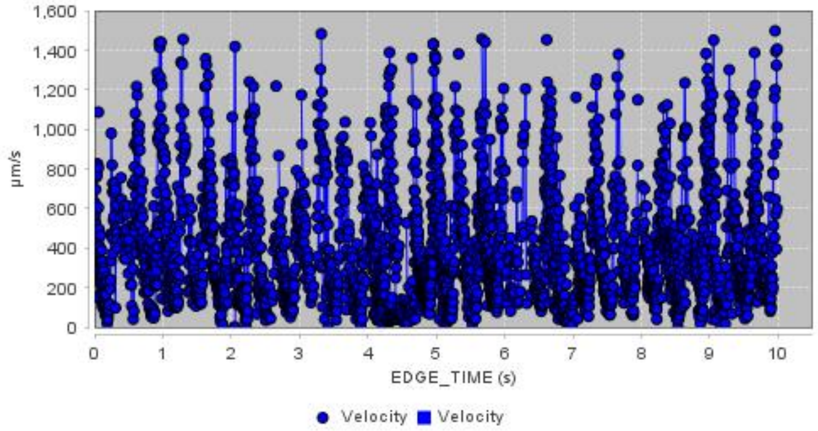


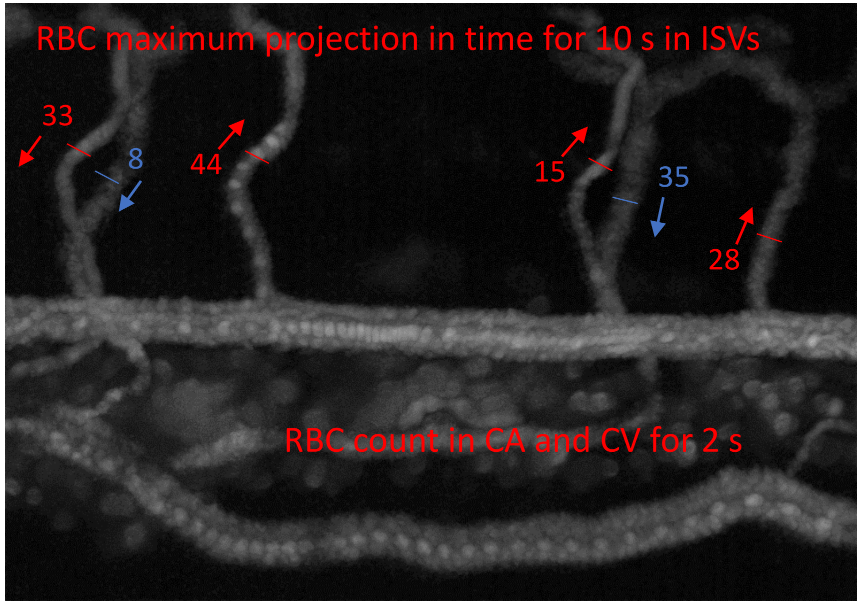


RBC count in ISVs for 10 s

Total in ISVs: 163

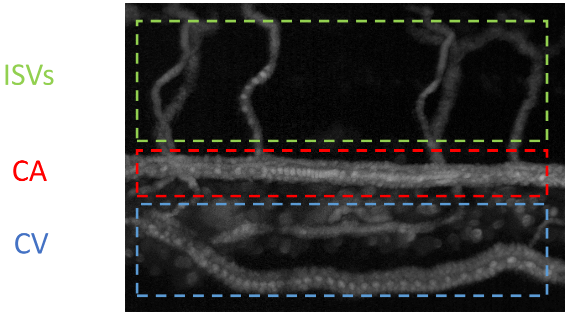

### MO Gata1 #6: HR = 147 bpm; Low hematocrit

CA 24.5 beats in 10 seconds

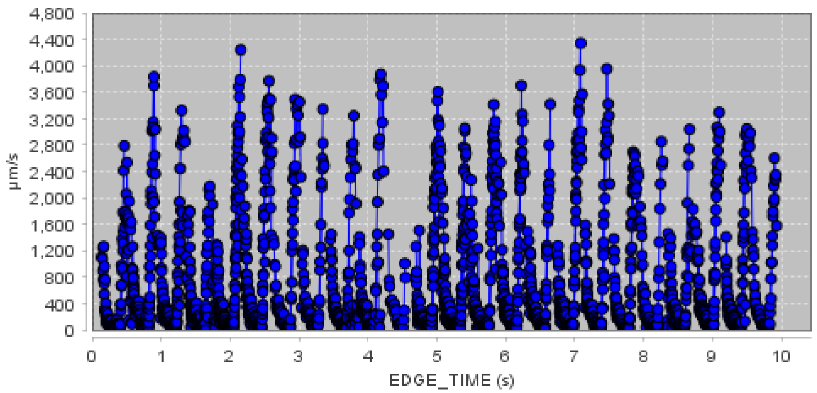


CV

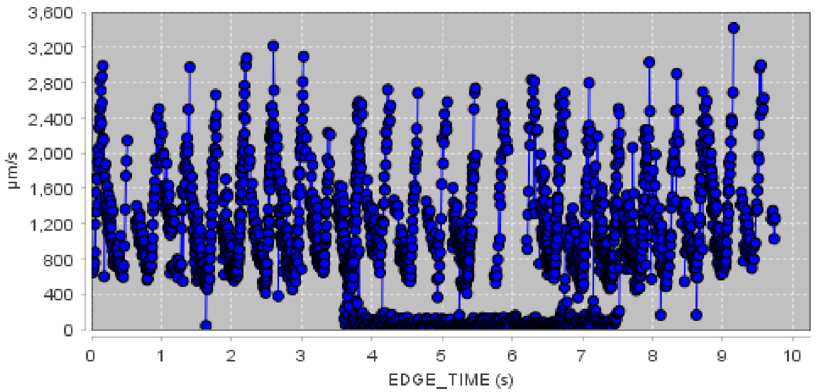


ISVs

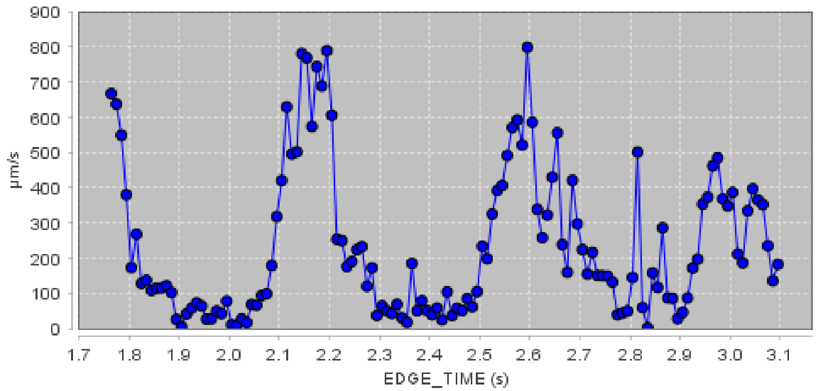


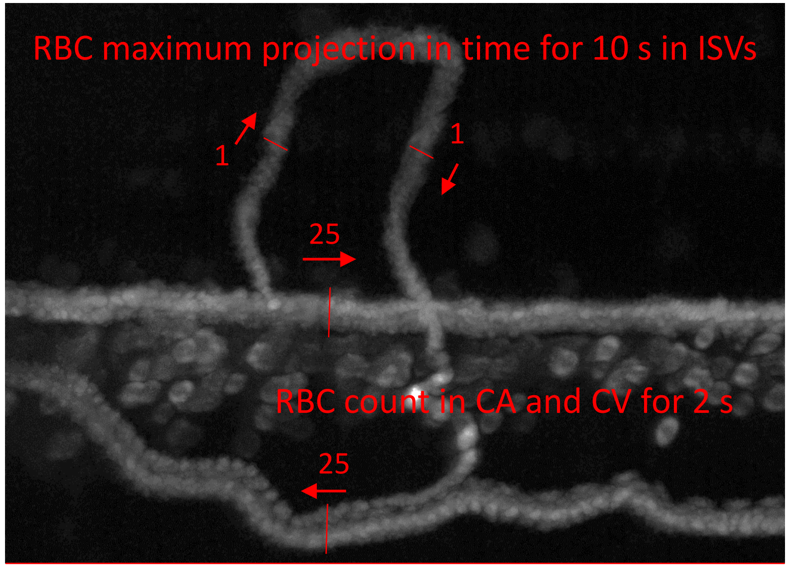


RBC count in ISVs for 10 s

Total in ISVs: 2

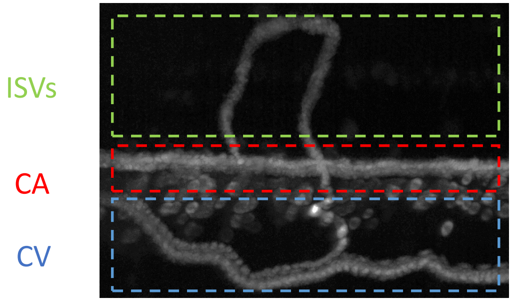

### MO Gata1 #7: HR = 150 bpm; Moderate hematocrit

CA

25 beats in 10 seconds

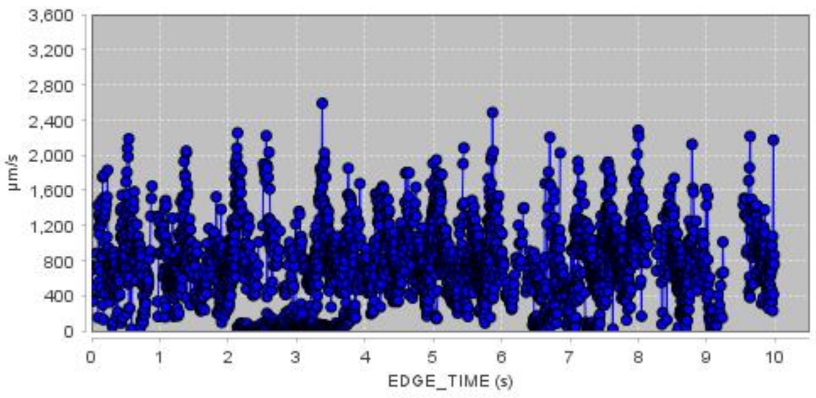


CV

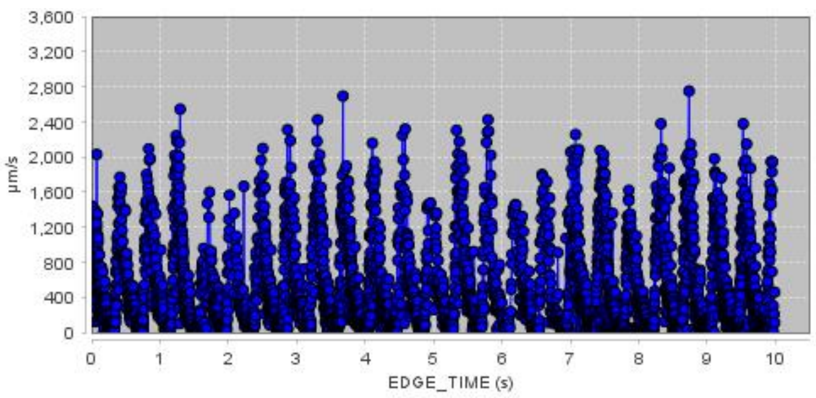


ISVs

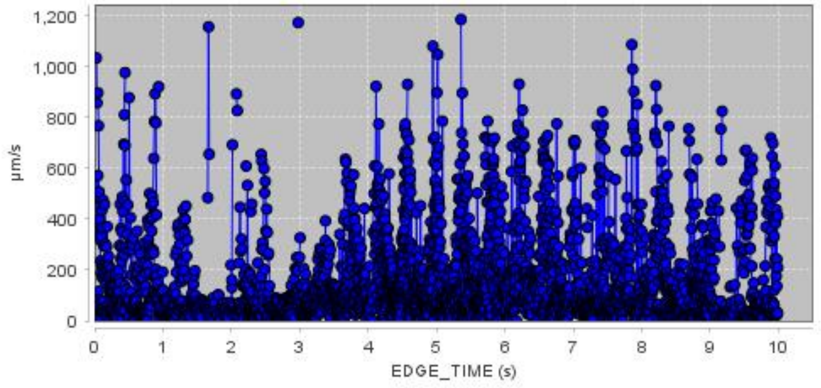


RBC maximum projection in time for 10 s in ISVs

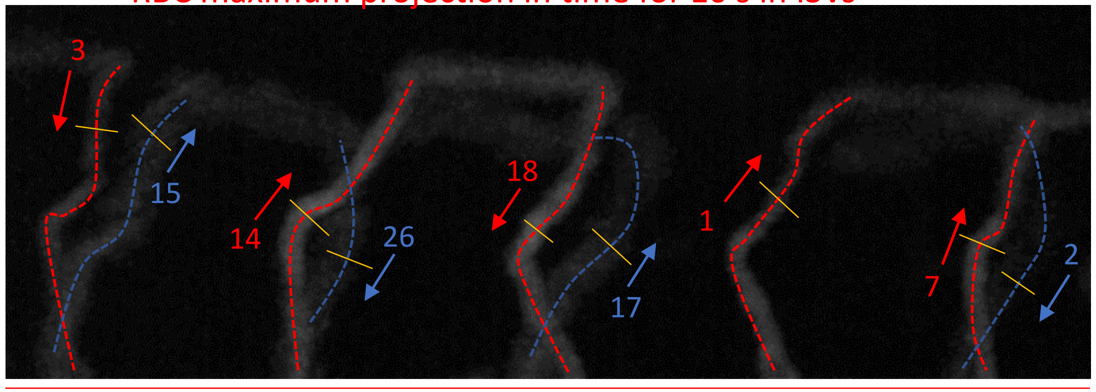


RBC count in ISVs for 10 s

Total in ISVs: 103

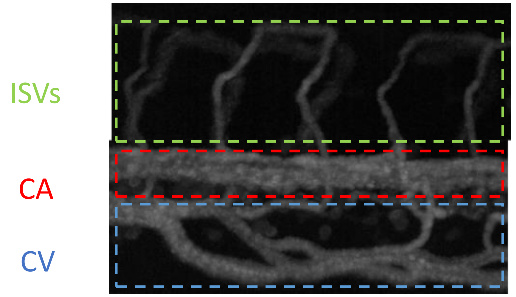

### Section B:

#### Wildtype Zebrafish, a perfusion quantification experiment

### WT #18: HR = 168 bpm; Good perfusion

CA

28 beats in 10 seconds

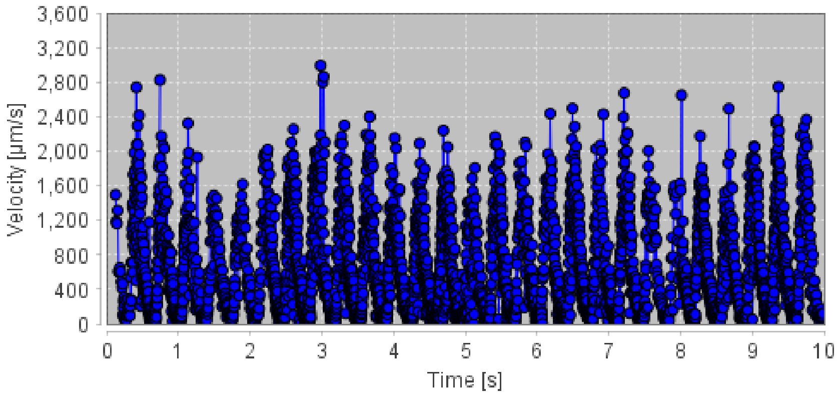


CV

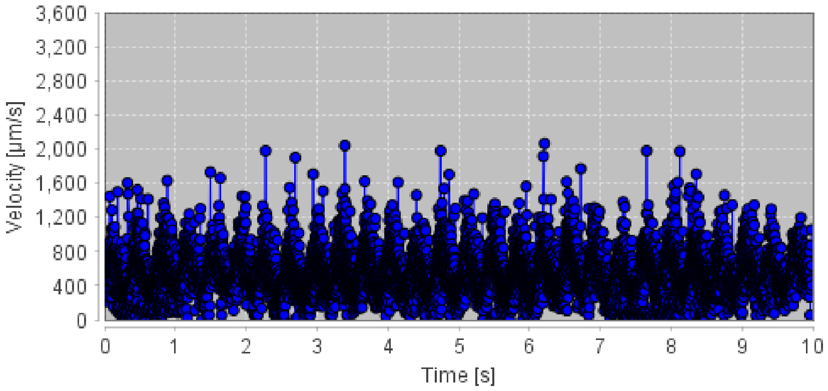


ISVs

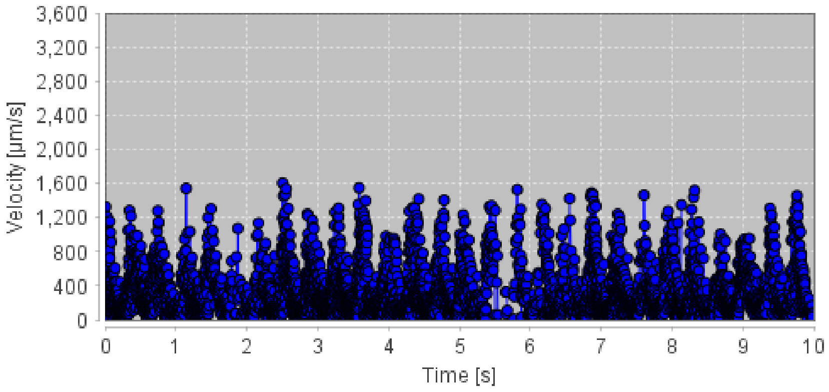


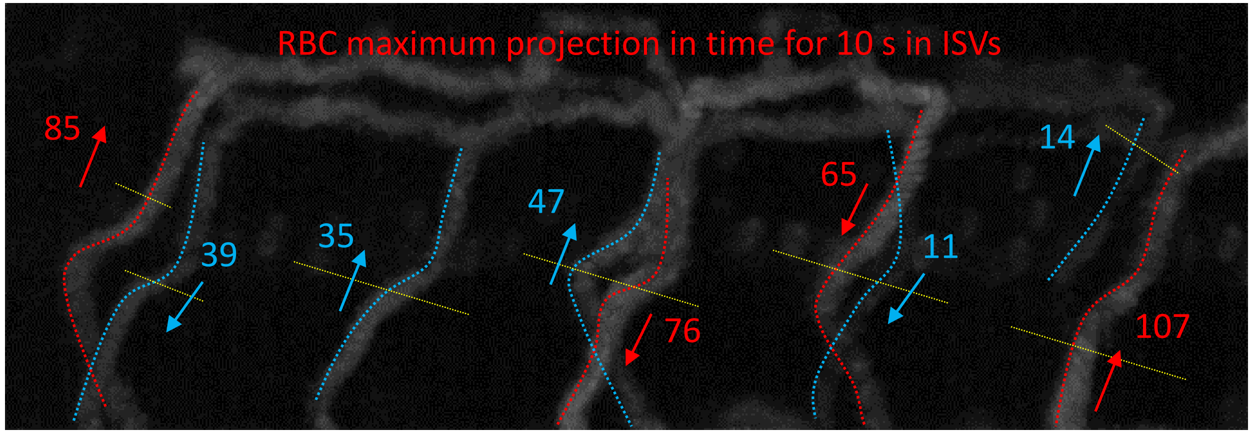


RBC count in ISVs for 10 s

Total: 479

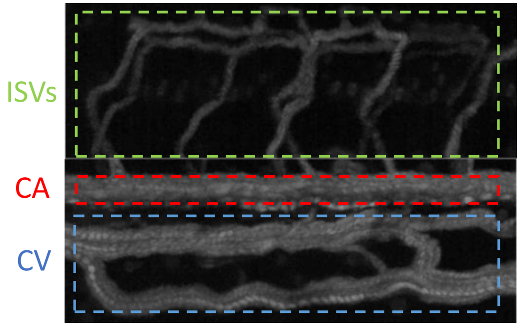

### WT #8: HR = 168 bpm; Good perfusion

CA

28 beats in 10 seconds

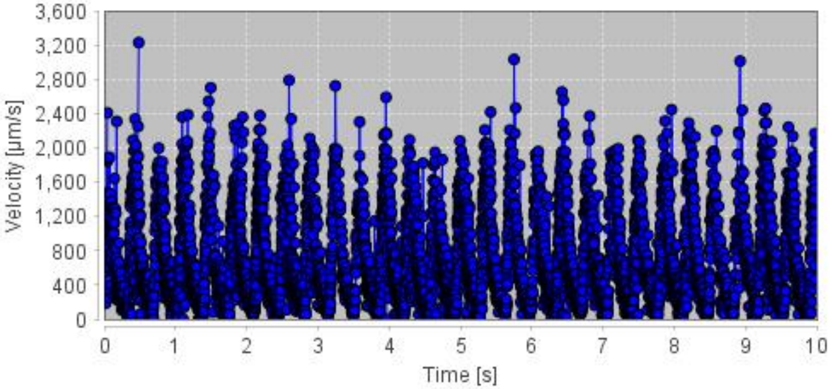


CV

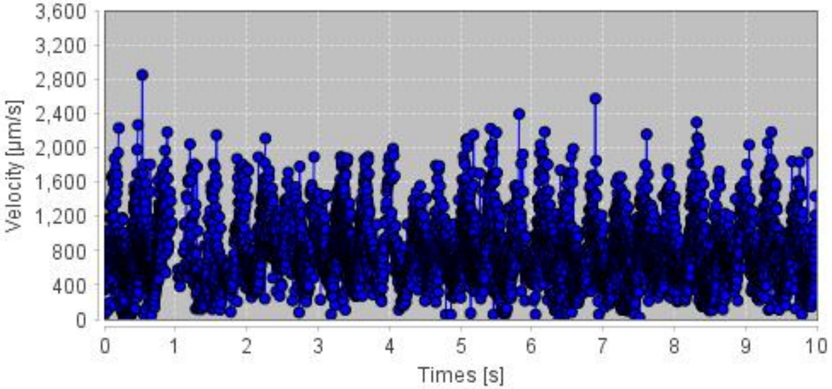


ISVs

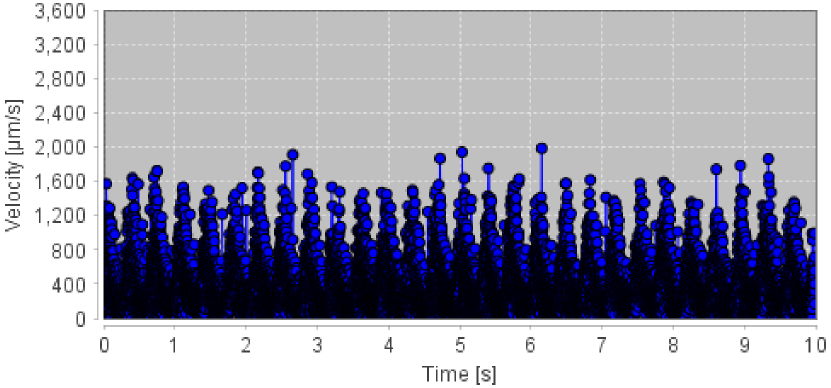


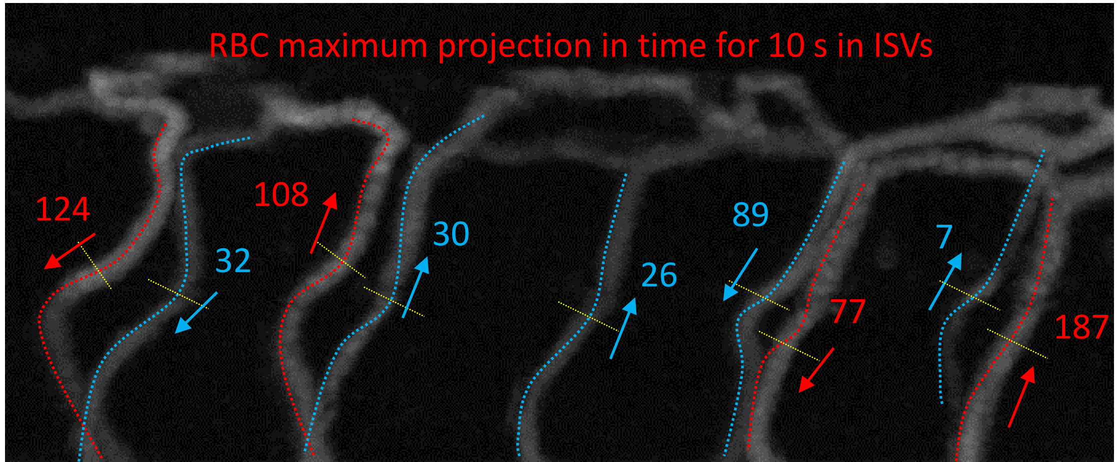


RBC count in ISVs for 10 s

Total: 680

ISVs

CA

CV

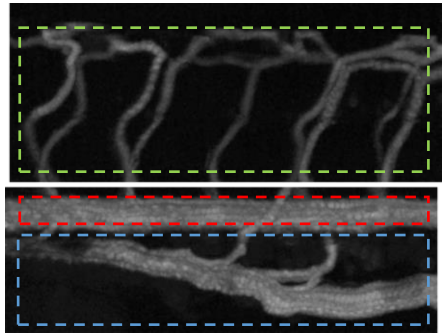

### WT #10: HR = 156 bpm; Good perfusion

CA 26 beats in 10 seconds

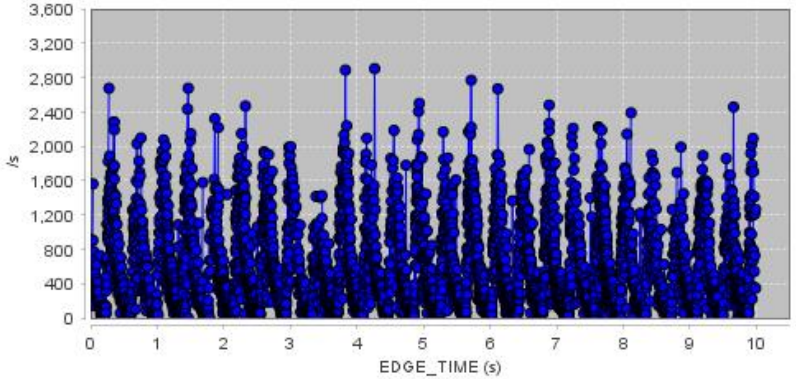


CV

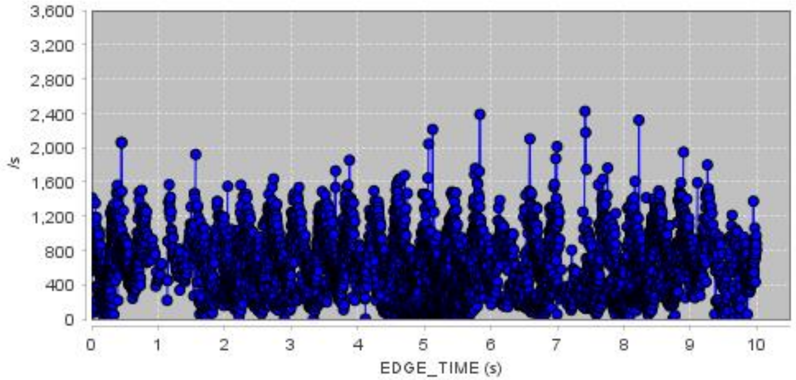


ISVs

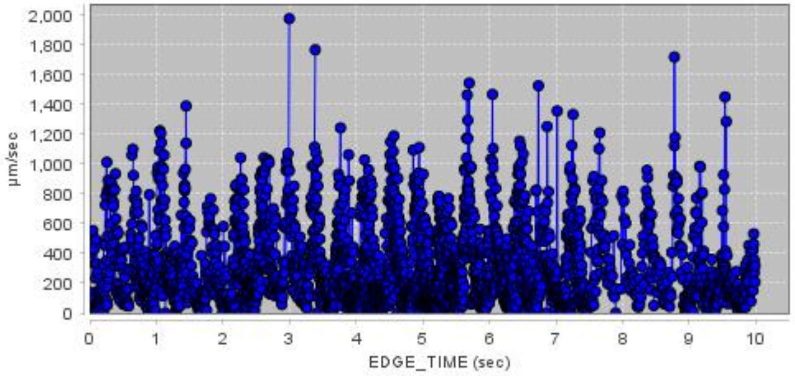


RBC maximum projection in time for 10 s in ISVs

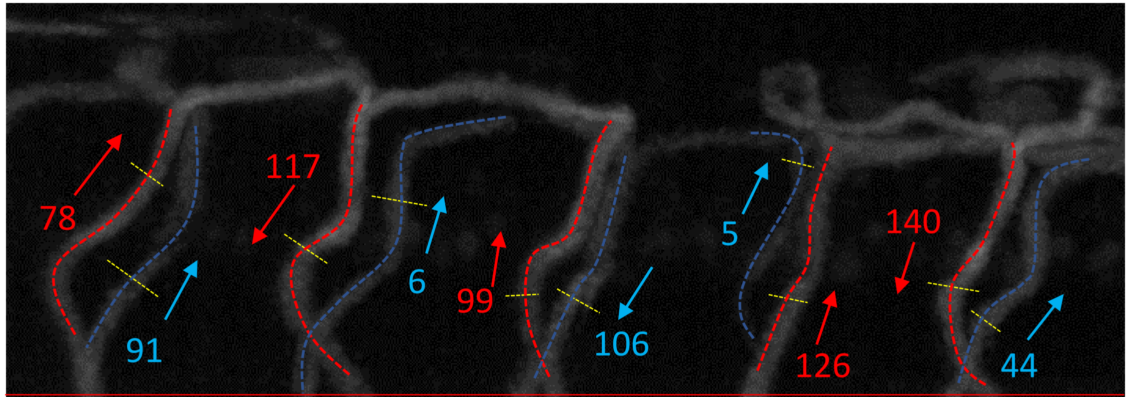


RBC count in ISVs for 10 s

Total: 812

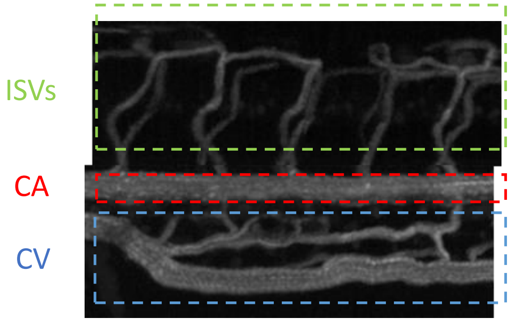

### WT #3: HR = 156 bpm; Good perfusion

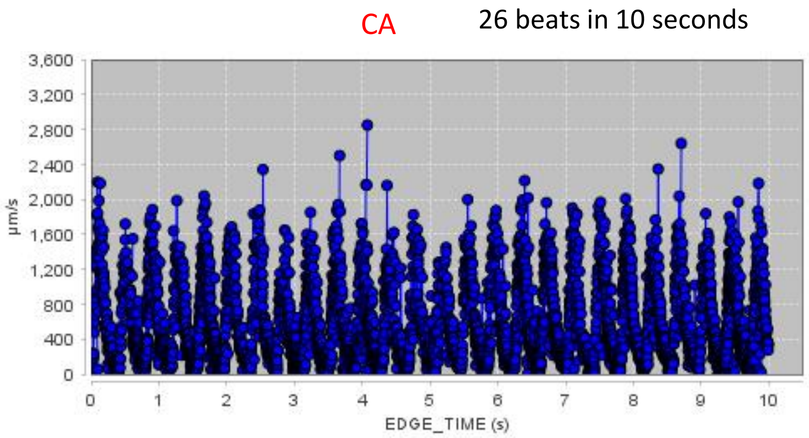


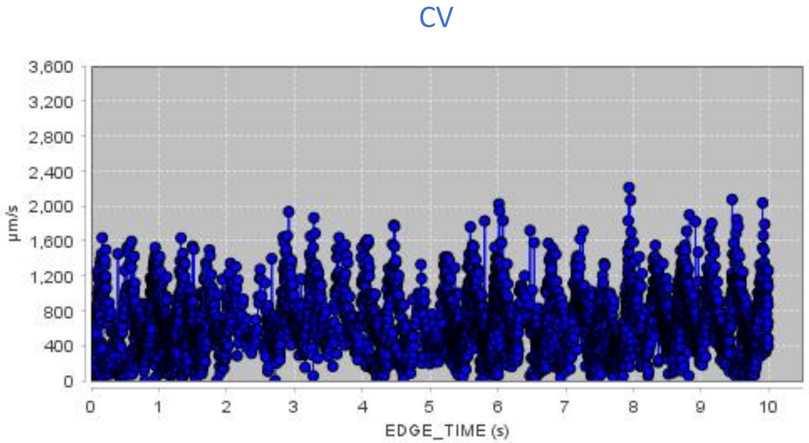


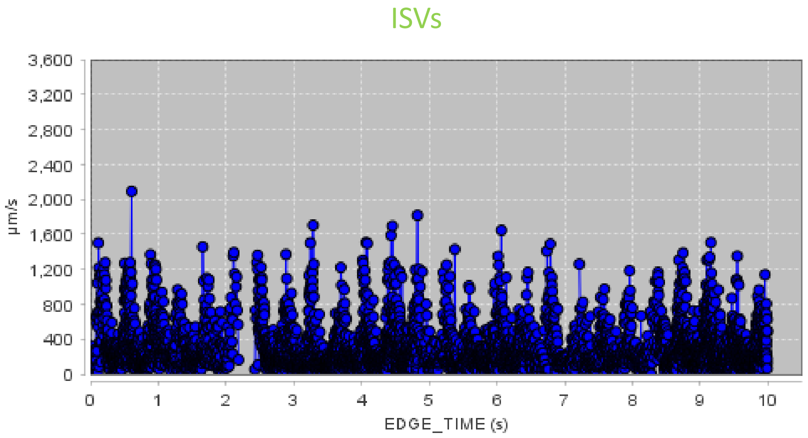


RBC maximum projection in time for 10 s in ISVs

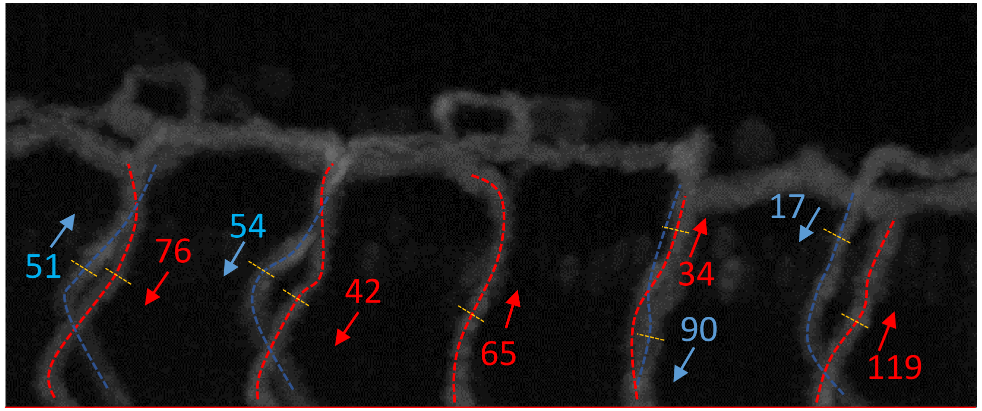


RBC count in ISVs for 10 s

Total: 548

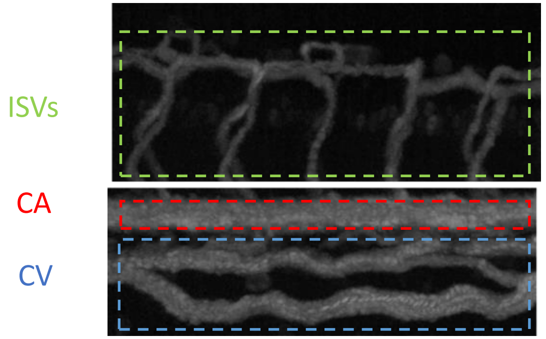

### WT #22: HR = 162 bpm; Low ISV perfusion

CA 27 beats in 10 seconds

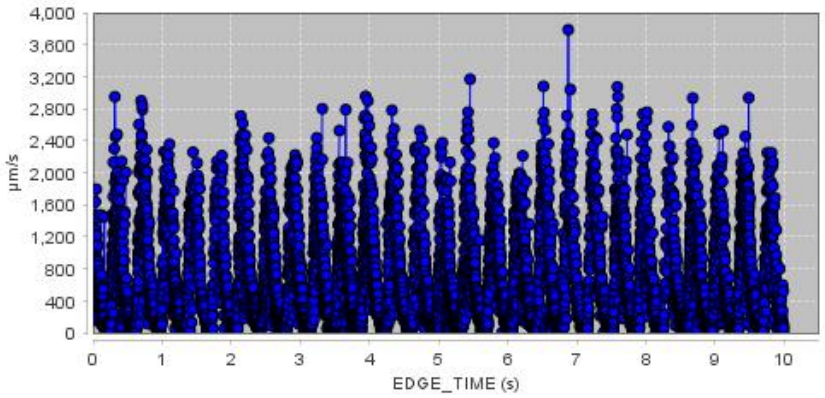


CV

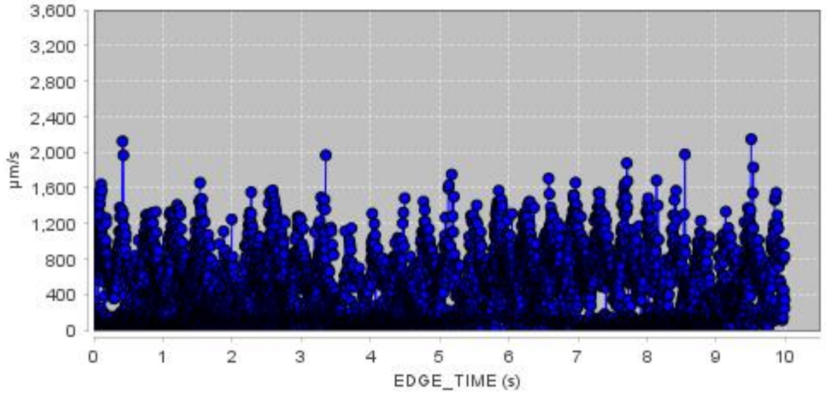


ISVs

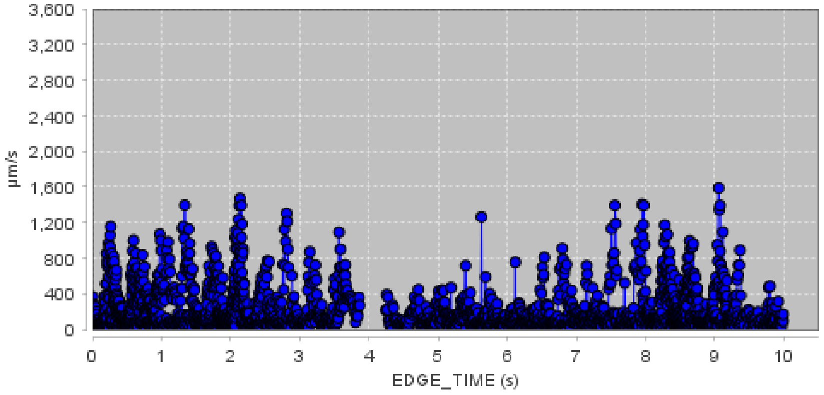


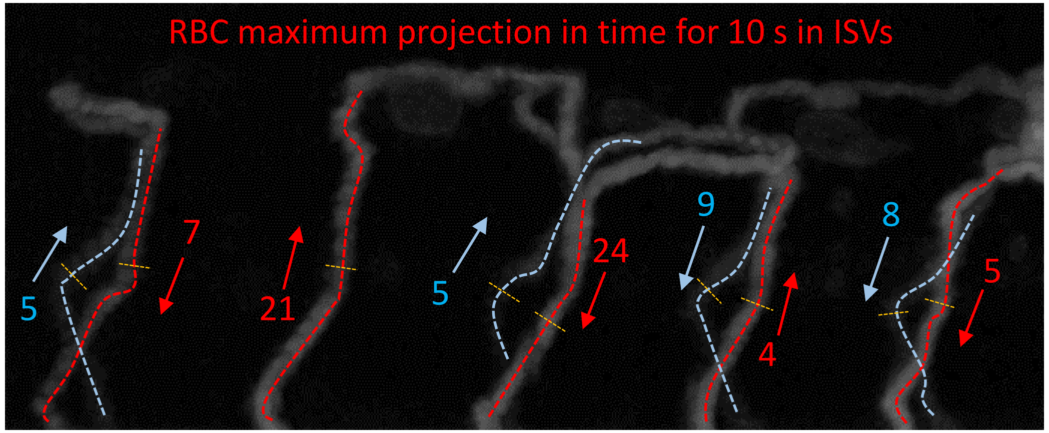


RBC count in ISVs for 10 s

Total: 88

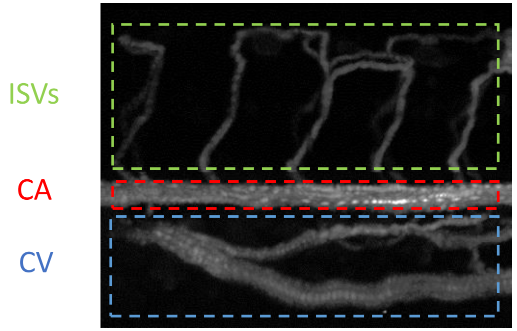

### Section C:

#### Marcksl1 la lb double knockout  Zebrafish, a lumen diameter alteration and perfusion quantification experiment

### Good perfusion levels in ISVs

### ML1dblKO #6: HR = 162 bpm; Good perfusion

CA 27 beats in 10 seconds

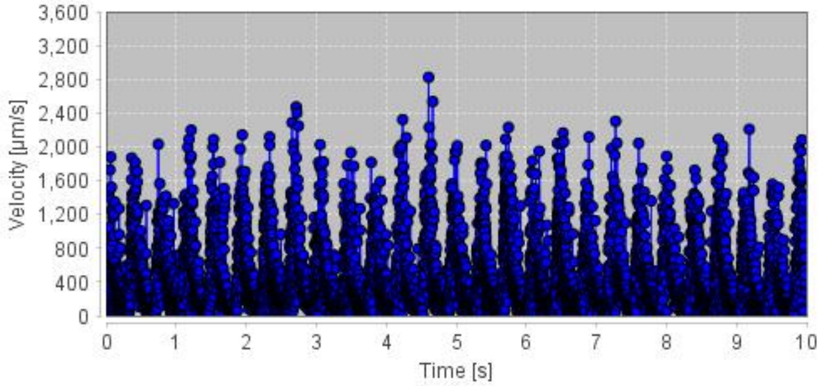


CV

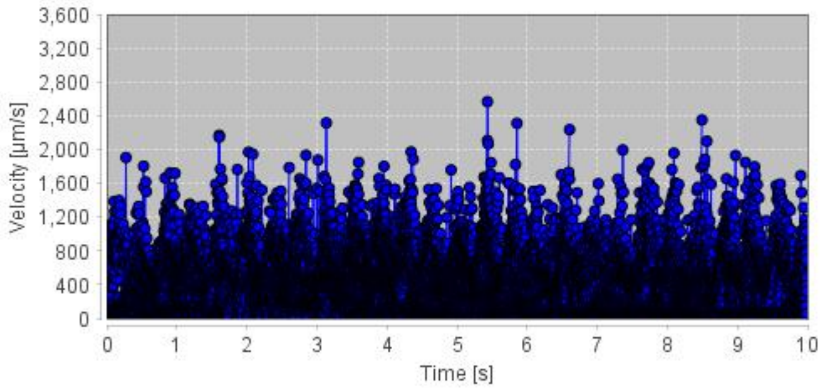


ISVs

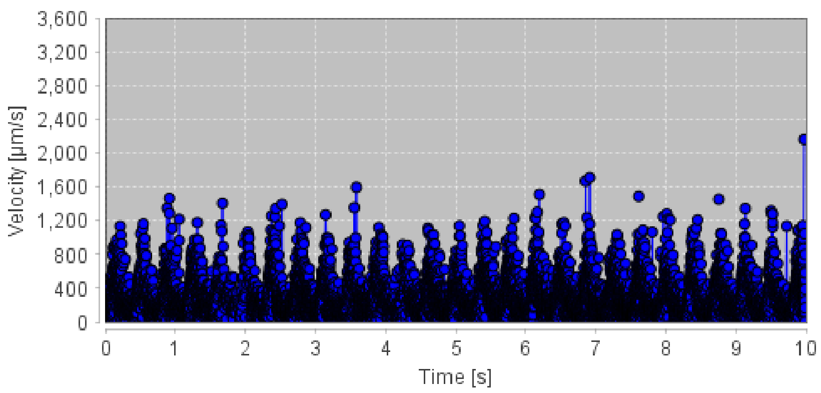


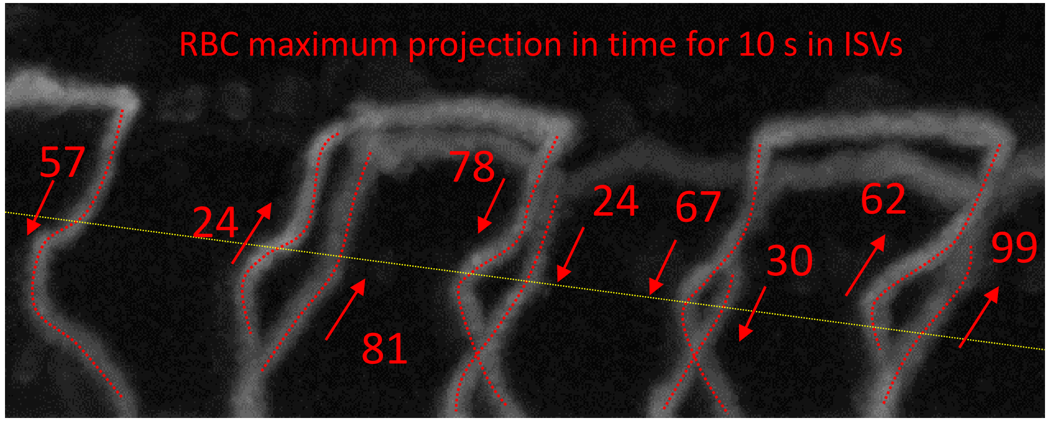


RBC count in ISVs for 10 s

Total: 522

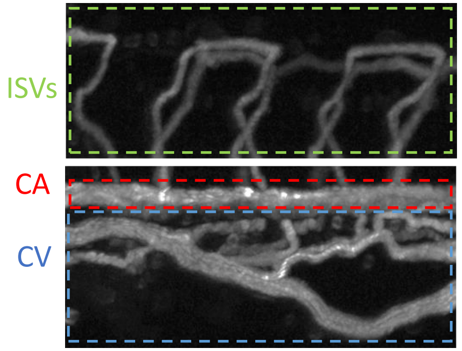

### ML1dblKO #10: HR = 156 bpm; Good perfusion

CA 26 beats in 10 seconds

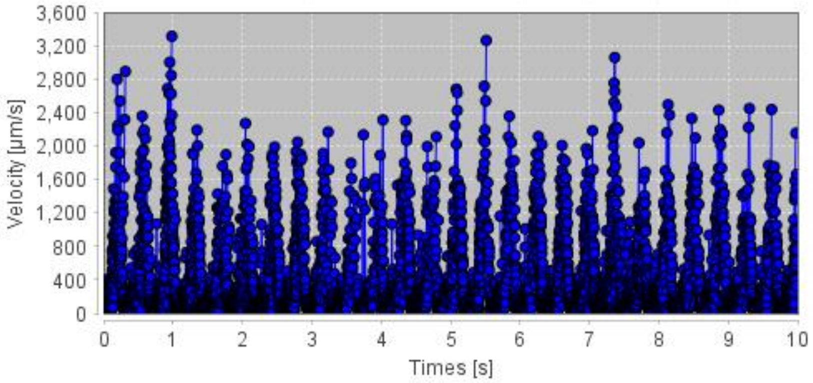


CV

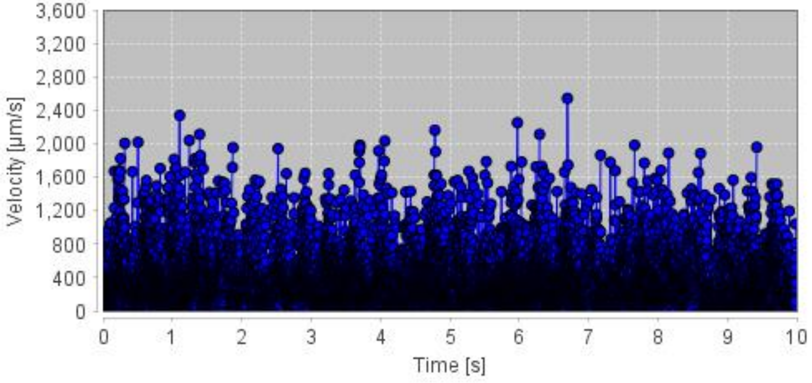


ISVs

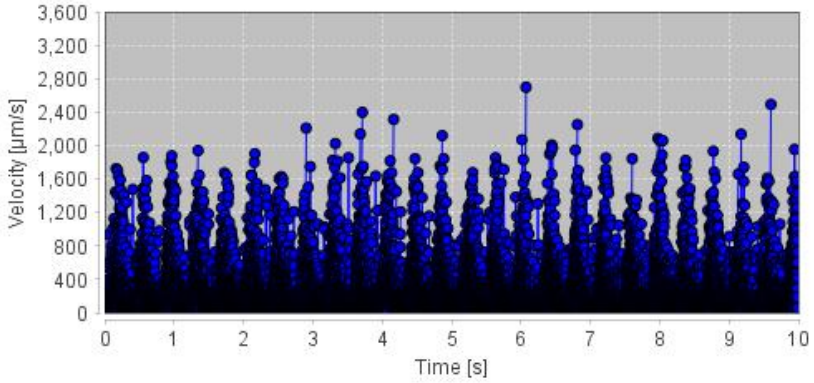


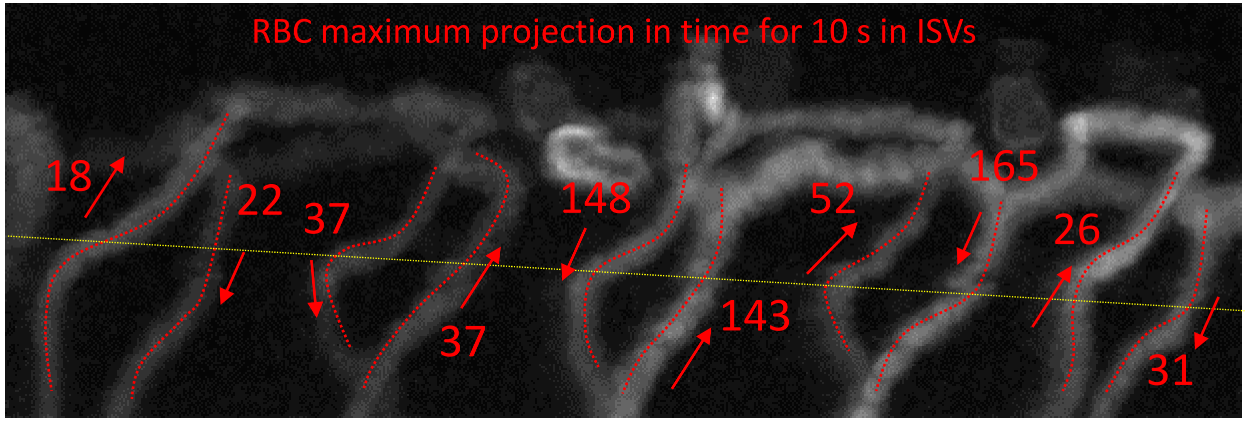


RBC count in ISVs for 10 s

Total: 642

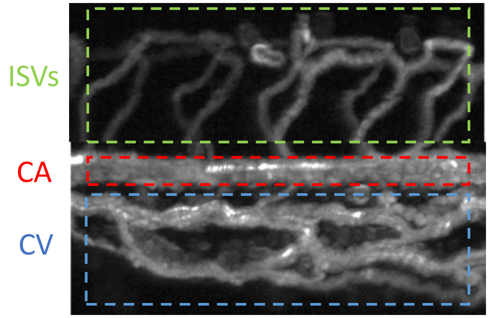

### ML1dblKO #13: HR = 138 bpm; Good perfusion

CA

23 beats in 10 seconds

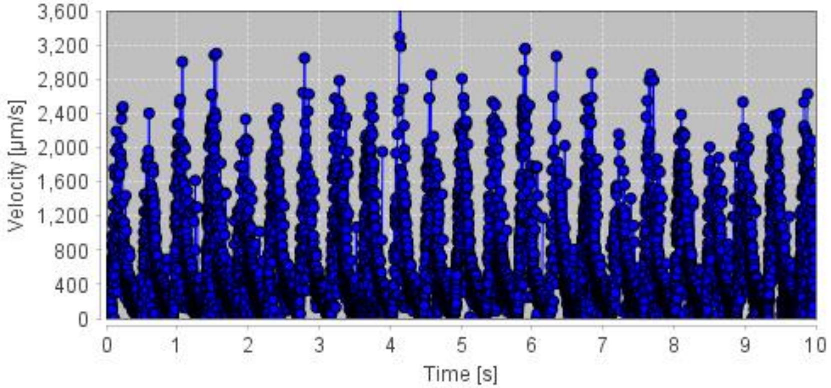


CV

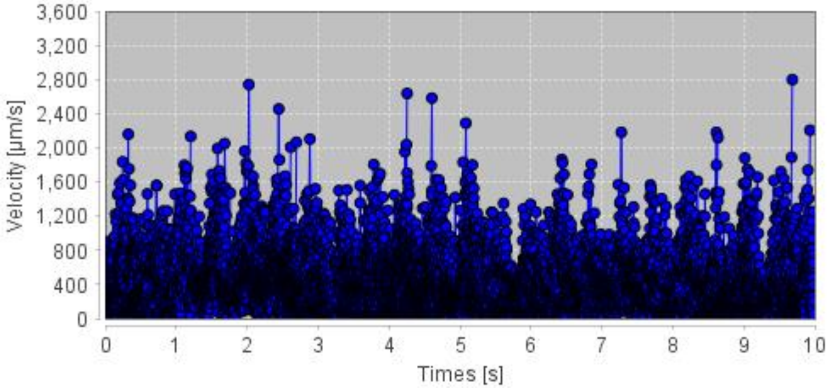


ISVs

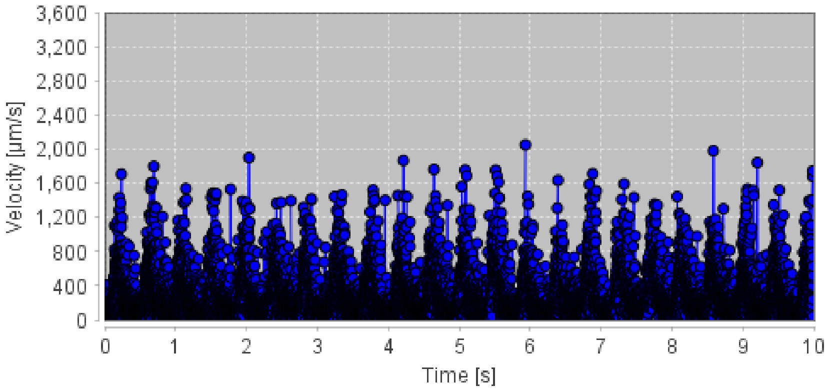


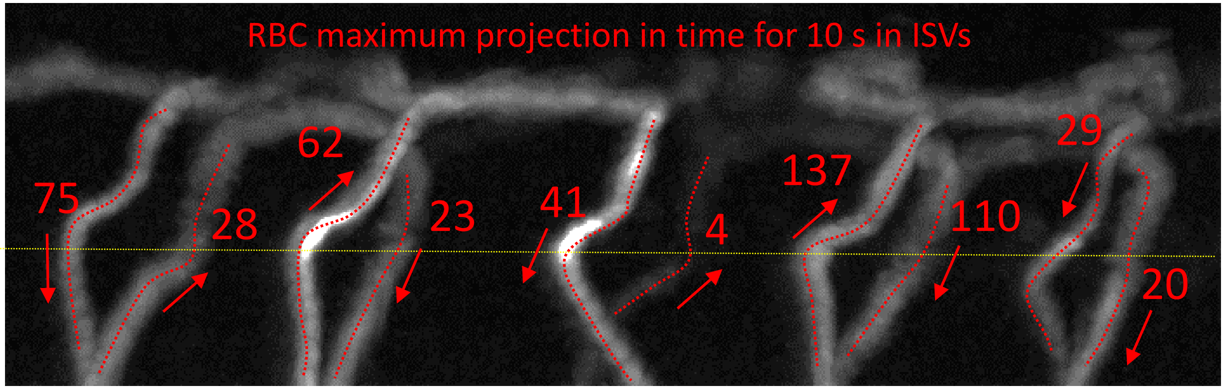


RBC count in ISVs for 10 s

Total: 529

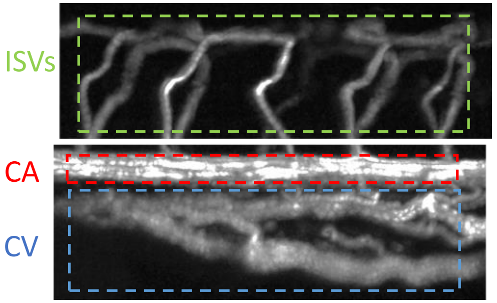

### ML1dblKO #15: HR = 138 bpm; Good perfusion

CA

23 beats in 10 seconds

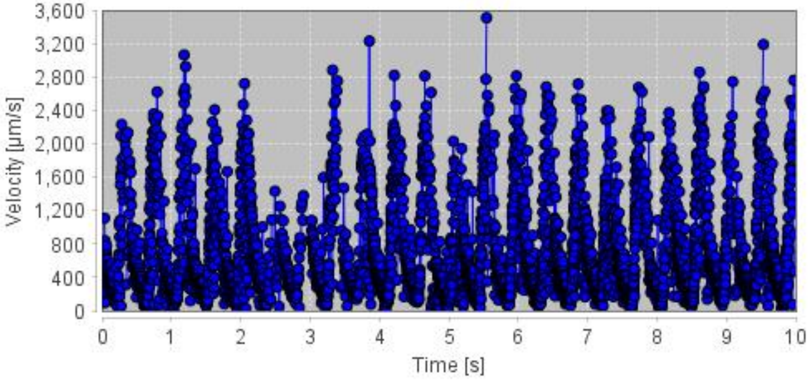


CV

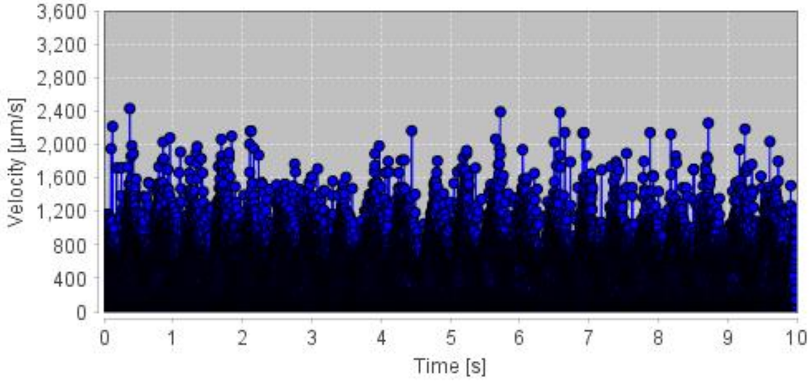


ISVs

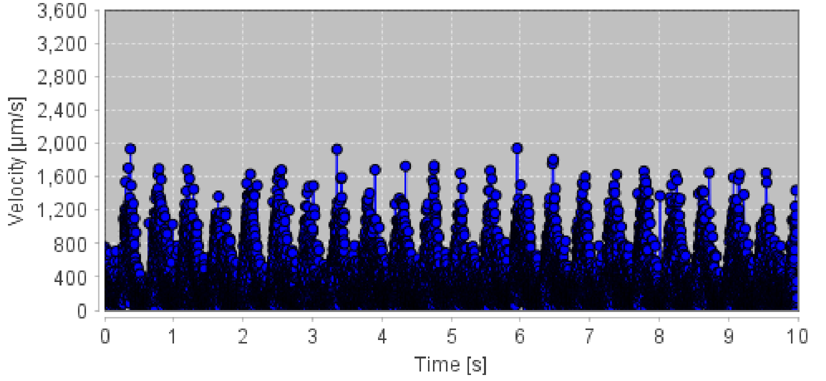


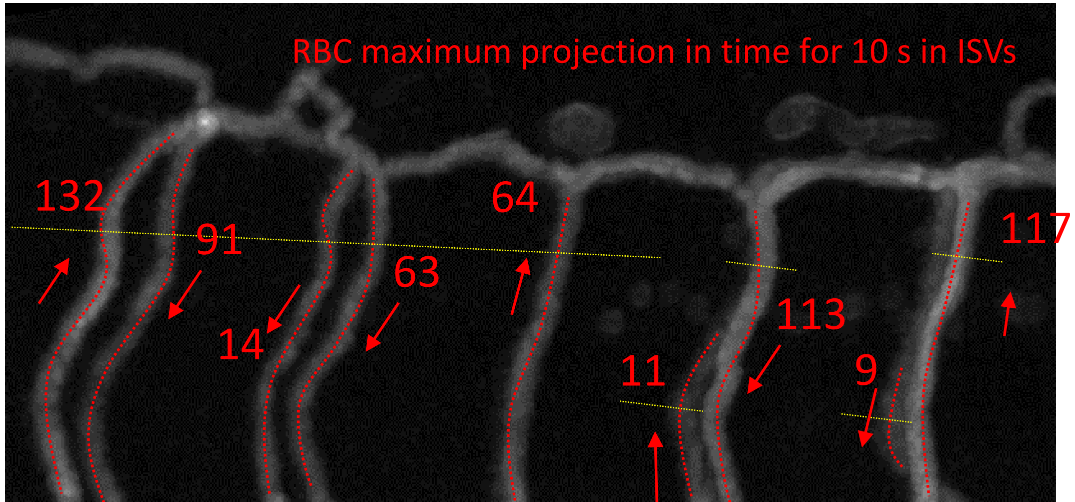


RBC count in ISVs for 10 s

Total: 614

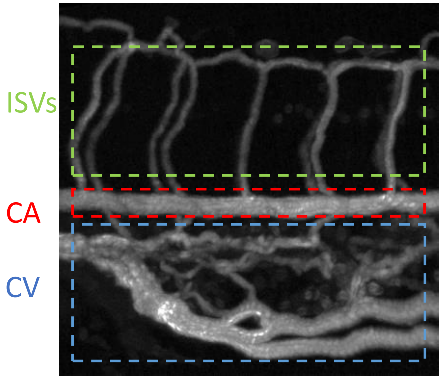

### Moderate perfusion levels in ISVs

### ML1dblKO #1: HR = 138 bpm; Moderate perfusion

CA

23 beats in 10 seconds

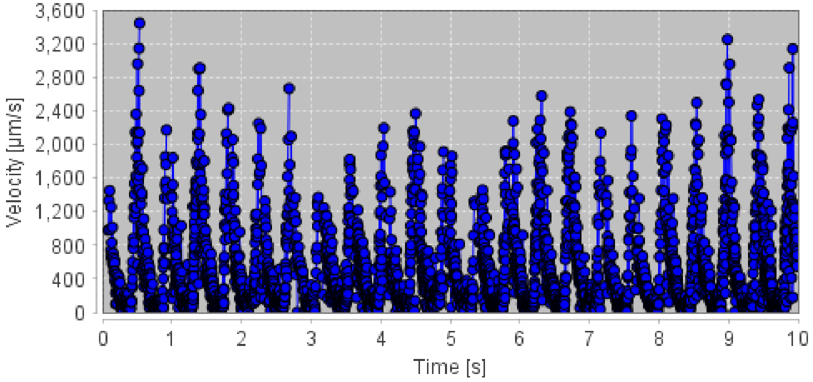


CV

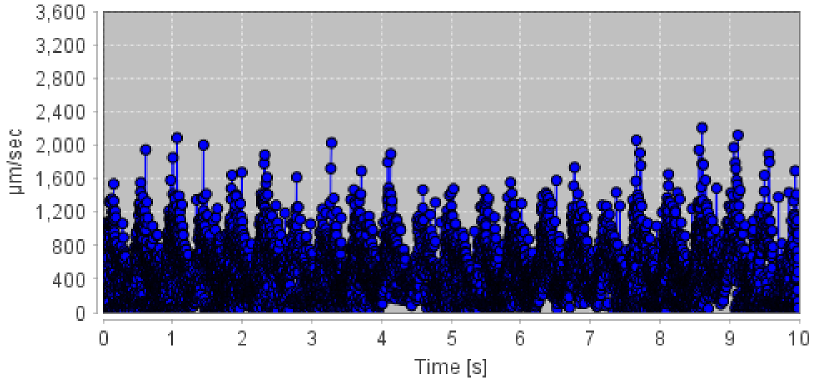


ISVs

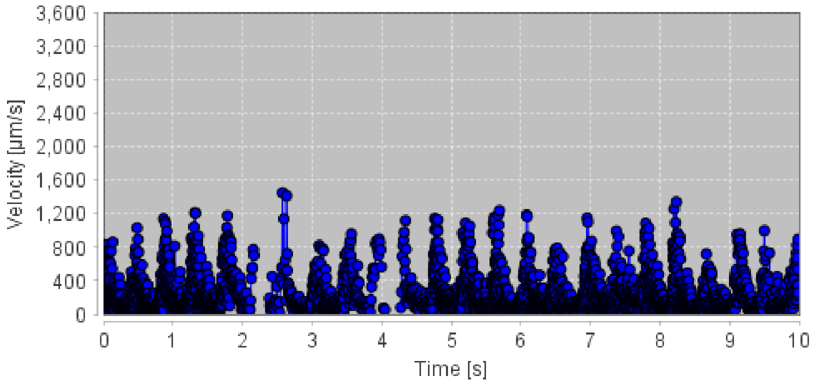


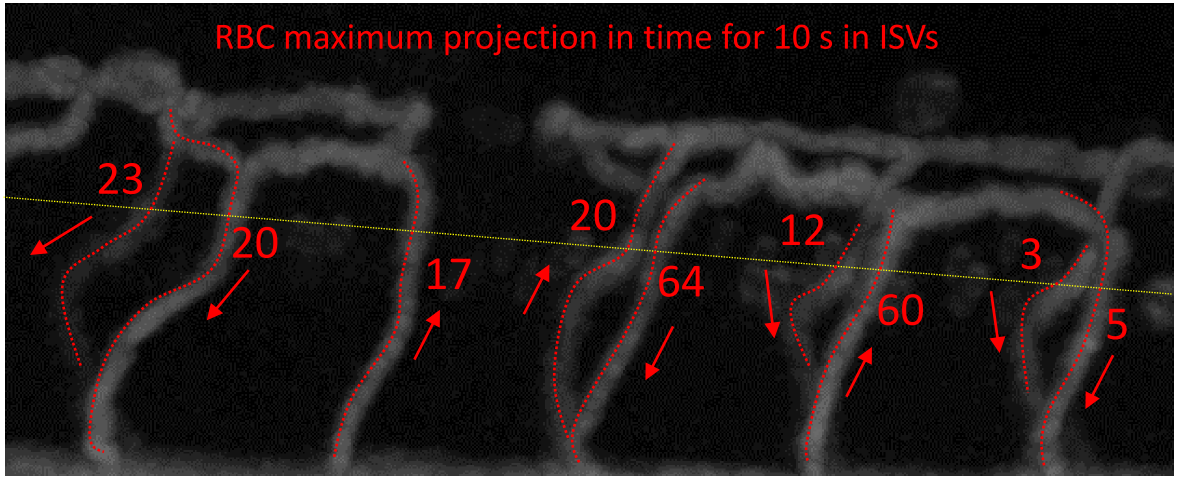


RBC count in ISVs for 10 s

Total: 224

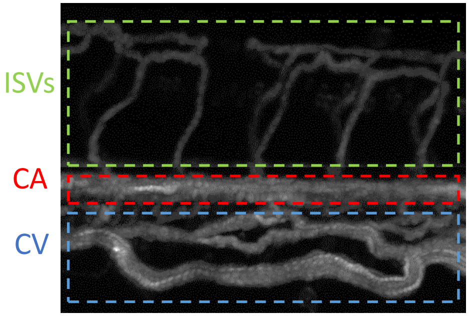

### ML1dblKO #2: HR = 144 bpm; Moderate perfusion

CA

24 beats in 10 seconds

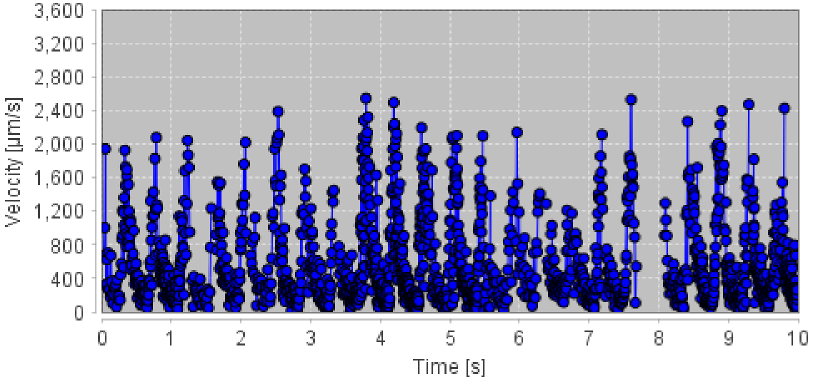


CV

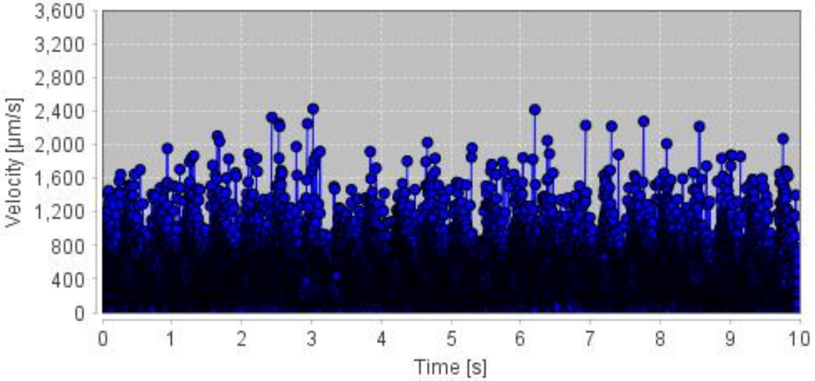


ISVs

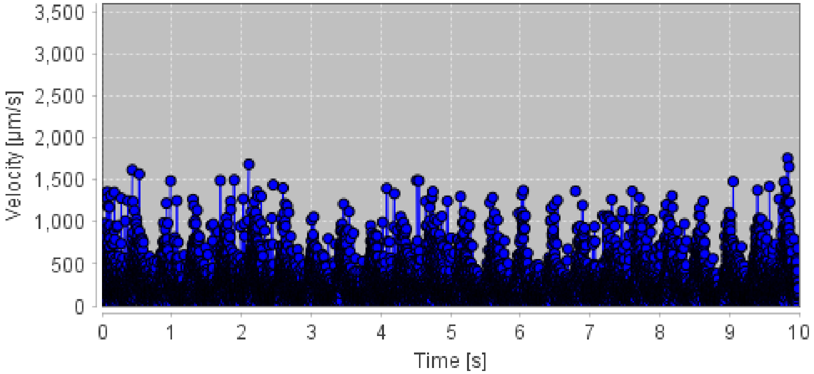


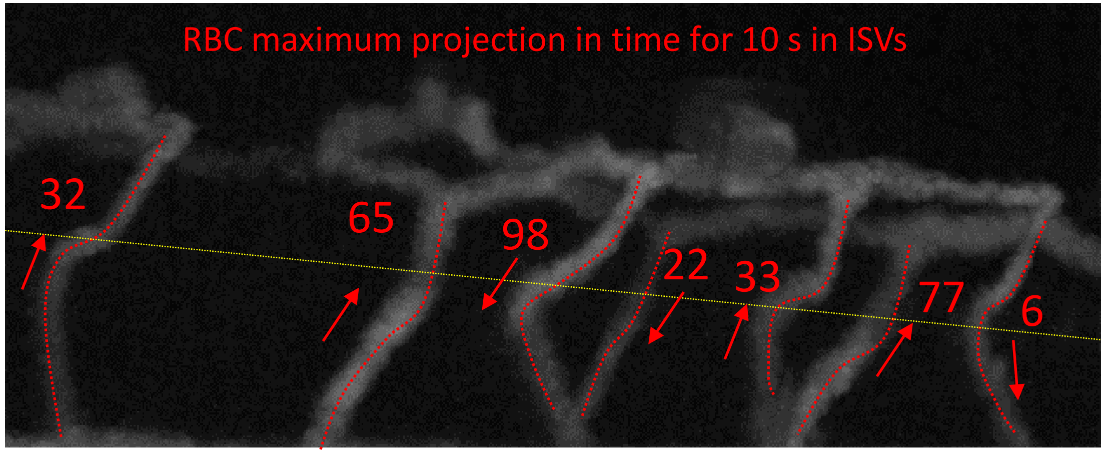


RBC count in ISVs for 10 s

Total: 333

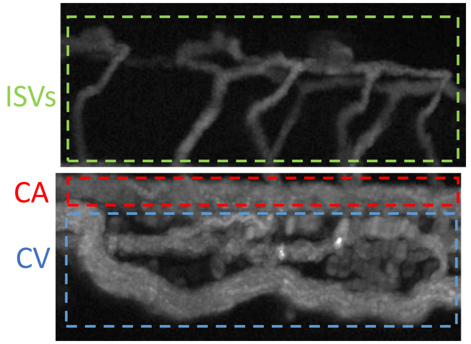

### ML1dblKO #12: HR = 138 bpm; Moderate perfusion

CA

23 beats in 10 seconds

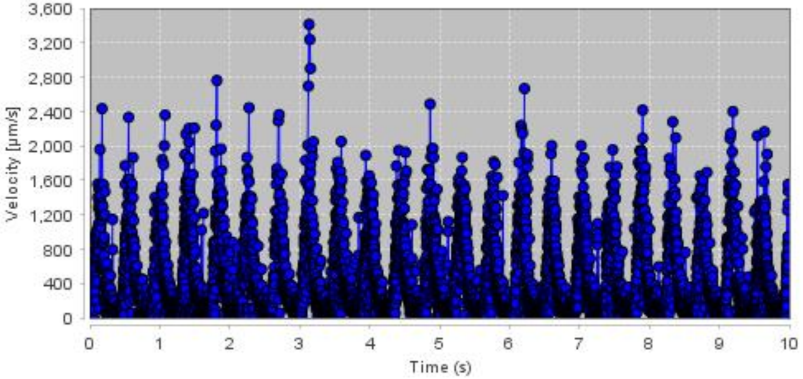


CV

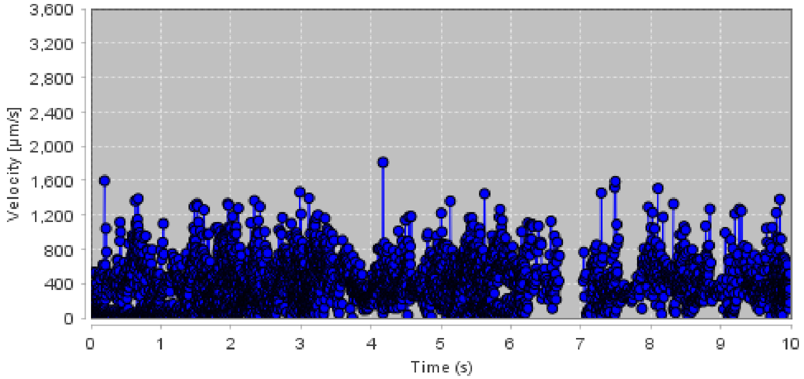


ISVs

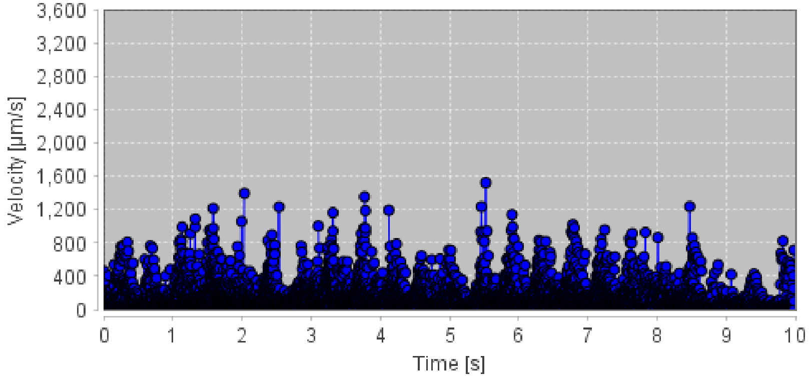


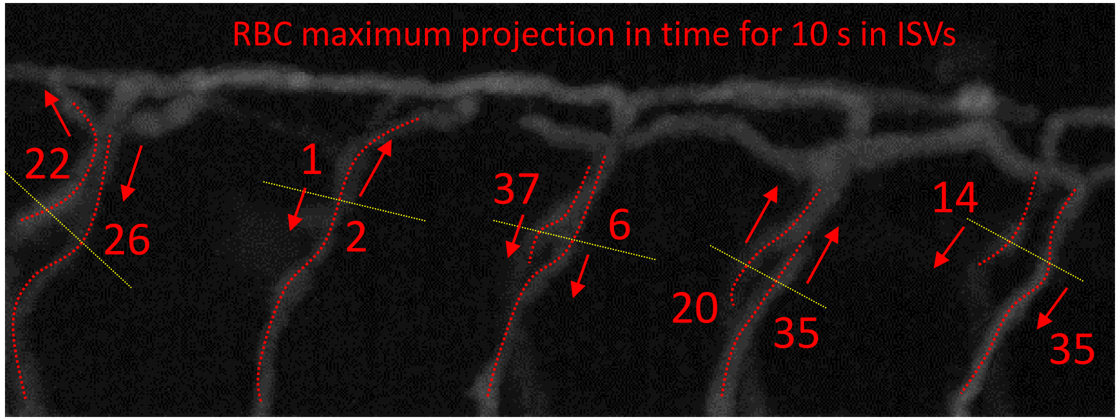


RBC count in ISVs for 10 s

Total: 198

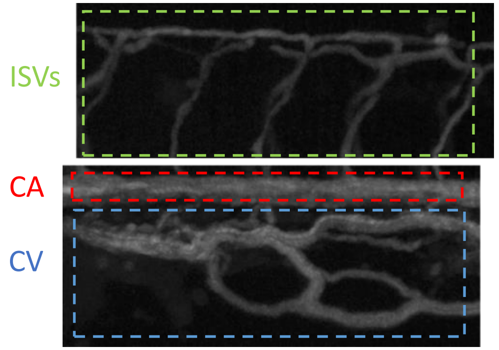

### ML1dblKO #3: HR = 162 bpm; Poor perfusion

CA 27 beats in 10 seconds

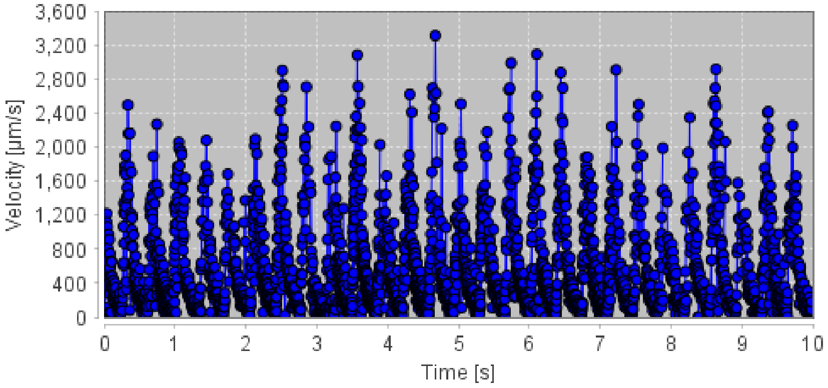


CV

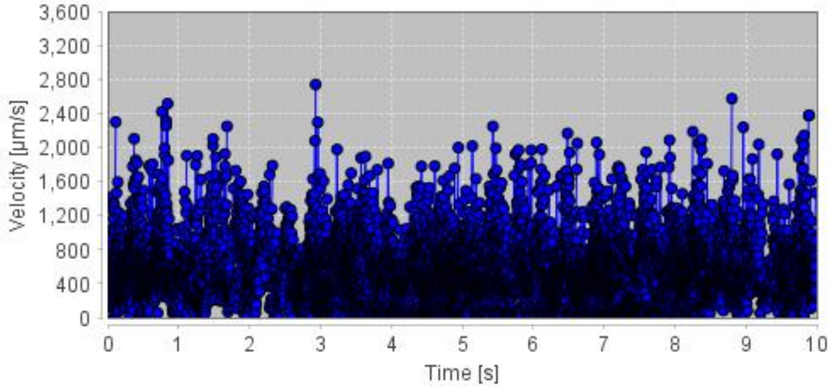


ISVs

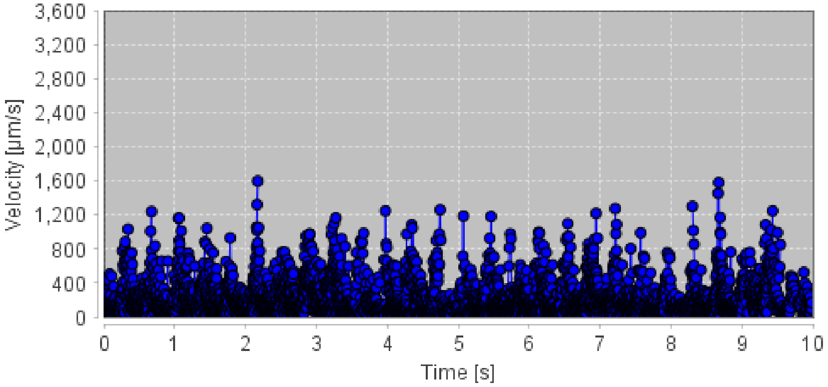


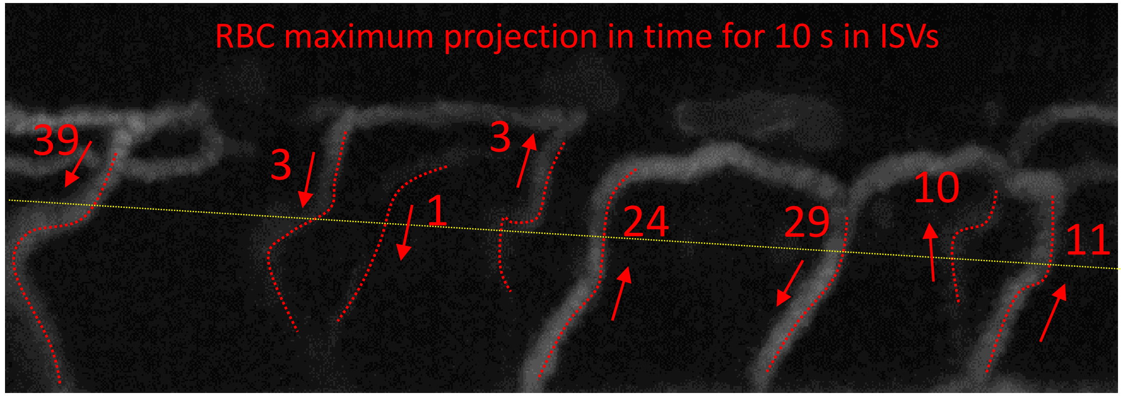


RBC count in ISVs for 10 s

Total: 120

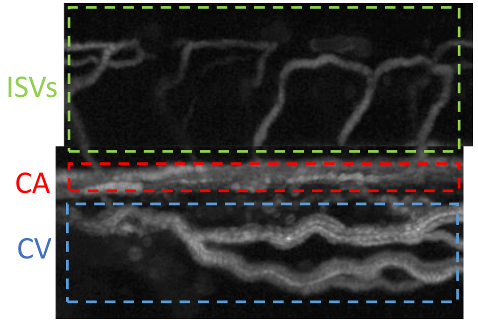

### ML1dblKO #7: HR = 140 bpm; Poor perfusion

14 beats in 6 seconds

CA

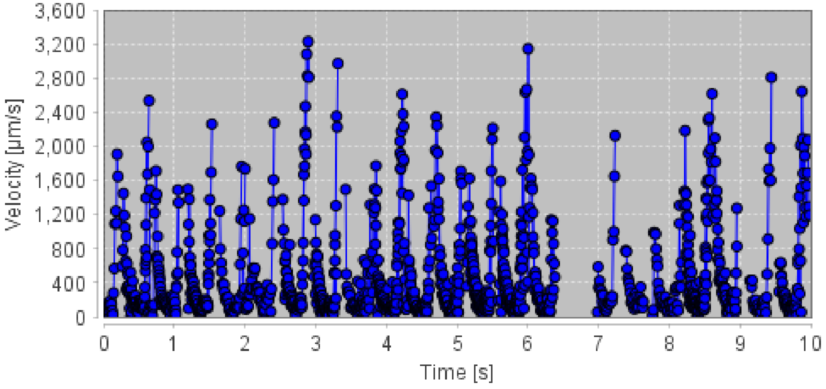


CV

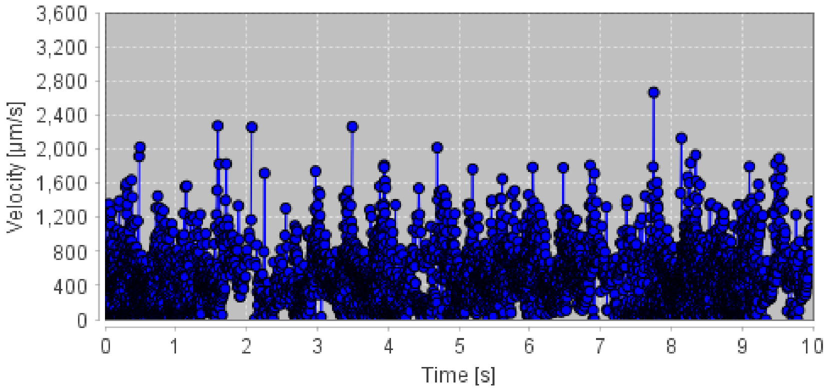


ISVs

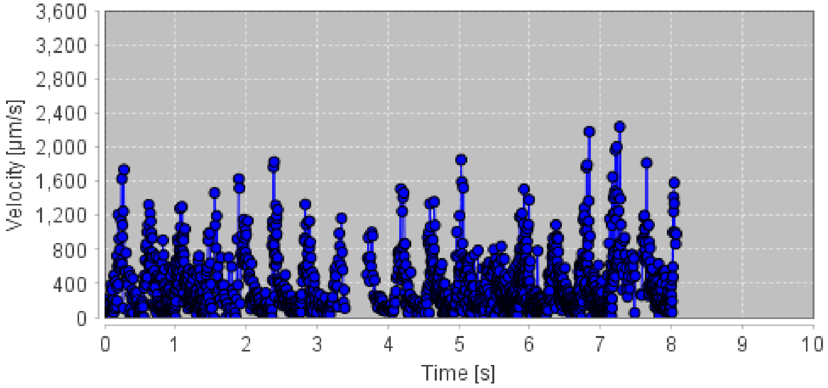


RBC maximum projection in time for 10 s in ISVs

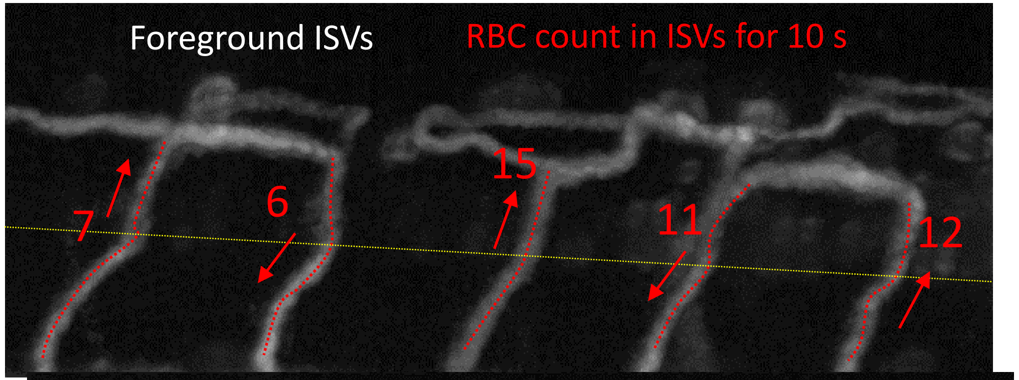


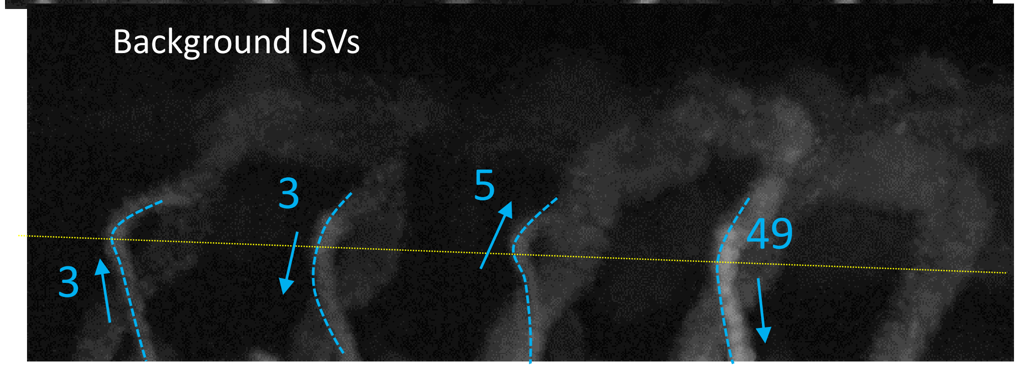


Total: 111

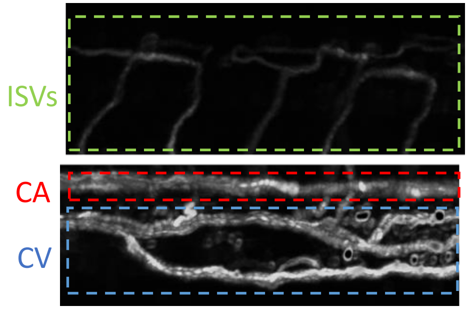

### ML1dblKO #9: HR = 140 bpm; Poor perfusion

CA

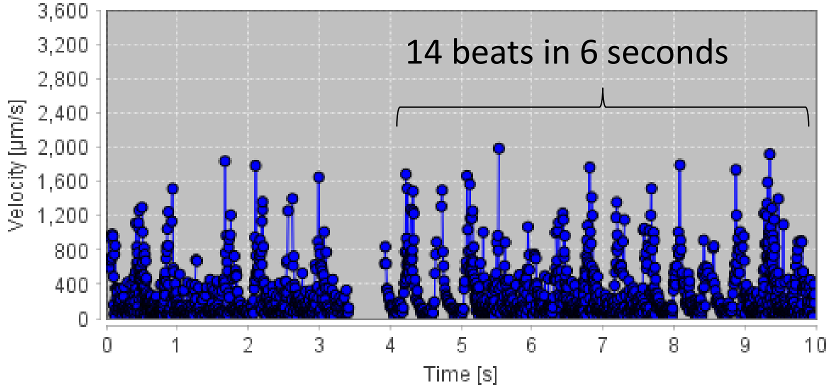


CV

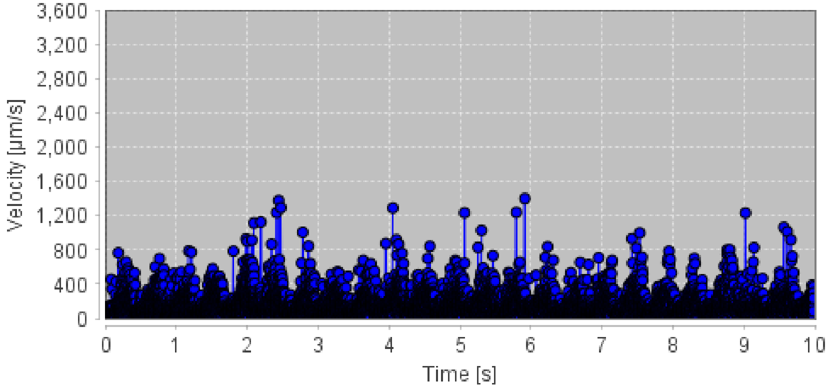


ISVs

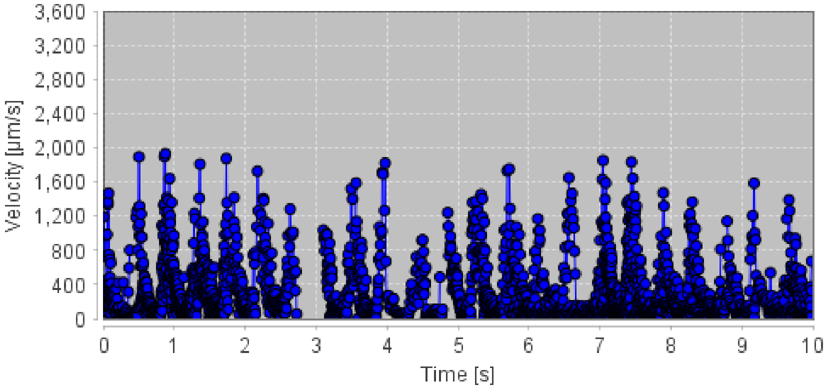


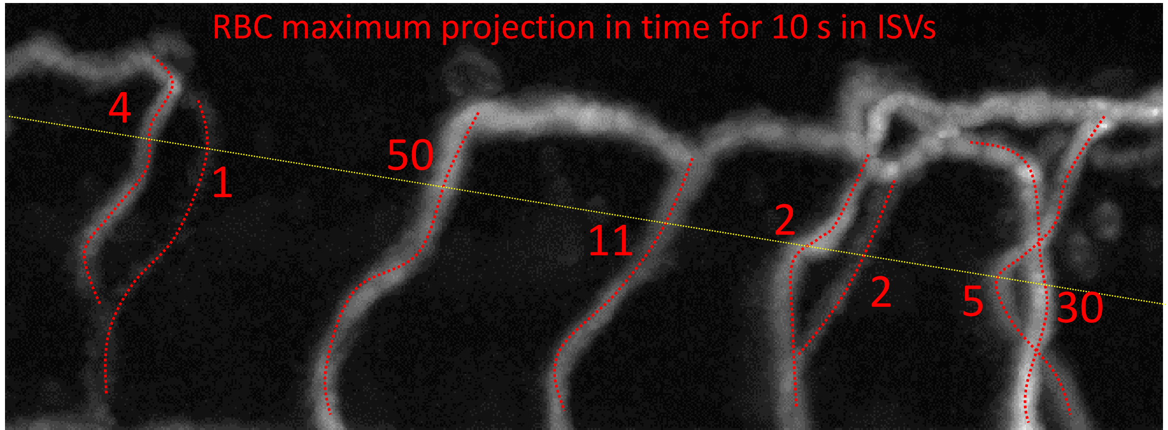


RBC count in ISVs for 10 s

Total: 105

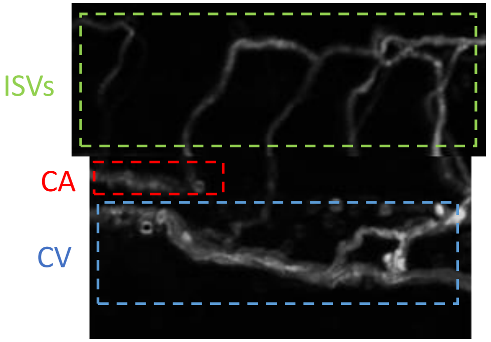

### Poor perfusion levels in ISVs

### ML1dblKO #5: HR = 140 bpm; Poor perfusion

CA

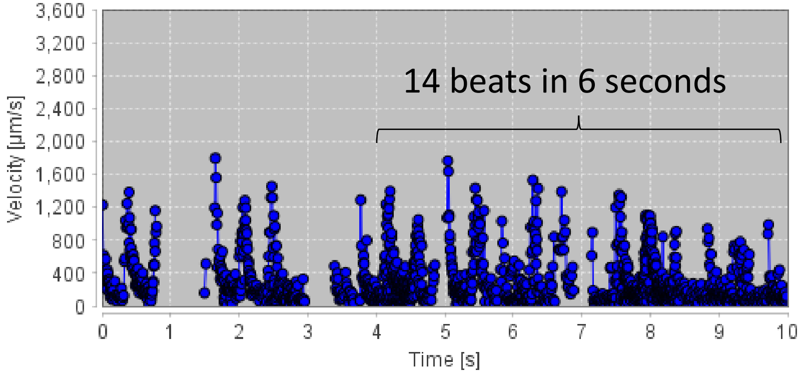


CV

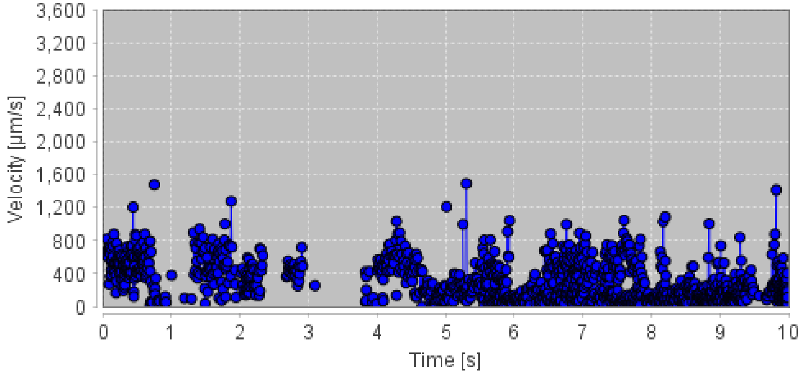


ISVs

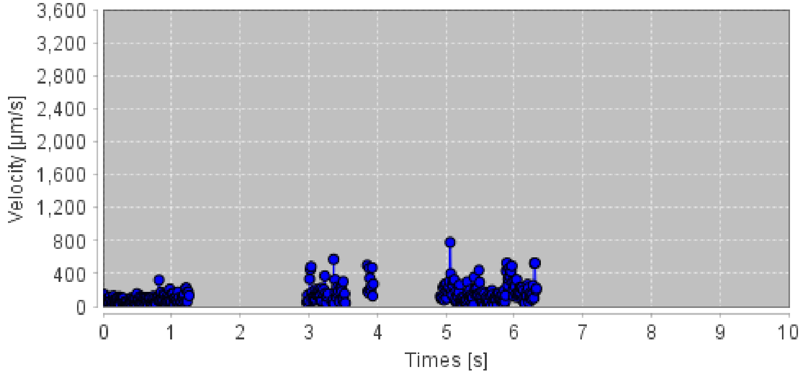


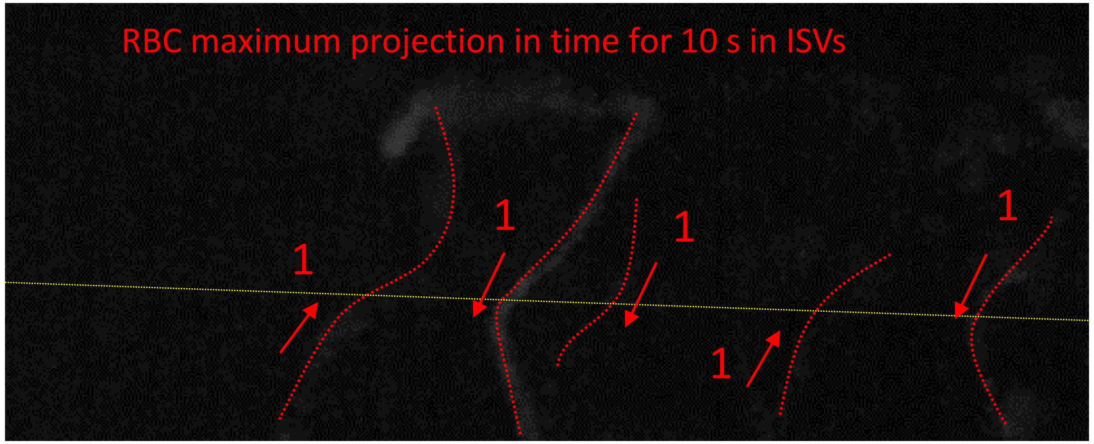


RBC count in ISVs for 10 s

Total: 5

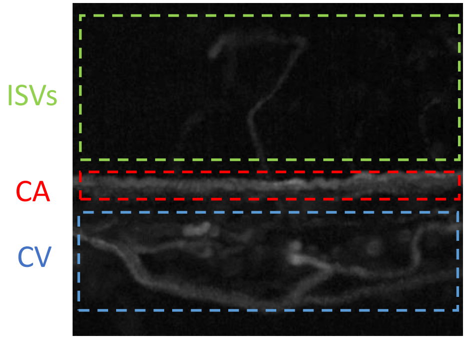

### ML1dblKO #8: HR = 126 bpm; Poor perfusion

CA

21 beats in 10 seconds

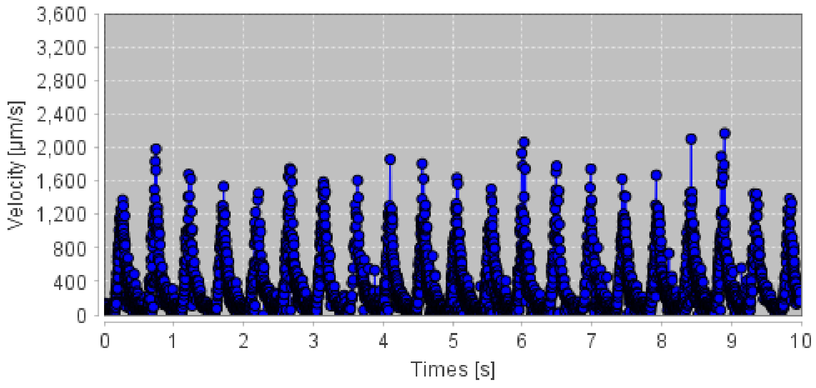


CV

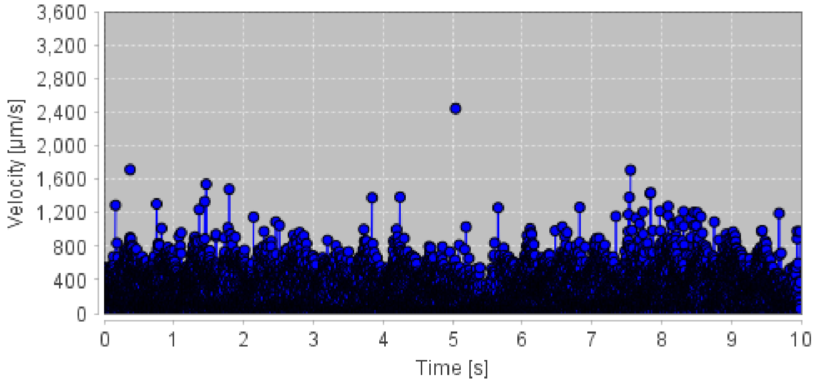


ISVs

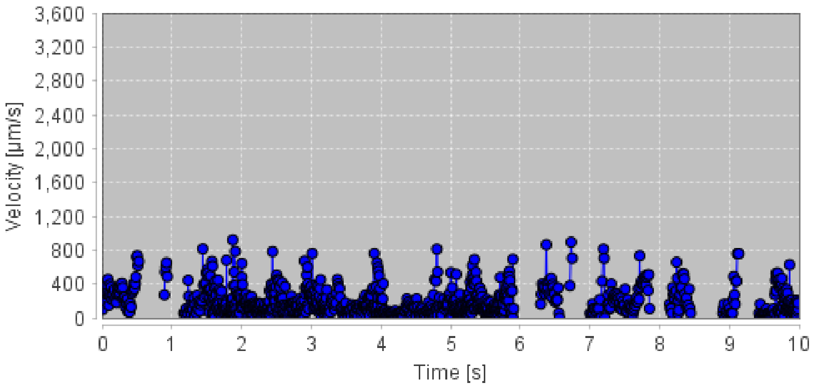


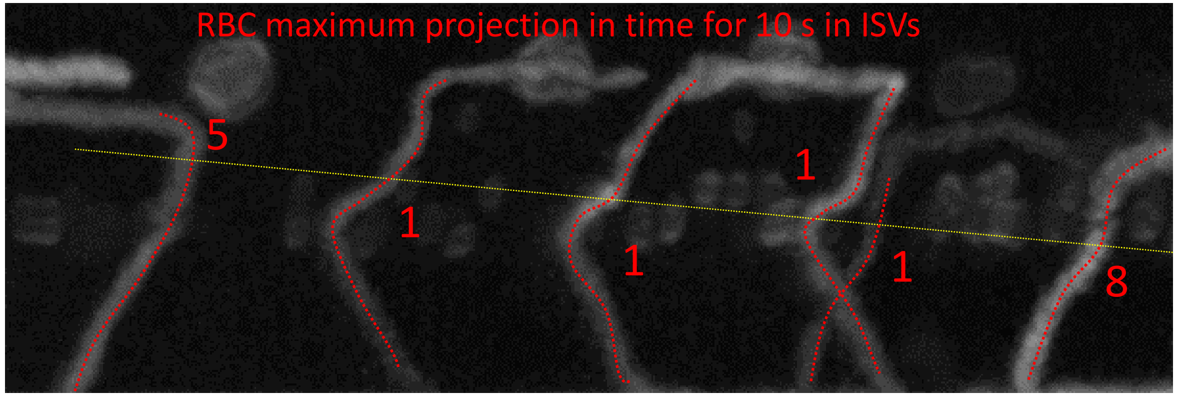


RBC count in ISVs for 10 s

Total: 16

ISVs

CA

CV

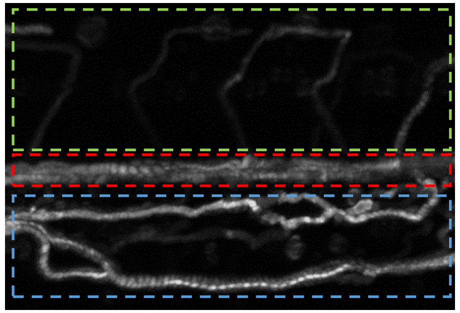

### ML1dblKO #14: HR = 120 bpm; Poor perfusion

CA

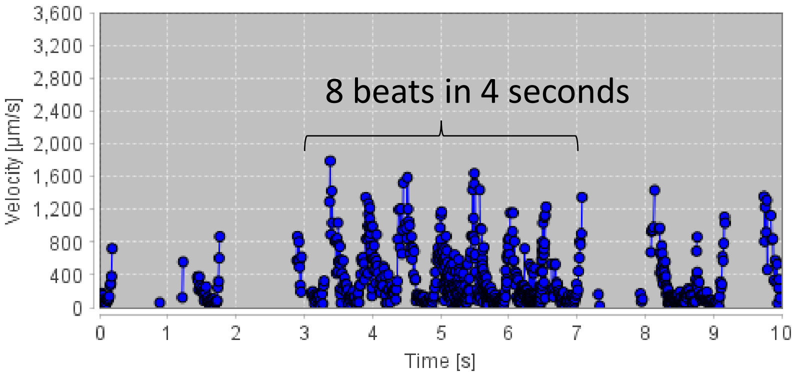


CV

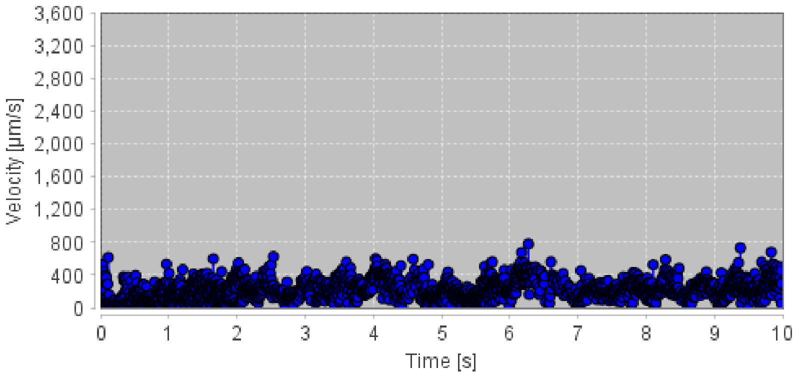


ISVs

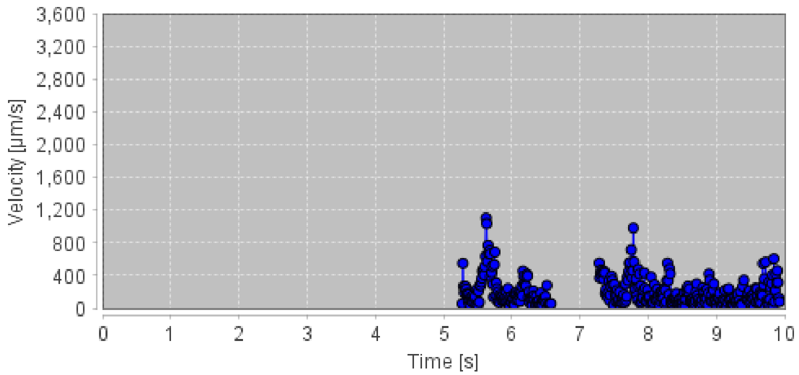


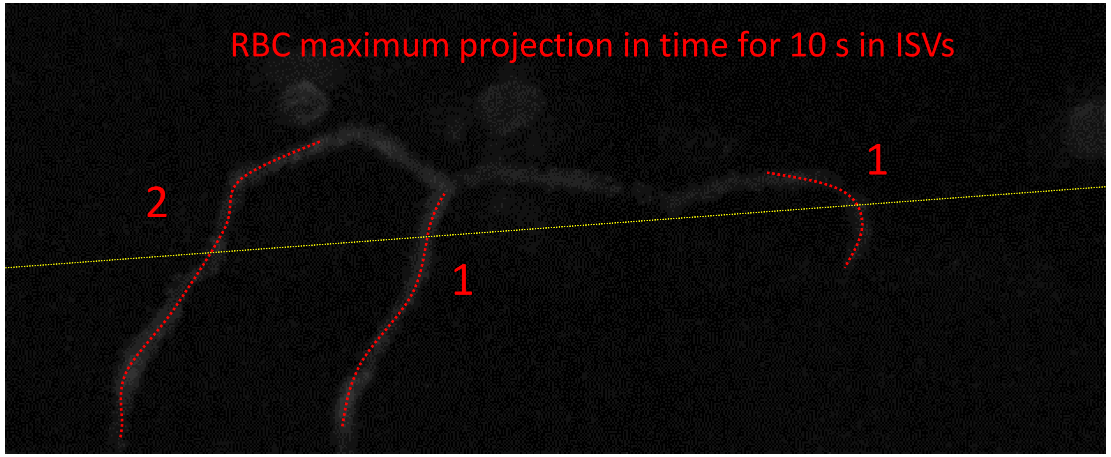


RBC count in ISVs for 10 s

Total: 4

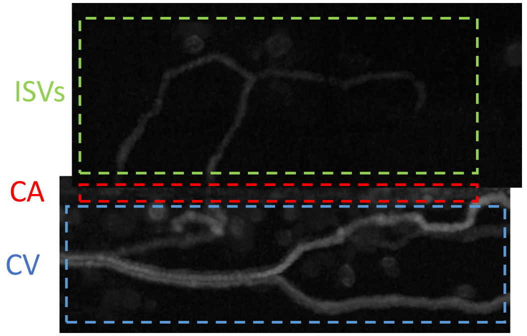

### Zero perfusion levels in ISVs

### ML1dblKO #4: HR = 150 bpm; No perfusion in ISVs

CA

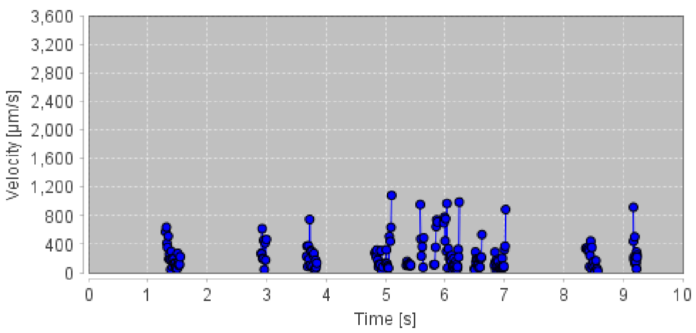


CV

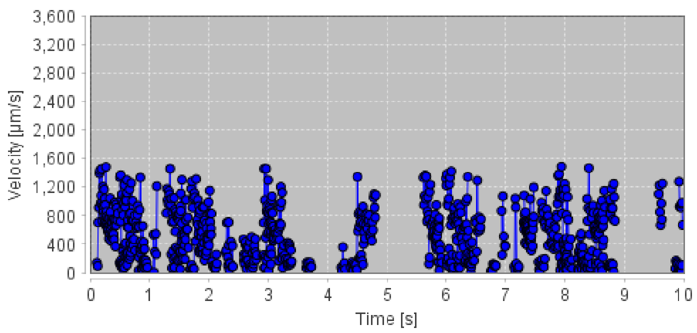


RBC tracking and velocity acquisition in ISVs not possible because RBCs are not entering the ISVs

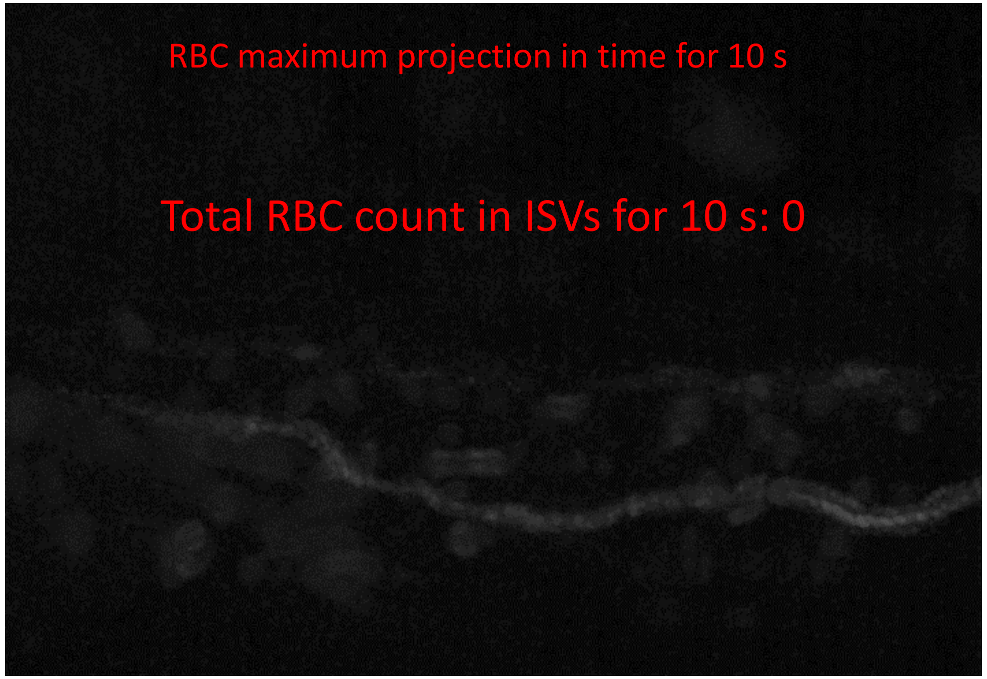


Heart beat determination

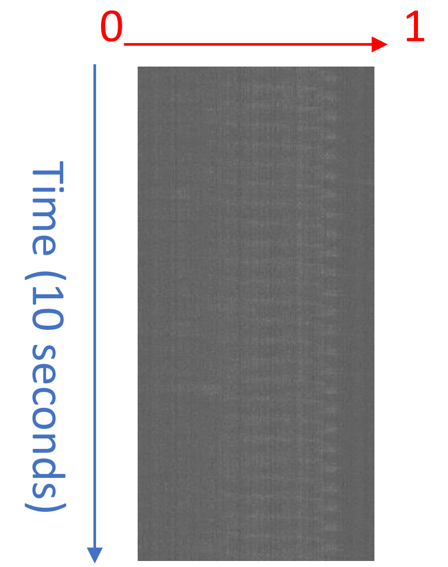


Kymograph along red arrow line

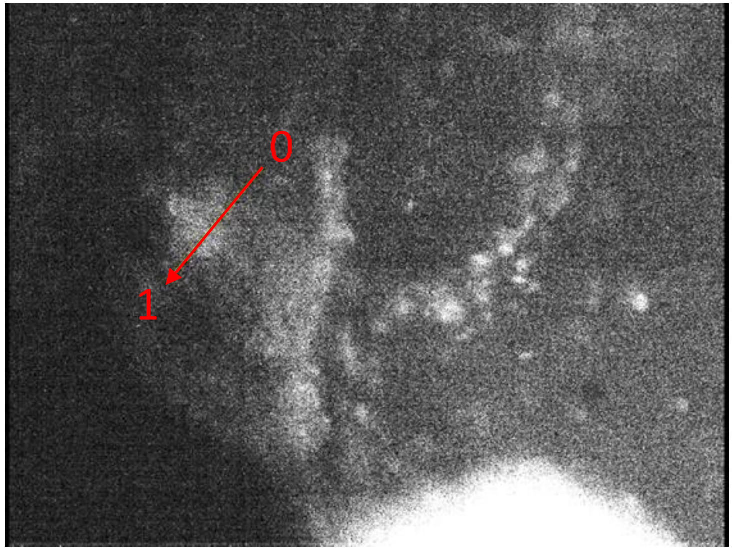


25 beats in 10 seconds

#### ML1dblKO #11: HR = 144 bpm; No perfusion in entire posterior trunk

Heart beat determination

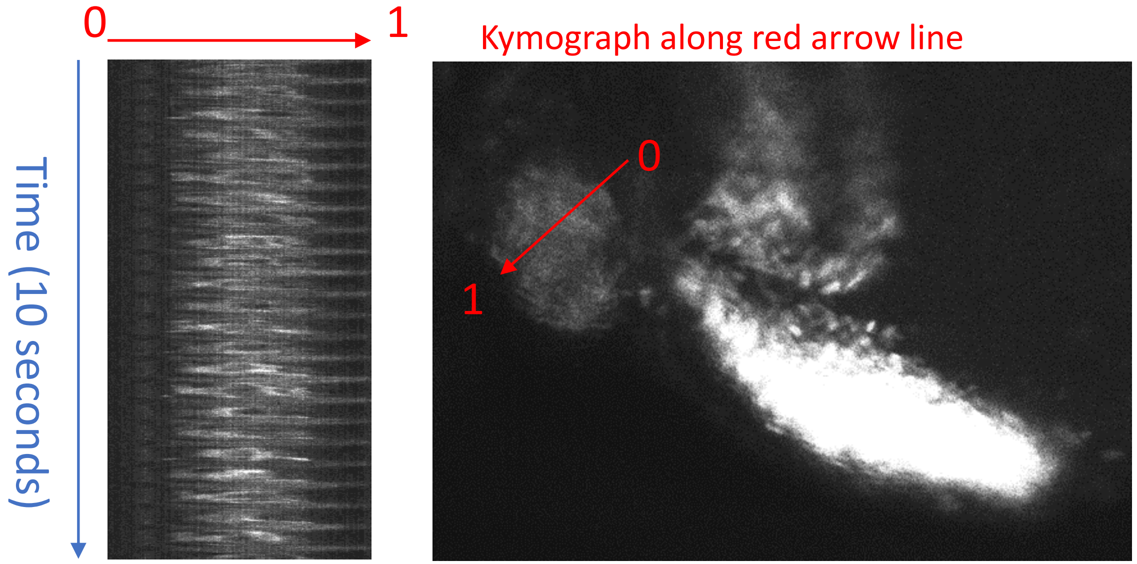


24 beats in 10 seconds

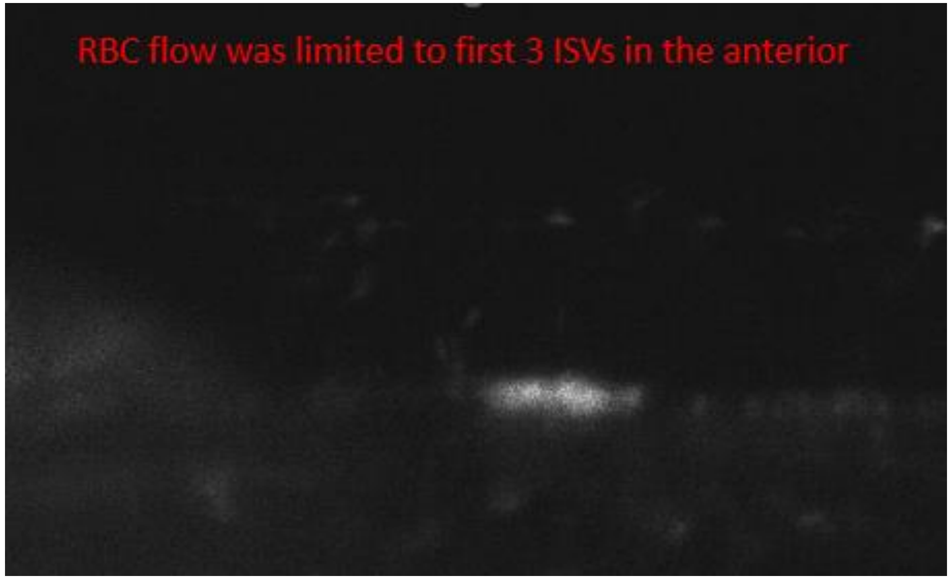


RBC tracking and velocity acquisition for entire trunk network is not possible because RBCs are not flowing

Total RBC count in ISVs for 10 s: 0

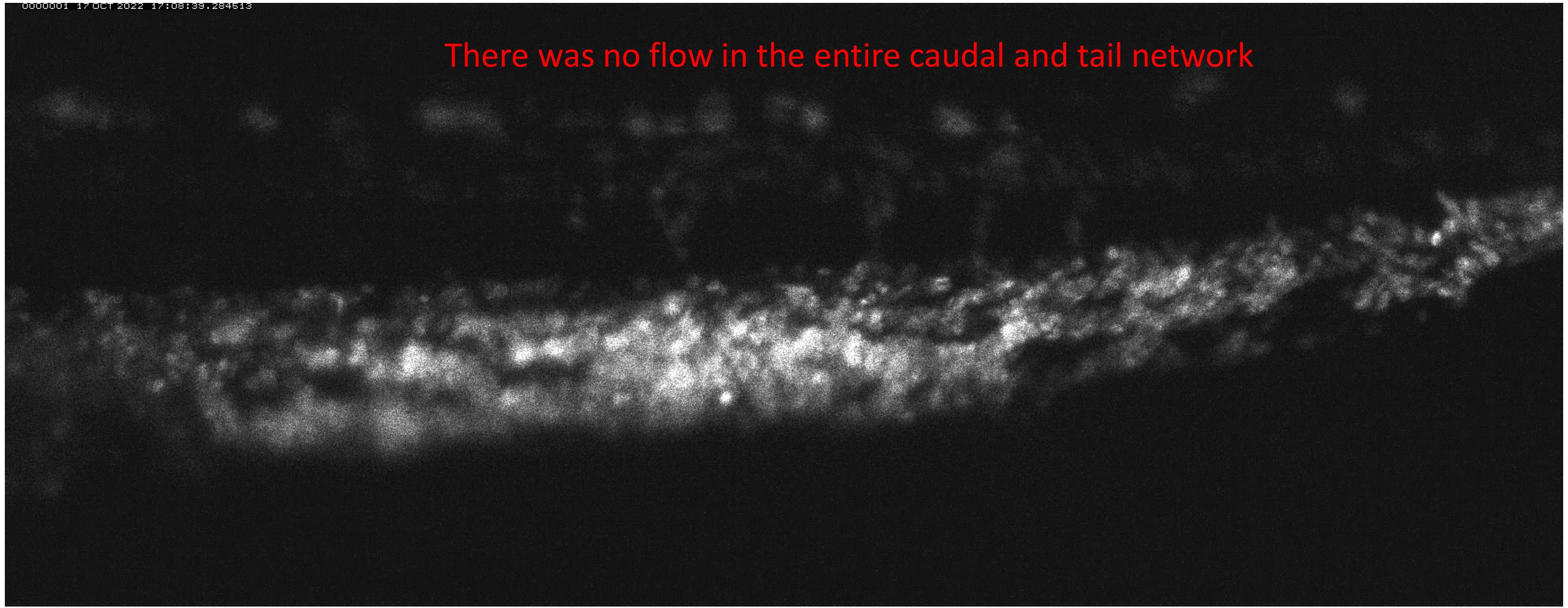
